# RETROFIT: Reference-free deconvolution of cell-type mixtures in spatial transcriptomics

**DOI:** 10.1101/2023.06.07.544126

**Authors:** Roopali Singh, Xi He, Xinyue Wang, Adam Keebum Park, Ross Cameron Hardison, Xiang Zhu, Qunhua Li

## Abstract

Spatial transcriptomics (ST) enables genome-wide measurement of gene expression in intact tissues, but typically captures mixtures of multiple cell types at each spatial location. Deconvolving these mixtures is essential for resolving cell-type-specific spatial organization and transcriptional programs. Existing approaches often rely on matched single-cell references or curated marker genes, which may be unavailable, incomplete, or difficult to integrate across platforms. We present RETROFIT, a Bayesian framework for reference-free deconvolution of spatial transcriptomics data that operates directly on sequencing measurements and incorporates external information only at a post hoc annotation stage when available. Across extensive simulations and multiple real datasets, RETROFIT demonstrates robust performance, outperforming existing reference-free methods and matching or exceeding reference-based approaches when references are imperfect. Notably, RETROFIT remains effective at near–single-cell resolution, as demonstrated on Visium HD data, recovering fine-grained spatial patterns without requiring single-cell references or marker genes. These results establish RETROFIT as a broadly applicable approach for reference-free spatial transcriptomics analysis across platforms and resolutions. RETROFIT is available at https://bioconductor.org/packages/retrofit/.

## Introduction

Spatial organization is fundamental to tissue development, homeostasis, and disease. Recent advances in spatial transcriptomics (ST) technologies^1,2^ have enabled genome-wide measurements of gene expression directly within intact tissue sections ^3^, providing new opportunities to study how cellular composition and transcriptional programs are organized in space^4^. These technologies have been widely applied to complex tissues such as brain^5^, intestine^6^, and tumor microenvironments ^7^, revealing spatially structured molecular patterns that are difficult to resolve using dissociative single-cell assays.

Most sequencing-based ST platforms ^1^ measure gene expression at discrete spatial locations (“spots”) arranged on a two-dimensional tissue section. Depending on platform resolution, each spot may capture transcripts from multiple cells. For example, Visium, which profiles spots approximately 55*µm* in diameter^8,9^, encompass 6-10 cells when applied to human intestinal samples^10^, whereas Slide-seq and related platforms approach cellular resolution^11^. Emerging technologies such as Visium HD further increase spatial resolution by densely sampling subcellular-scale regions. Across platforms, however, ST measurements typically reflect mixtures of transcripts from multiple cell types, motivating the development of computational deconvolution methods to infer cell-type composition and transcriptional profiles within each spatial location.

A wide range of computational methods has been developed for cell-type deconvolution in spatial transcriptomics^12^, including supervised approaches that leverage matched single-cell RNA-seq references^13–22^, as well as reference-free methods that incorporate curated marker genes or histological images to guide inference^17,23–27^ and fully reference-free methods that perform deconvolution using sequencing measurements alone^28^. While these strategies can be highly effective when high-quality auxiliary information is available, they may be limited in settings where suitable references, marker sets, or well-aligned histology are unavailable or difficult to obtain. Moreover, for emerging high-resolution spatial transcriptomics platforms, matched histological images are not always available or directly aligned with the profiled tissue section, which can further constrain the applicability of image-integrative approaches in some experimental settings. Over the course of the field’s development, most deconvolution methods have relied on such auxiliary information, most commonly matched single-cell references, and increasingly marker sets or image-derived features, to improve identifiability and interpretability. In contrast, methods that perform deconvolution using sequencing measurements alone remain a smaller subset of available approaches ^28^, despite their practical value in studies where external data are missing, incomplete, or challenging to integrate.

Here, we introduce RETROFIT, an unsupervised Bayesian framework for reference-free cell-type deconvolution of spatial transcriptomics data. Built on a Bayesian hierarchical model, RETROFIT factorizes the spatial transcriptomics count matrix into two interpretable components: one capturing the transcriptional profiles of latent cellular components and the other representing their relative contributions across spatial locations. Deconvolution is performed using sequencing measurements alone, while external information, such as marker genes or single-cell references, can be incorporated post hoc through simple annotation strategies when available. This separation of data decomposition from biological annotation enhances robustness to variability in reference quality and availability, while preserving interpretability and reusability of inferred components.

Across simulated and real datasets, RETROFIT demonstrates strong and consistent performance across varying spot sizes, tissue heterogeneity, and reference quality, with particularly robust behavior in settings with incomplete references and in ultra–high-resolution spatial transcriptomics data. It recovers known cellular organization in mouse cerebellum Slide-seq data, reveals spatiotemporal patterns of cellular composition and transcriptional specificity in human intestinal development, and remains effective at near–single-cell resolution on Visium HD data—without relying on curated marker genes, matched single-cell references, or histological images. Together, these results highlight RETROFIT’s flexibility and broad applicability across platforms, spatial resolutions, and data-availability settings.

## Results

### RETROFIT deconvolves ST data independent of single-cell gene expression references

RETROFIT is a reference-free approach for cell-type deconvolution of ST data (Fig. 1). It takes an ST count matrix **X**, which consists of *G* genes across *S* spots, as its sole input, and performs an unsupervised projection of **X** onto a lower-dimensional space spanned by *L* non-negative latent components, independent of any external reference. The number of factors, *L*, is typically chosen to be greater than the actual number of cell types (*K*) present in the ST sample, allowing RETROFIT to capture all the relevant cellular components in each spot. The expression of each gene at each spot for each component is further decomposed into gene-specific expression and background expression shared by all genes. The *L* latent components, mined solely from ST data, often contain information that distinguishes cell types with distinct transcriptomic profiles, forming the basis for cell-type deconvolution.

**Figure 1.**
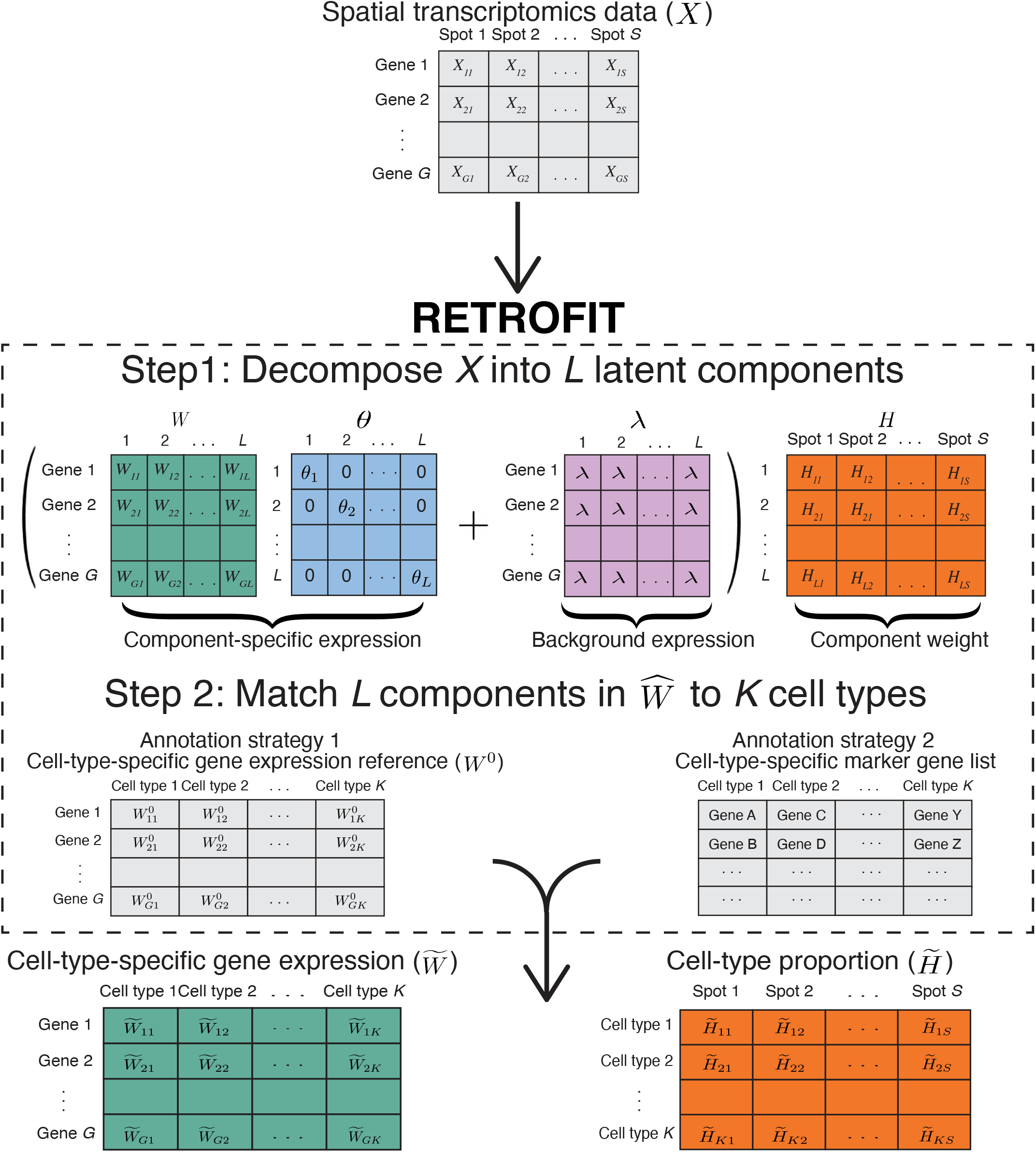
Overview of RETROFIT. Step 1: RETROFIT takes a ST data matrix as the only input and decomposes this matrix into latent components in an unsupervised manner (Algorithm 1). Step 2: RETROFIT matches these latent components to known cell types using either a cell-type-specific gene expression reference (Algorithm 2) or a list of cell-type-specific marker genes (Algorithm 3) for the cell types present in the ST sample, and outputs a cell-type-specific gene expression matrix and a cell-type proportion matrix.

RETROFIT is formulated as a Bayesian hierarchical model with a Poisson likelihood for the observed ST data and Gamma priors for the unknown parameters (Methods). It deconvolves the ST data matrix into two matrices: one representing component-specific gene expression and the other representing the proportion of each component. To enable the analysis of large-scale ST data, RETROFIT employs a structured stochastic variational inference (SSVI) algorithm ^29^, which scales efficiently with thousands of genes and spots (Algorithm 1; Supplementary Table 1). The software is available as a Bioconductor R package at https://bioconductor.org/packages/retrofit/.

Like any unsupervised learning method, RETROFIT produces unlabeled results. To assign known cell types to the latent components inferred by RETROFIT, we develop two simple post hoc cell-type annotation strategies. The first strategy requires a cell-type-annotated gene expression reference (**W**^0^) for all *K* cell types present in the ST data, a standard assumption in most ST deconvolution methods^18,30^. This reference can be derived from external single-cell transcriptomics data that match the tissue type of the ST data. With this reference, we can calculate correlations between the component-specific expression profiles estimated by RETROFIT and the observed cell-type-specific expression profiles in the reference. The cell type with the highest correlation for a component is considered the most probable annotation (Algorithm 2). The second strategy does not require any gene expression references, but instead requires a curated list of cell-type-specific marker genes for all *K* cell types present in the ST data. This approach complements the first strategy when a proper cell-type-specific expression reference is unavailable. Using the marker gene list, we calculate a marker expression score for each component in each cell type, defined as the sum of normalized component-specific expression levels of marker genes in that cell type. We then annotate each component by the cell type with the highest marker expression score (Algorithm 3). Once the latent components are matched to cell types by either strategy, RETROFIT outputs a cell-type-specific expression matrix for all genes 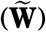 and a cell-type proportion matrix for all spots 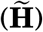 as the final results.

### RETROFIT surpasses existing methods across a wide variety of synthetic ST data

We benchmarked RETROFIT against existing methods through extensive simulations based on a scRNA-seq dataset from the mouse cerebellum^11^. To mimic sequence-based ST data from different platforms and samples, we generated datasets with varying levels of spot size and cell-type heterogeneity by altering the number of cells per spot (*N*) and the maximum number of cell types per spot (*M*), respectively, and we also introduced variation in sequencing depth across spots within the same ST slide (Algorithm 4). In addition, to examine the dependence of reference-based deconvolutions on the quality of single-cell transcriptomic data, we created single-cell references with varying degrees of correspondence to the cell types present in the target ST data. On each simulated ST dataset, we compared RETROFIT with three reference-based methods that often achieved state-of-the-art performance in recent benchmarking studies ^18–21^: Stereoscope ^13^, RCTD^15^ and Cell2location^16^, as well as two reference-free methods: STdeconvolve^28^, which performs unsupervised topic modeling on ST counts, and CARDfree ^17^, which incorporates marker-gene information and spatial smoothing to guide deconvolution. We evaluated each method in two aspects: (1) explanatory power measured by the root-mean-square error (RMSE; Fig. 2a) and correlation (Fig. 2b) between the true and estimated cell-type proportions at each spot; (2) predictive power measured by the normalized RMSE (NRMSE; Fig. 2c; Supplementary Fig. 1) and correlation (Fig. 2d) between the observed and reconstructed gene expression profiles at each spot, where the reconstructed expression profiles were sums of the single-cell expression profiles in individual cell types weighted by the estimated cell-type proportions. Details of simulation and evaluation are provided in Methods.

**Figure 2.**
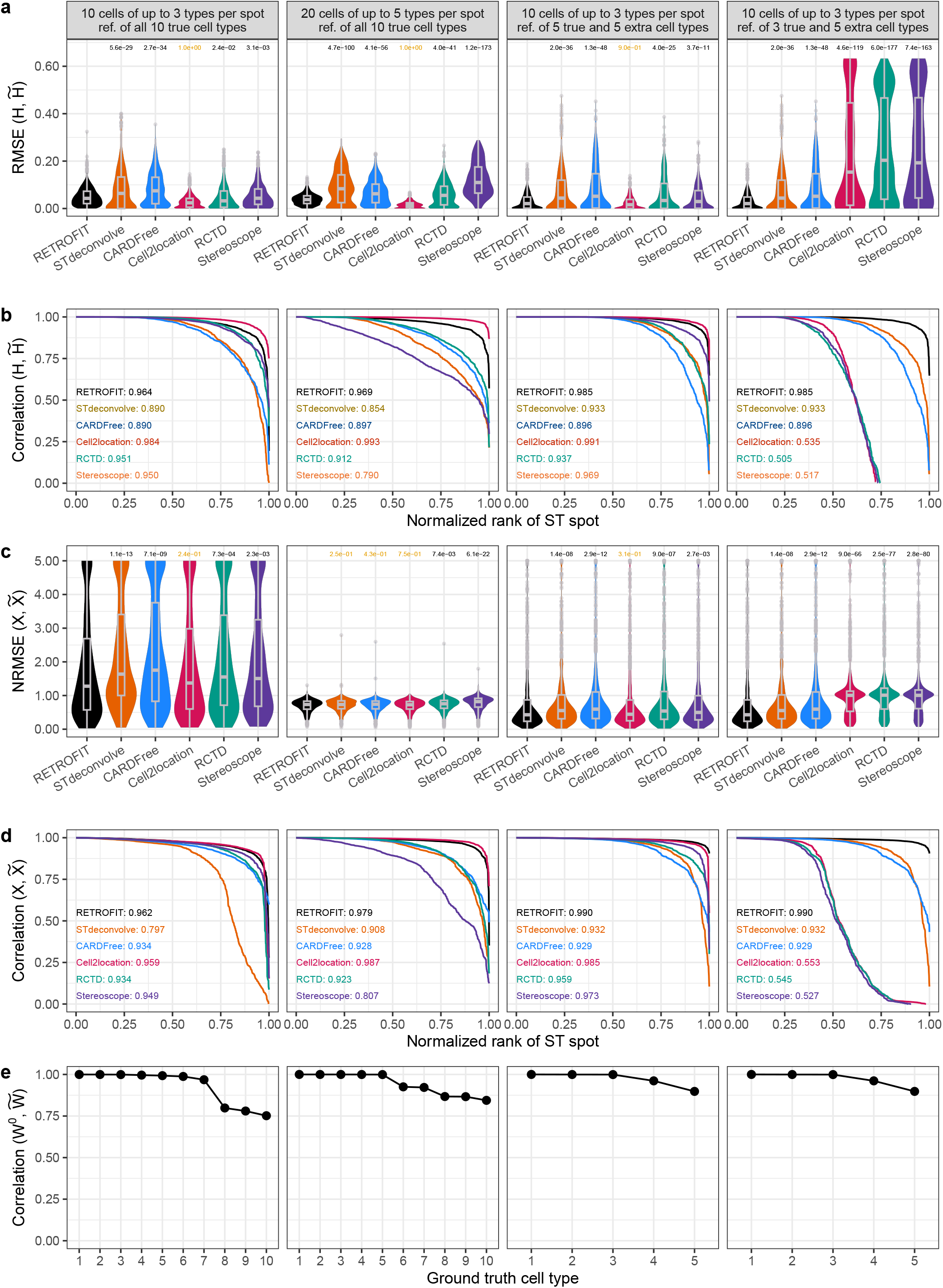
Evaluating RETROFIT on synthetic ST data with different spot size, cell-type complexity and reference quality. Column 1: small spots (*N* = 10 cells per spot) with low cell-type complexity (up to *M* = 3 cell types per spot from *K* = 10 cell types). Column 2: large spots (*N* = 20) with high cell-type complexity (*M* = 5 and *K* = 10). Columns 3-4: *N* = 10, *M* = 3 and *K* = 5. All synthetic data have 1000 spots with 500 genes. Reference-based methods were provided with the following single-cell transcriptomic references. Columns 1-2: exact reference of all 10 ground-truth cell types. Column 3: all 5 ground-truth plus 5 irrelevant cell types. Column 4: only 3 out of 5 ground truth plus 5 irrelevant cell types. **a** Distribution of RMSE and **b** ranked correlation between true (**H**) and estimated cell-type proportions 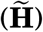 across all cell types at each spot. **c** Distribution of NRMSE and **d** ranked correlation between observed (**X**) and reconstructed expression 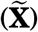 across all genes at each spot. The one-sided KS *P*-values are shown in **a** and **c** (black: *P* < 0.05; brown: *P* > 0.05). A small *P*-value indicates that RETROFIT estimates have stochastically lower RMSEs compared to another method. Box plots in **a** and **c** show medians (center lines), 25th–75th percentiles (box), and whiskers extending to the most extreme values within 1.5 × IQR; values beyond are outliers. The AUC of ranked correlations is shown for each method with a matching color in **b** and **d**. The NRMSEs are clamped at 5 in **c** to aid visualization and a non-clamped version is provided in the Supplementary Fig. 1.e Ranked correlation between the single-cell observation (**W**^0^) and RETROFIT estimation 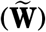 of cell-type-specific expression across all genes for each cell type.

We started with an ideal use case for reference-based methods, providing them with an exact reference of the single-cell expression profiles for all 10 cell types used to simulate the ST data. In contrast, reference-free methods would benefit little from such an exact reference, as they decompose the ST data independent of external references. The first two columns of Fig. 2 show the simulation results for this scenario in two settings: (1) smaller spot size and lower cell-type heterogeneity: *N* = 10 cells and up to *M* = 3 cell types per spot; and (2) larger spot size and higher cell-type heterogeneity: *N* = 20 cells and up to *M* = 5 cell types per spot. In both settings, we simulated ST data for *G* = 500 genes and *S* = 1000 spots with *K* = 10 cell types. Although these simulations were designed to favor reference-based methods, RETROFIT performed comparably to the best-performing reference-based method (Cell2location) and significantly outperformed the two reference-free methods (STdeconvolve and CARDfree).

We next considered a more realistic use case involving imperfect single-cell transcriptomic references that either included irrelevant cell types or excluded relevant cell types. We evaluated how these imperfections impacted reference-based and reference-free deconvolutions of ST data. Here we simulated ST data for *G* = 500 genes and *S* = 1000 spots, with *N* = 10 cells from up to *M* = 3 out of *K* = 5 cell types per spot, using the same data and scheme as before (Methods). We then created two imperfect single-cell expression references: (1) the complete set of the 5 ground truth cell types used to generate ST data plus 5 extra cell types; and (2) 3 out of the 5 ground truth cell types plus 5 extra cell types. All methods were evaluated on the same ST data using the 5 ground truth cell types for both scenarios. For the second scenario, we also evaluated all methods using only the 3 ground truth cell types present in the reference and arrived at the same conclusion (Supplementary Fig. 2).

For all reference-based methods, we observed a heavy reliance on the completeness of relevant cell types in the single-cell reference. These methods maintained robustness to irrelevant cell types when the reference included five additional cell types alongside the five ground truth cell types (Fig. 2, column 3). However, their performance markedly declined when two of the five ground truth cell types were missing from the reference (Fig. 2, column 4), highlighting a major limitation of reference-based deconvolutions.

In contrast, RETROFIT consistently demonstrated optimal performance regardless of reference quality. When two of the five ground-truth cell types were missing from the single-cell reference, RETROFIT showed substantial gains in accuracy over all reference-based methods for both cell-type proportion estimation (KS test *P* ≤ 4.6 × 10^−119^; Fig. 2a) and gene expression reconstruction (KS test *P* ≤ 9.0 × 10^−66^; Fig. 2c). In nearly all spots (≥ 97.4%), the estimated cell-type proportions (AUC= 0.985) and reconstructed gene expression profiles (AUC=0.990) from RETROFIT were strongly correlated with the ground truth (Pearson *R* ≥ 0.9), whereas only less than 49.9% and 41.5% of spots achieved the same level of correlation for estimated proportions (AUC = 0.505 − 0.535; Fig. 2b) and reconstructed expression profiles (AUC = 0.527 − 0.553; Fig. 2d) from reference-based methods, respectively.

Like RETROFIT, STdeconvolve and CARDfree also showed robustness against the incompleteness of cell-type in the single-cell reference (Fig. 2, column 4), as all three methods deconvolve ST data independently of single-cell transcriptomic references. However, STdeconvolve and CARDfree performed worse than RETROFIT in cell-type proportion estimation, as indicated by significantly larger RMSE values (KS test *P* = 2.0 ×10^−36^ for STdeconvolve and *P* = 1.3 ×10^−48^ for CARDfree; Fig. 2a) and smaller AUC scores (0.985 for RETROFIT versus 0.933 for STdeconvolve and 0.896 for CARDfree; Fig. 2b). Additionally, in gene expression reconstruction, Stdeconvolve and CARDfree had significantly larger NRMSE values (KS test *P* = 1.4 × 10^−8^ for STdeconvolve and *P* = 2.9 × 10^−12^ for CARDfree; Fig. 2c) and smaller AUC scores (0.990 for RETROFIT versus 0.932 for STdeconvolve and 0.929 for CARDfree; Fig. 2d).

Using these synthetic ST data, we also evaluated the correlation between cell-type-specific gene expression profiles estimated by RETROFIT and observed single-cell expression profiles for each cell type (Fig. 2e). Across all simulations, RETROFIT estimates were highly correlated with the single-cell data for all cell types (Pearson *R* > 0.75 when *N* = 10, *M* = 3 and *K* = 10; *R* > 0.84 when *N* = 20, *M* = 5 and *K* = 10; *R* > 0.89 when *N* = 10, *M* = 3 and *K* = 5), confirming that the reference-free estimation in RETROFIT effectively captures cell-type-specific transcriptional characteristics.

Lastly, we performed a wide range of secondary simulations and comparisons to further benchmark RETROFIT against existing methods. Using the same mouse cerebellum scRNA-seq data ^11^, we simulated ST slides with small spot size (Supplementary Fig. 3) or uneven cell-type prevalence (Supplementary Fig. 4), and compared the performance of RETROFIT with that of existing methods. Additionally, we evaluated RETROFIT’s performance in simulations based on MERFISH data from the mouse hypothalamus^31^, which preserve the spatial patterns of transcriptomic profiles found in real tissues (Supplementary Fig. 5). We further utilized the five subtypes of inhibitory cells in the same MERFISH data ^31^ to simulate ST data from highly similar cellular contexts and assess the robustness of RETROFIT in this scenario (Supplementary Fig. 6). Despite the diverse nature of these secondary analyses, all results consistently highlight the competitive and robust performance of RETROFIT, aligning with the conclusions drawn from the primary simulations (Fig. 2).

Taken together, the simulation results consistently show RETROFIT’s performance gain over existing reference-free deconvolution methods and its competitive standing compared to the best-performing reference-based deconvolution approach, even without exploiting an exact single-cell expression reference. These results underscore the particular advantage of RETROFIT over reference-based methods when key cell types relevant to the ST data are absent from the single-cell transcriptomic reference.

### RETROFIT outperforms existing methods to deconvolve mouse cerebellum Slide-seq data

We compared RETROFIT with existing approaches using a mouse cerebellum Slide-seq dataset ^11^ of 17919 genes at 27261 spots, a dataset widely used for benchmarking ST deconvolution methods. For the three reference-based methods – RCTD, Stereoscope, and Cell2location – a scRNA-seq reference for 10 mouse cerebellar cell types from the same study^11^ was provided. In contrast, RETROFIT and two other reference-free methods, STdeconvolve and CARDfree, did not use this single-cell reference for deconvolution; instead, they used the scRNA-seq dataset solely for post hoc matching of latent components to known cell types. Details of applying each method to the Slide-seq study are available in Methods.

To benchmark the deconvolution methods on the Slide-seq dataset, we focused on three cerebellum cell types: granule cells, oligodendrocytes, and Purkinje cells, for which curated marker genes were available (Methods; Supplementary Table 2). We found that all methods produced cell-type proportions (Fig. 3, columns 2-7) that corresponded well with the spatial patterns of marker gene expression (Fig. 3, column 1).

**Figure 3.**
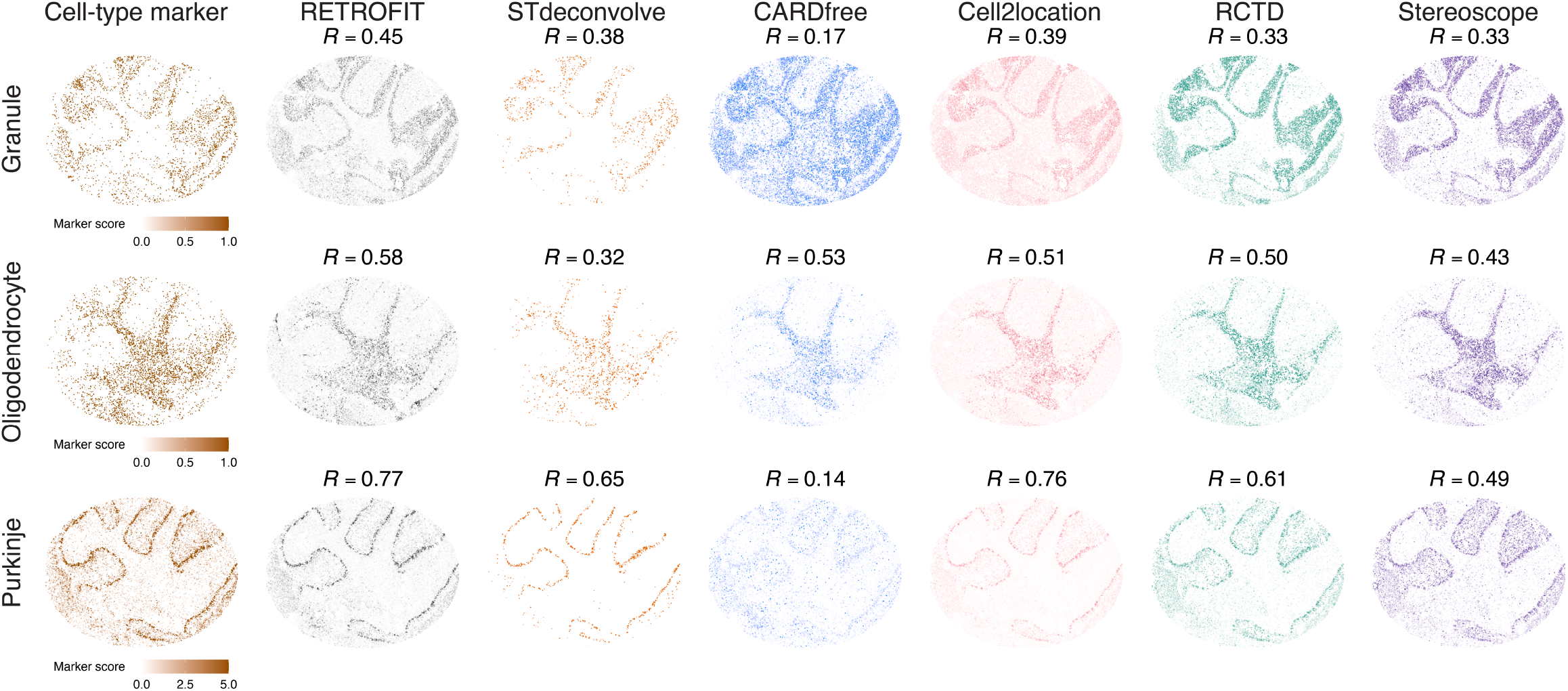
Benchmarking RETROFIT on mouse cerebellum Slide-seq data. Column 1 (leftmost) shows spatial patterns of ST expression scores (Methods) for curated cell-type marker genes in granule cells, oligodendrocytes, and Purkinje cells. Columns 2-7 show cell-type proportions at each spot estimated by each of the 6 ST deconvolution methods. Pearson correlations (*R*) between cell-type marker ST expression scores and estimated cell-type proportions are shown for all methods and cell types.

To quantify differences, we calculated the correlation between estimated cell-type proportions and marker-gene ST expression scores across all spots for each cell type (Methods). RETROFIT achieved the highest correlation for all three cell types (Fig. 3; Supplementary Tables 3-5). Notably, RETROFIT consistently outperformed other reference-free methods, with significant improvements for specific cell types (STdeconvolve: oligodendrocytes; CARDfree: granule and Purkinje cells). It also slightly surpassed Cell2location, the best performing reference-based method, despite using no single-cell reference during deconvolution. Together, these results demonstrate that RETROFIT outperforms existing deconvolution methods on Slide-seq dataset, consistent with our simulation studies.

### RETROFIT identifies fine anatomical structures directly from ST data

Beyond benchmarking accuracy, we also assessed RETROFIT’s ability to uncover biological structure directly from spatial transcriptomics measurements, entirely without the annotation step that uses reference profiles or marker genes. Applying RETROFIT to a mouse main olfactory bulb (MOB) dataset ^32^, we recovered a latent component corresponding to the rostral migratory stream (RMS)—a narrow region containing neuronal precursor cells that is not detectable by clustering^28^. This component localized to the expected anatomical region and exhibited high expression of canonical RMS markers (*Nrep, Sox11, Dcx*). High-resolution in situ hybridization (ISH) patterns for these genes corroborated the RETROFIT-derived structure. A full analysis is provided in the Supplementary Note 4.

### RETROFIT demonstrates strong performance on ultra–high-resolution Visium HD data

To assess RETROFIT’s applicability to emerging single-cell and subcellular-resolution spatial platforms, we analyzed a Visium HD colorectal cancer dataset ^33^, which provides 2*µ*m subcellular bins. Following 10x Genomics recommendations, we aggregated the raw data to 8*µ*m × 8*µ*m bins and applied RETROFIT to the resulting ST matrix (Methods). For comparison, we evaluated RCTD, Cell2location, and STdeconvolve. CARDfree was not included in this analysis because it requires a minimum number of marker genes per cell type, whereas the Visium HD dataset provides only one or two markers for validation, making it unsuitable for this setting.

RETROFIT was run with *L* = 10 latent components and components were annotated post hoc using Algorithm 2 by correlating RETROFIT-derived expression profiles with the accompanying scRNA-seq reference. The same annotation procedure was used for STdeconvolve. RCTD and Cell2location were run with the reference profiles provided in the original study. As in^33^, deconvolution accuracy was assessed by computing the Pearson correlation between spatial expression of five marker genes—*CEACAM6* (Tumor), *COL1A1* (Fibroblast), *TRAC* and *CD3E* (T cells), and *PECAM1* (Endothelial)— and the estimated proportions of their corresponding cell types.

Across the four evaluated cell types (Figure 4), RETROFIT achieved higher correlations than RCTD for three cell types and comparable performance for the remaining one. RETROFIT also outperformed Cell2location for two cell types, performed slightly worse for one, and was comparable for the last—despite operating entirely without single-cell reference information during deconvolution. STdeconvolve showed substantially lower correlations than other methods across all cell types. To verify that this was not due to component annotation in STdeconvolve, we report its highest achievable correlation for each cell type in the Supplementary Note 5; performance remained substantially lower.

**Figure 4.**
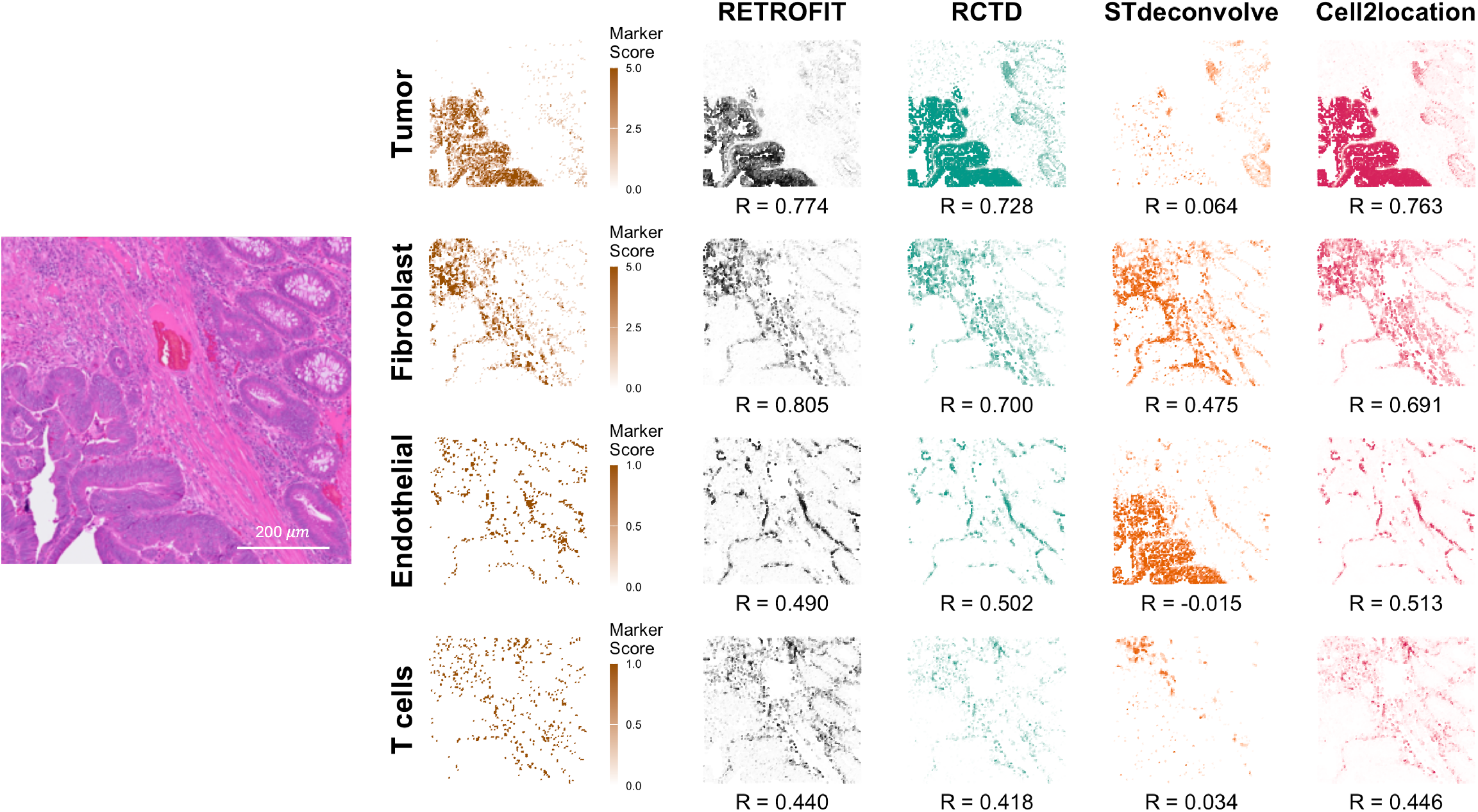
Benchmarking RETROFIT on Visium HD data. Left: Histological image of the colorectal cancer tissue section. Scale bar in this panel represents 200 *µ*m. Right: Rows correspond to four cell types (with markers): Tumor (CEACAM6), Fibroblast (COL1A1), Endothelial (PECAM1), and T cells (TRAC, CD3E). Column 1 shows spatial ST marker expression scores for each cell type (Methods). Columns 2–5 report Pearson correlation coefficients (*R*) between marker expression scores and the estimated cell-type proportions produced by each deconvolution method. All spatial plots correspond to the same tissue area at a resolution of 8 *µ*m per spot.

Together, these results show that RETROFIT accurately captures fine-grained cellular structure even at near–single-cell resolution, demonstrating strong robustness and suitability for emerging ultra–high-resolution platforms such as Visium HD.

### RETROFIT extracts relevant cellular compartments from human intestine Visium data

We applied RETROFIT to a Visium spatial gene expression study of human intestinal development ^10^. This study provided ST data of 33,538 genes and 9,330 spots on intestinal tissues from adults and from fetuses at 12 and 19 post-conceptual weeks (PCW). The study also provided scRNA-seq data on fetal samples, revealing 101 intestinal cell types categorized into 8 cellular compartments with distinct transcriptional signatures: endothelial, epithelial, fibroblast, immune, muscle, myofibroblast (MyoFB)/mesothelial (MESO), neural and pericyte. For each of the three developmental stages, we selected the ST slide with the clearest anatomical markings (Fig. 5a) and input the ST expression count matrices to RETROFIT after quality control (Methods). Specifically, we combined the 37 curated marker genes^10^ from the 8 compartments with significantly overdispersed genes ^28^ across spots identified in each ST slide to construct a matrix of 722 genes and 1080 spots for 12 PCW, a matrix of 681 genes and 1242 spots for 19 PCW, and a matrix of 1051 genes and 2649 spots for adult. To reduce computation and avoid ambiguity caused by a large number of highly correlated cell types, we estimated the proportions of these 8 distinct compartments at each tissue-covered spot of fetal and adult intestinal samples using RETROFIT. The most abundant compartment identified at each spot is shown in Fig. 5b.

**Figure 5.**
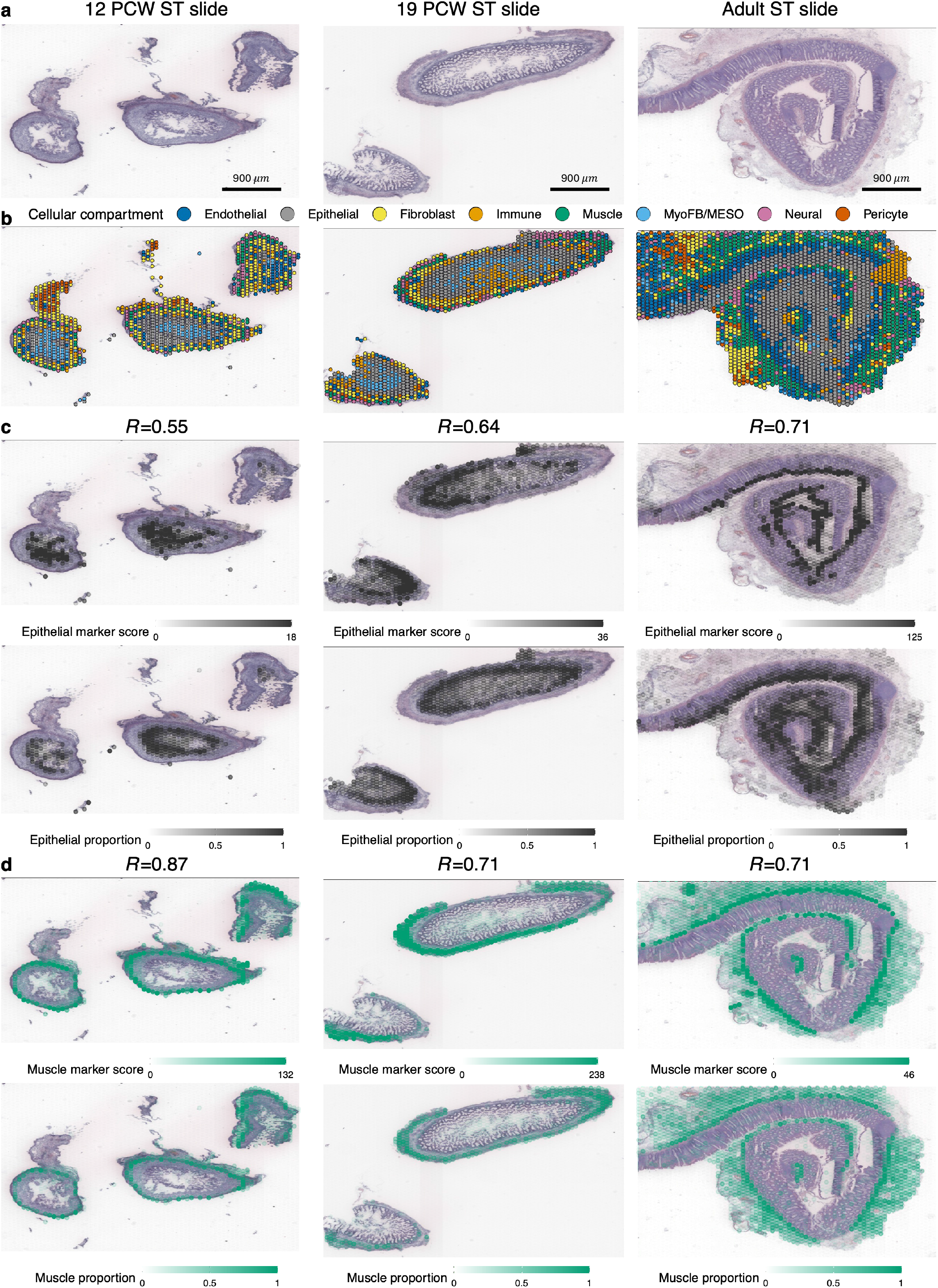
Cellular compartments identified by RETROFIT in human fetal and adult intestine Visium data. **a** H&E images of human fetal (12 and 19 PCW) and adult intestinal tissues. Scale bar in this panel represents 900 *µ*m. **b** Localization of all 8 cellular compartments in each ST slide, marked by the compartment with the largest proportion estimate at each spot. **c**-**d** ST expression scores of compartment marker genes (row 1) and RETROFIT estimates of compartment proportion (row 2) across spots for **c** epithelial and **d** muscle compartments in 3 ST slides. Pearson correlation (*R*) between compartment marker ST expression scores and compartment proportion estimates across all spots is shown for every combination of cellular compartments and developmental stages in **c** and **d**. Expression scores of marker genes for each cell type are calculated as described in Methods section “Human intestine data analysis”.

To estimate compartment proportions at each spot, we matched the *L* = 16 latent components extracted by RETROFIT to the *K* = 8 cellular compartments. Unlike the simulations and the mouse cerebellum Slide-seq study, which had tissue-matched scRNA-seq data for all ST slides, the human intestine study^10^ only provided scRNA-seq data for the 12 and 19 PCW stages, but not the adult stage. Consequently, we annotated the RETROFIT-extracted components using a curated list of 37 intestinal compartment marker genes^10^ for all three stages (Algorithm 3; Supplementary Tables 6-8). This marker-based approach was utilized in all our primary analyses for the human intestine ST data. As an alternative, we also annotated the RETROFIT results for 12 and 19 PCW stages using compartment-specific gene expression derived from the corresponding scRNA-seq data (Algorithm 2; Supplementary Tables 9-10). We then compared the compartment proportions from the two annotation strategies for the same ST slide. For both 12 and 19 PCW stages, the proportion estimates from the two strategies were concordant across spots in 4 compartments: muscle (12 PCW: *R* = 0.93; 19 PCW: *R* = 0.92), endothelial (12 PCW: *R* = 0.85; 19 PCW: *R* = 0.92), fibroblast (12 PCW: *R* = 0.60; 19 PCW: *R* = 0.88) and epithelial (12 PCW: *R* = 0.49; 19 PCW: *R* = 0.86). Additionally, the two strategies produced highly comparable proportions across spots in the immune compartment at 12 PCW (*R* = 0.86) and the neural compartment at 19 PCW (*R* = 0.94).

To evaluate the performance of the two annotation strategies, we examined the correlation between compartment proportions estimated by each strategy and compartment marker ST expression scores across all spots for each compartment and slide (Supplementary Table 11). For the four compartments where the two annotation strategies produced consistent results in the 12 and 19 PCW samples, their proportion estimates from both strategies were positively correlated with the corresponding marker ST expression scores across spots (*R* > 0.54). In compartments where the results of two annotations differed, the proportion estimates based on the marker annotation aligned better with the compartment marker ST expression scores than those based on the scRNA-seq annotation. For example, MyoFB/MESO marker ST expression scores were positively correlated with the marker-based proportion estimates of MyoFB/MESO across spots for both stages (12 PCW: *R* = 0.34; 19 PCW: *R* = 0.61), whereas they were negatively correlated with the proportion estimates based on the scRNA-seq annotation (12 PCW: *R* = −0.09; 19 PCW: *R* = −0.20). Furthermore, we aggregated the compartment-specific correlations into a single metric to compare the overall performance of the two annotation strategies (Methods). We found that Algorithm 3 consistently outperformed Algorithm 2 based on this metric in both ST slides (Supplementary Table 12). Together, these results support the use of the marker-based annotation strategy (Algorithm 3) in the RETROFIT analysis of human intestine ST data.

Across all three stages of intestinal development, RETROFIT’s estimated cellular compartment proportions correlated well with the anatomical locations and ST profiles of compartment-specific marker genes for most cell comparments (Figs. 5b-d; Supplementary Fig.s 7-12). We further compared RETROFIT with Cell2location – the top-performing reference-based method – and two reference-free methods, STdeconvolve and CARDfree. This comparison was limited to ST data from two fetal samples with companion scRNA-seq data, as Cell2location requires a sample-matched single-cell gene expression reference.

Across the 8 compartments at both fetal stages (Table **??**; Supplementary Table 13), RETROFIT produced a significantly higher median correlation than both reference-free methods (one-sided Wilcoxon signed rank *P* = 7.6 × 10^−5^ for STdeconvolve and *P* = 1.4 × 10^−2^ for CARDfree). Notably, even without access to single-cell transcriptomic information during deconvolution, RETROFIT still slightly outperformed Cell2location, yielding higher correlations in 10 of 16 compartment-stage combinations (Table **??**). Certain compartments remained challenges for all methods. For example, for the immune compartment in the 19 PCW slide, all three reference-free methods produced negative correlations, while Cell2location managed only a marginally positive correlation close to zero (*R* = 7.8 × 10^−2^). In some cases, RETROFIT and other reference-free approaches substantially outperformed CELL2location, such as MyoFB/MESO across both slides. These findings underscore the complementary value of reference-free methods like RETROFIT in analyzing complex tissues, where reference-based approaches alone may be insufficient^6^. Overall, the results demonstrate RETROFIT’s competitive performance relative to existing methods, consistent with trends observed in simulation studies (Fig. 2; Supplementary Figs. 1–6) and in the mouse cerebellum Slide-seq data (Fig. 3).

### RETROFIT identifies spatiotemporal patterns of cellular composition in intestinal development

The cellular compositions inferred by RETROFIT on the ST samples of 3 developmental stages shed light on the temporal dynamics in human intestine development (Fig.s 6a-b). The 12 PCW slide had more than twice as high an average proportion of fibroblasts as the other two stages (12 PCW: 24.4% across 1080 spots, 19 PCW: 11.0% across 1242 spots, adult: 11.4% across 2649 spots), aligning with abundant presence of stromal 1–4 (S1–S4) fibroblasts ^10^ in the formation of submucosal structure (S1), crypt-villus axis (S2), enteric vasculature (S3) and lymphoid tissue (S4) during early intestinal development. The 19 PCW slide had the highest average proportions of epithelial (26.3%) and immune (15.6%) cells, indicating the maturation of fetal intestinal epithelium and lymphoid tissue to form the structural basis for essential functions of nutrient absorption and host immunity^10,34^. The adult slide had the highest average proportions of endothelial (19.8%) and muscle (15.8%) cells, reflecting the fully developed enteric vessels and smooth muscle layers in the mature intestine ^10^.

The vast majority of spots in all 3 ST slides encompassed cells from multiple intestinal compartments (Fig. 6b). To help elucidate the dynamics of cell-type complexity across intestinal development, we categorized spots into 3 groups based on their cellular diversity estimated by RETROFIT (Fig. 6c). Group 1 comprised spots dominated by a single compartment, where at least 50% of cells in each spot belonged to one compartment (Fig.s 6d diagonals and 6e). These spots mark regions in a tissue slide dominated by a single cellular compartment. Group 2 comprised spots with at least two moderately representative compartments, each contributing between 25-50% to the spot’s compartment composition (Fig.s 6d off-diagonals and 6f-h). These spots indicate boundaries between two compartments in the slide. Group 3 comprised spots with highly heterogeneous composition, with at most one compartment contributing 25-50% and no other compartment proportion exceeding 25%. These spots represent regions with highly complex compositions of cell types.

**Figure 6.**
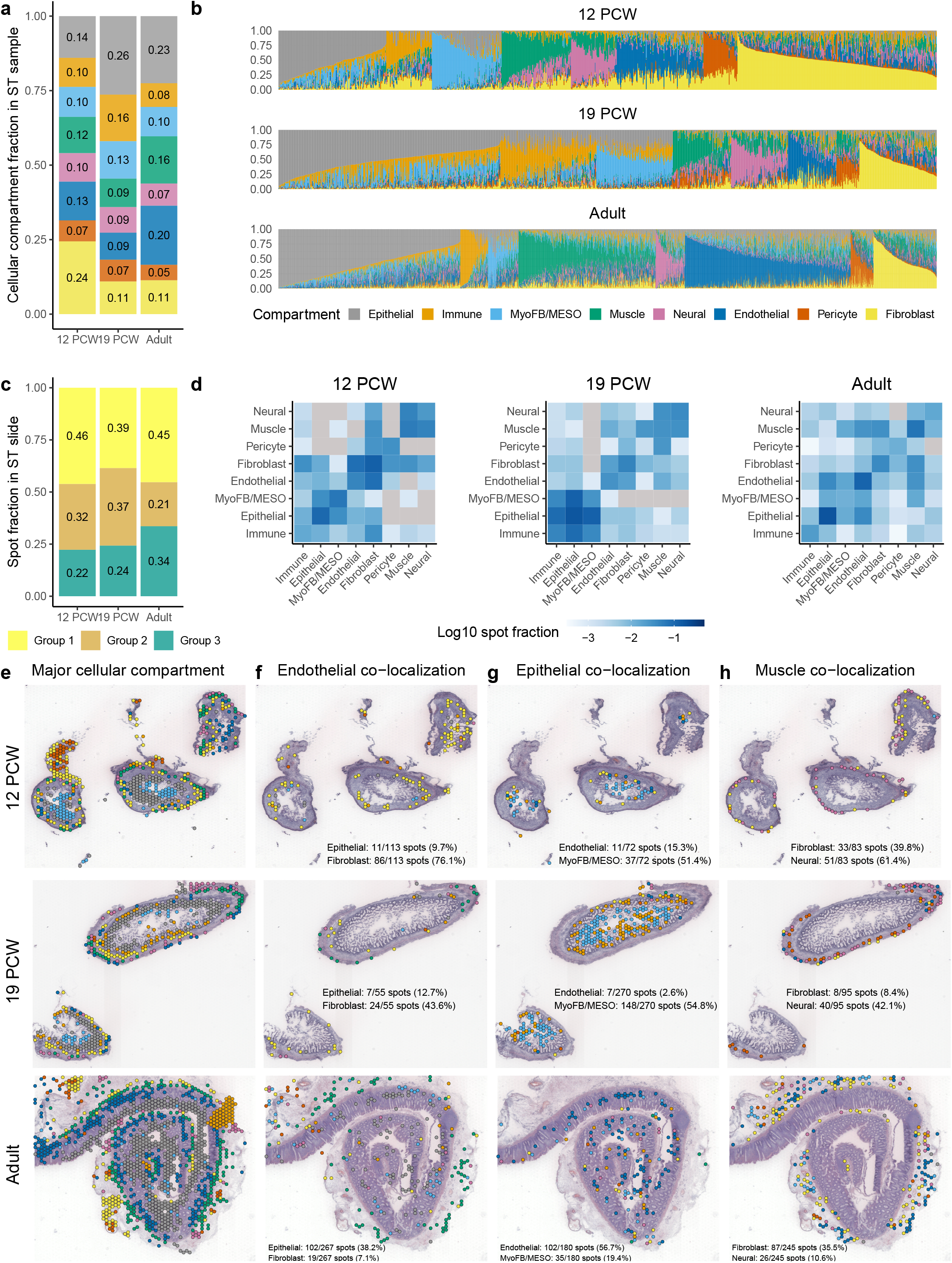
Cellular compositions and spatiotemporal patterns identified by RETROFIT in human intestinal development. **a** Distribution of 8 cellular compartments across all spots in each ST slide. **b** Compartment composition of each spot in each ST slide. **c** Distribution of spots with 3 levels of cellular diversity in each slide. Group 1: spots with a dominant compartment. Group 2: spots with at least two moderately representative compartments. Group 3: spots with highly heterogeneous composition. **d** In each heatmap, each off-diagonal entry shows the fraction of Group 2 spots for each compartment pair, and each diagonal entry shows the fraction of Group 1 spots for each compartment. The off-diagonal entry colored in grey indicates that the number of Group 2 spots is 0 for the corresponding compartment pairs. **e** Spatial distribution of spots with a dominant compartment (Group 1) in each ST slide. **f**-**h** Spatial distribution of spots with at least two moderately representative compartments (Group 2), with the anchor compartment as **f** endothelial, **g** epithelial or **h** muscle compartment. The color of each spot in **f**-**h** represents the other cellular compartment that co-localizes with the anchor compartment and has the largest proportion estimate. Counts and percentages of Group 2 spots for 6 pairs of co-localized compartments are shown in **f**-**h**.

Compositions of the 3 spot groups varied across development (Fig.s 6c-d). Group 1 spots were the most prevalent in all three stages (12 PCW: 46.1%; 19 PCW: 38.5%; adult: 45.3%), and they exhibited layering and clustering patterns that matched known cellular anatomy of the human intestine ^10^, particularly evident in the adult sample (Fig. 6e). Group 2 spots were less common in the adult sample than fetal samples (12 PCW: 31.6%; 19 PCW: 37.2%; adult: 21.1%), but they exhibited a higher degree of pairwise cellular diversity in the adult sample. Out of 28 possible pairwise co-localization patterns among 8 cellular compartments, 27 were present in Group 2 spots for the adult sample, compared to 20 and 24 for 12 and 19 PCW samples respectively (Fig. 6d). The adult sample also had the largest fraction of Group 3 spots (12 PCW: 22.3%; 19 PCW: 24.3%; adult: 33.6%), highlighting the intricate composition of cell-types in the adult intestine. Taken together, the dynamics of spot-level cellular diversity inferred by RETROFIT effectively captures the increasing complexity of cellular compositions as the human intestine develops.

We then examined the co-localization patterns of 8 cellular compartments in Group 2 spots across the 3 developmental stages. We identified some commonalities in cellular co-localization across intestinal development (Fig. 6d). For example, muscle cells consistently exhibited the highest prevalence of co-localization with neural cells in Group 2 spots across all stages (12 PCW: 51/69 spots, 73.9%; 19 PCW: 40/56 spots, 71.4%; adult: 26/68 spots, 38.2%; Supplementary Fig. 13), recapitulating the intestinal anatomy that myenteric plexuses are surrounded by muscles ^10^. Similarly, epithelial cells consistently displayed the highest prevalence of co-localization with immune cells across Group 2 spots throughout development (12 PCW: 12/62 spots, 19.4%; 19 PCW: 112/166 spots, 67.5%; adult: 18/53 spots, 34.0%; Supplementary Fig. 13), highlighting the crucial role of epithelial cells in mediating homeostasis of immune cells in the intestine ^35^.

Notably, distinct cellular co-localization patterns emerged in Group 2 spots between fetal and adult samples (Fig. 6d). In both fetal stages, fibroblasts were the most common in Group 2 spots co-localized with endothelial cells (Fig. 6f), supporting the coordination of S3 fibroblasts and endothelial cells during fetal intestinal angiogenesis^10^. In the adult sample, however, epithelial cells prevailed in Group 2 spots co-localized with endothelial cells (Fig. 6f). The endothelial-epithelial co-localization in the adult sample, which was obtained from a patient undergoing intestinal surgery^10^, aligns with a recent mouse study showing that lymphatic endothelial cells reside in proximity to crypt epithelial cells and support renewal and repair of intestinal epithelium after injury ^36^. Contrasting the predominant co-localization of endothelial and epithelial compartments in the adult sample, MyoFB/MESO compartment prominently co-localized with epithelial cells in both fetal samples (Fig. 6g). This finding reflects the signaling circuit between epithelial stem cells and myofibroblasts during fetal intestinal development^10^. Moreover, cellular co-localization of muscle cells also exhibited temporal variation across stages. For Group 2 spots co-localized with muscle cells, neural cells were predominant in the fetal stages, while fibroblasts were the most common in the adult stage (Fig. 6h). This difference can be attributed to the role of S1 fibroblasts in forming submucosa structures that join mucosa to smooth muscle layers of the mature intestine ^10^.

Overall, the reference-free inference of cellular composition and co-localization enabled by RETROFIT provides insights into the dynamic interplay of cellular processes that shapes intestinal development and function, demonstrating the potential for RETROFIT to yield new hypotheses of tissue biology from ST data alone.

### RETROFIT captures cell-type transcriptional specificity without using single-cell references

RETROFIT estimates cell-type-specific gene expression and cell-type composition simultaneously (Fig. 1). In simulations we demonstrated the high correlation between cell-type-specific transcriptional profiles estimated by RETROFIT and those measured by single-cell technologies (Fig. 2e). Here we examined compartment-specific transcriptional profiles estimated by RETROFIT on the human intestine ST data (Supplementary Tables 20-25).

We first compared RETROFIT-estimated compartment-specific expression with observed single-cell expression for 37 curated marker genes^10^ across 8 cellular compartments (Fig. 7a; Methods), using companion scRNA-seq data from 12 and 19 PCW stages. For each marker gene, we computed Pearson correlation between RETROFIT estimates and scRNA-seq measurements across cellular compartments. Of the 37 marker genes, 25 (67.6%) showed high correlation (*R* > 0.95) in at least one stage, and 12 (32.4%) in both. Many of these 12 genes, such as *ACTG2* (muscle), *PECAM1* (endothelial), *PHOX2B* (neural) and *PTPRC* (immune), exhibited strong cellular specificity as expected.

**Figure 7.**
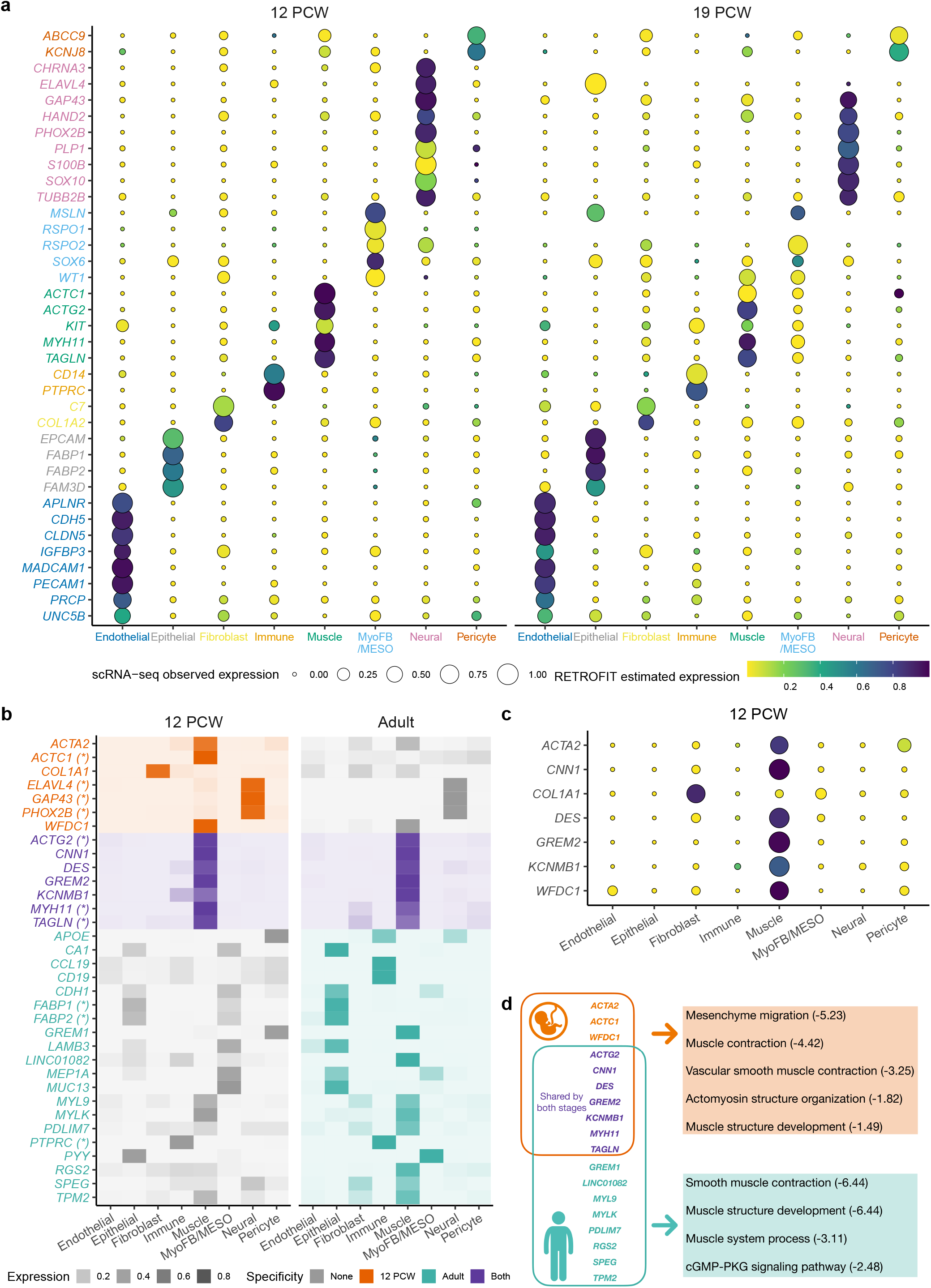
Transcriptional signatures and biological pathways identified by RETROFIT in human intestinal development. **a** Normalized expression of 37 marker genes for 8 cellular compartments in two fetal stages, obtained from RETROFIT estimates and scRNA-seq data. Each marker gene (*y*-axis) has a matching color with the compartment it characterizes (*x*-axis). **b** Normalized expression of 34 putative compartment-specific genes estimated by RETROFIT for 12 PCW and adult stages. Gene colors represent the compartment-specific transcriptional specificity identified in 12 PCW (orange) or adult (green) or both stages (purple). Grey colors indicate that genes were not identified as compartment-specific in a given stage. Asterisks (*) indicate that the identified genes are also markers in **a. c** Normalized expression of 7 compartment-specific non-marker genes obtained from RETROFIT estimates and scRNA-seq data for 12 PCW stage. The 7 genes were identified by RETROFIT in **b** but were not labeled as markers in **a. d** Top-ranked biological pathways enriched in muscle-specific genes identified by RETROFIT in **b** for 12 PCW and adult stages (FDR < 0.05), with the multiplicity adjusted enrichment *P*-value (FDR) in log base 10 shown after each pathway.

To further demonstrate RETROFIT’s ability to uncover cell-type-specific expression patterns without single-cell references, we identified compartment-specific genes using RETROFIT estimates alone. To ensure reliable results, we only considered stages (12 PCW and adult) with biological replicates (Supplementary Tables 22-25) and selected genes showing consistent patterns of high expression (count > 40) and strong compartment specificity (entropy < 1.5 and Gini index > 0.85) across all replicates in a given stage (Methods). Despite stringent criteria, RETROFIT identified 34 compartment-specific genes (14 in 12 PCW and 27 in adult stages), 7 of which were shared between stages (Fig. 7b). Among them, 7 (50.0%) and 6 (22.2%) overlapped with the curated compartment markers (Fig. 7a) at 12 PCW and adult stages, respectively ^10^.

For genes not included in our curated list, RETROFIT’s estimated compartment-specific expression strongly agreed with scRNA-seq measurements at 12 PCW sample (Pearson *R* = 0.96, *P* = 3.1 ×10^−31^), confirming their biological relevance. Many have been independently reported as cell-type-specific in other studies. For example, *COL1A1*, identified by RETROFIT as fibroblast-specific, encodes a fibril-forming collagen broadly expressed in connective tissues^37^, and *DES*, identified as muscle-specific, encodes an intermediate filament critical for muscular function^38^. These assignments are consistent with a recent mouse scRNA-seq study^39^, which identified *COL1A1* as a fibroblast marker and *DES, CNN1* and *ACTA2* as mural cell markers. These findings show that RETROFIT accurately recovers biologically meaningful, cell-type-specific genes directly from ST data alone, even without prior reference information.

We next examined the developmental regulation of these compartment-specific genes (Fig. 7b). RETROFIT revealed stage-specific transcriptional programs, with 27 of 34 showing specificity in only one stage. For example, 3 known neural marker genes (*ELAVL4, GAP43, PHOX2B*) were identified as neural-specific exclusively in 12 PCW but not in the adult stage, consistent with their roles in neuronal differentiation (Supplementary Table 26) and embryonic ventral midbrain development (Supplementary Table 27). In contrast, *FABP1, FABP2* and *MUC13* were identified as epithelial-specific only in adults. They are known to participate in the digestive and absorptive functions (Supplementary Table 26), and show strong transcriptional specificity for epithelial cells in multiple single-cell transcriptomic studies of human intestinal tissues (Supplementary Table 27). Since nutrient absorption manifests late in intestinal development (typically after villus formation ^34^), these adult-specific genes likely represent functional maturation signatures in the mature intestinal epithelium. These support RETROFIT’s ability to capture transient, developmentally regulated transcriptional activity.

RETROFIT also uncovered tissue-context-specific smooth muscle programs. It identified 10 muscle-specific genes at 12 PCW and 15 in adult stages, with 7 shared between stages (Fig.s 7b and d). These genes were significantly enriched for pathways related to muscle contraction and muscle structure development in both stages (12 PCW: FDR = 3.8 × 10^−5^ and FDR = 3.2 × 10^−2^; adult: FDR = 3.6 × 10^−7^ for both; Supplementary Table 26). When compared with single-cell transcriptomes from 15 human organs^40^, these genes showed markedly stronger enrichment in smooth muscle cells from intestinal than from heart or other muscle-rich tissues: 8/10 fetal (FDR = 1.0×10^−16^) and 10/15 adult (FDR = 1.0×10^−20^) vs 4/10 (FDR = 3.2×10^−6^) and 3/15 (FDR = 1.6 × 10^−3^) for heart. This indicates that RETROFIT identifies organ-specific transcriptional programs, rather than generic smooth muscle signatures.

Finally, RETROFIT distinguished developmental differences within the muscle compartment. For example, the mesenchyme migration pathway was significantly enriched only at 12 PCW (FDR = 5.9×10^−6^), driven by two fetal-only muscle-specific genes (*ACTA2* and *ACTC1*), consistent with the known mesothelial-to-mesenchymal transition during vascular smooth muscle formation in fetal stage ^41^.

Together, these analyses demonstrate RETROFIT’s capability to go beyond reproducing known markers—identifying stage-specific, functionally distinct, and tissue-restricted transcriptional programs directly from ST data, capturing biologically meaningful patterns that reflect developmental transitions and spatial context without reliance on single-cell references.

## Discussion

We present RETROFIT, an unsupervised Bayesian framework for reference-free cell-type deconvolution of spatial transcriptomics data. In contrast to reference-based approaches^13–21^ that require matched single-cell transcriptomic profiles, and to marker-gene–assisted reference-free methods ^17,23–25^ that incorporate curated gene sets during inference, RETROFIT performs data decomposition using sequencing measurements alone and incorporates external information only at a post hoc annotation stage when available. This design, shared by a small number of reference-free approaches^28^, enhances robustness to variability in auxiliary data and allows inferred components to remain reusable as biological annotations and reference resources evolve.

Across a diverse set of simulation settings and real ST datasets, RETROFIT shows robust and consistent performance across platforms and resolutions. In controlled simulations, it outperforms existing reference-free methods and remains competitive with reference-based approaches, with clear advantages when single-cell references are incomplete. In real data, RETROFIT achieves strong accuracy on mouse cerebellum Slide-seq and human intestine Visium datasets. Notably, on ultra–high-resolution Visium HD colorectal cancer data, RETROFIT substantially outperforms reference-free alternatives and achieves accuracy comparable to state-of-the-art reference-based methods—despite operating without single-cell references or marker genes during deconvolution. RETROFIT also recovers fine-scale anatomical structures, such as the rostral migratory stream in the mouse main olfactory bulb, directly from ST data alone, demonstrating its capacity for unsupervised discovery of spatially organized biological features. Together with recent work on reference-free deconvolution ^17,23–25,28^, these results highlight the practical value of reference-free approaches for robust spatial deconvolution across platforms, resolutions, and data-availability regimes.

Beyond estimating cell-type composition, RETROFIT jointly infers cell-type-specific gene expression profiles directly from ST data. These in situ expression estimates enable analyses that extend beyond proportion estimation, including identification of stage-specific transcriptional programs, tissue-context-specific signatures, and developmentally regulated pathways. In the human intestine analysis, RETROFIT reveals compartment- and stage-specific transcriptional signatures that align with known biology and provide insights into intestinal development and tissue specialization. Such expression estimates can also be integrated with disease-centric resources – for example, by correlating spatially resolved, cell-type-specific expression with curated disease gene sets^10^ or genome-wide association study signals^42^ – to localize disease-relevant genes to specific cellular and spatial contexts.

Recent deconvolution methods ^26,27,43^ have explored integrating paired histological images to provide complementary morphological information. When high-quality, well-aligned histology from the same tissue section is available, such image-informed approaches can provide valuable additional context. However, histological images are not uniformly available across ST platforms and sometimes obtained from adjacent rather than identical tissue sections, which can complicate integration. RETROFIT is designed to operate robustly using sequencing data alone, enabling deconvolution in scenarios where histology, curated marker genes, or matched single-cell references are unavailable or difficult to integrate. By remaining agnostic to axillary information, while still recovering biologically meaningful spatial structure and expression profiles, RETROFIT offers a broadly applicable and complementary framework for spatial transcriptomics analysis. Additional sensitivity analyses examining the influence of auxiliary marker information are provided in Supplementary Note 6.

As with other reference-free approaches, RETROFIT requires specifying the number of latent components. In practice, we recommend choosing a value that exceeds the expected number of cell types and relying on post hoc annotation to identify biologically meaningful components. While heuristic strategies for automatic component selection exist^28^, our analyses suggest that they may underestimate cellular diversity in complex tissues. More principled alternatives, such as sparsity-inducing priors ^44^ or automatic relevance determination ^45^, could be incorporated within RETROFIT’s Bayesian framework. RETROFIT currently treats spatial transcriptomic spots as exchangeable and does not explicitly model spatial dependence. Nevertheless, it consistently recovers anatomically meaningful spatial patterns and outperforms CARDfree^17^, a reference-free method that incorporates spatial smoothing through conditional autoregressive priors, in our mouse cerebellum and human intestine analyses. These results suggest that much of the spatial structure can be recovered directly from transcriptional variation, while integrating spatial dependence in a flexible and scalable manner represents an important direction for future extensions.

Overall, RETROFIT provides a statistically principled and broadly applicable framework for reference-free deconvolution and in situ transcriptomic characterization. By producing interpretable components and biologically meaningful expression profiles directly from sequencing data, RETROFIT complements image-informed and reference-based approaches and expands the analytical toolkit for spatial biology across diverse tissues, platforms, and experimental contexts.

## Methods

### Bayesian hierarchical model

Let **X** = [*X*_*gs*_] be the *G* × *S* count matrix of expression levels for *G* genes at *S* spots obtained from a ST experiment. Since only a finite number of cell types constitute the ST sample, we represent **X** as a low-rank matrix spanned by *L* non-negative components that capture transcriptional signatures of distinct cell types in the ST sample. Specifically, we model the observed expression level of gene *g* at spot *s, X*_*gs*_, as the sum of unobserved expression counts in *L* latent components:

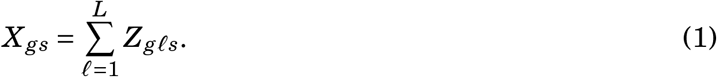

We further attribute each latent component *Z*_*gℓs*_ to two independent sources in an additive manner:

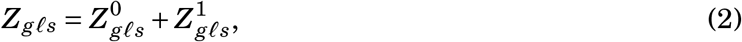

where 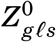 denotes the background expression level shared by all genes in component *ℓ* at spot *s* and 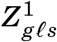 denotes the expression level specific to gene *g* in component *ℓ* at spot *s*. We model the unobserved gene expression counts 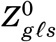 and 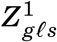 as two independent Poisson random variables ^15,46^:

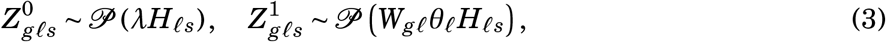

where the background expression levels 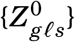 for a given component *ℓ* and a given spot *s* follow the same Poisson distribution across all genes. In Eq. (3), *W*_*gℓ*_ > 0 denotes the average expression level of gene *g* in component *ℓ, H*_*ℓs*_ > 0 denotes the weight of component *ℓ* at spot *s*, and *λ* ≥ 0 denotes an ‘offset’ constant capturing the background expression level shared by all genes across all components and spots ^47,48^. The parameter *θ*_*ℓ*_ > 0 captures the global contribution of component *ℓ* to the measured transcript counts across the entire slide. Biologically, *θ*_*ℓ*_ reflects systematic cell-type–specific differences in measurement efficiency—such as differences in RNA content, capture efficiency, or other biological and technical factors that influence how strongly each component contributes to the observed ST signal beyond its local abundance *H*_*ℓs*_. When the prior on ***θ*** = [*θ*_*ℓ*_] induces shrinkage toward zero, only a subset of components are expected to retain *θ*_*ℓ*_ values that are substantially greater than zero ^44^. This encourages a more parsimonious and interpretable factorization by allowing the model to downweight components that contribute little to explaining the observed ST signal. Taken together, we obtain the following generative model of ST data:

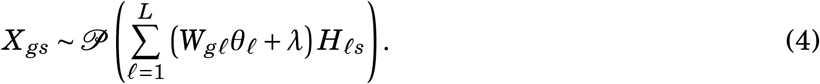

The mean of Poisson model (4) implies two non-negative matrix factorization (NMF) models. When *λ* = 0, the mean of Poisson model (4) implies the Gamma Process NMF^44^ (GaP-NMF): 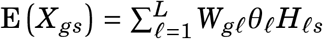. When *λ* = 0 and *θ*_*ℓ*_ = 1 for *ℓ* = 1,…, *L*, the mean of Poisson model implies the standard NMF^49^: 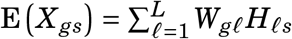. According to various benchmarking analyses in this study, RETROFIT outperforms both the standard NMF and GaP-NMF in deconvolving synthetic and real ST datasets (Supplementary Fig. 14). We take a Bayesian approach to learn the unknown parameters {*W*_*gℓ*_, *θ*_*ℓ*_, *H*_*ℓs*_} in the Poisson generative model (4) from the observed ST data **X**. Specifically, we place independent Gamma priors ^44^ on them:

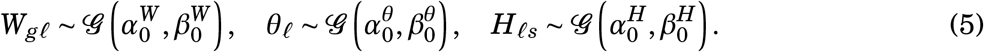

We choose the Gamma priors (5) mainly for computational convenience, because combining the Poisson generative model (4) with the Gamma priors (5) leads to conditional conjugacy, which will simplify the development of SSVI algorithm described in the next section.

### Structured stochastic variational inference

To compute the posteriors of {*W*_*gℓ*_, *θ*_*ℓ*_, *H*_*ℓs*_} , we implement a SSVI algorithm^29^ that scales well with thousands of genes and spots (Supplementary Table 1). To formulate the SSVI algorithm, we use the following notation: 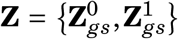 for *g* = 1,… *G* and *s* = 1,…, *S, L*-length vector 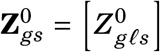 , *L*-length vector 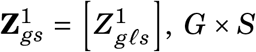 , *G* × *S* matrix **W** = [*W*_*gs*_] , *L*-length vector ***θ*** = [*θ*_*ℓ*_] and *L* × *S* matrix **H** = [*H*_*ℓs*_]. SSVI seeks a variational distribution *q*(**Z, W, *θ*, H**) of the following form to minimize its Kullback–Leibler (KL) divergence to the actual posterior distribution *p*(**Z, W, *θ*, H** | **X**):

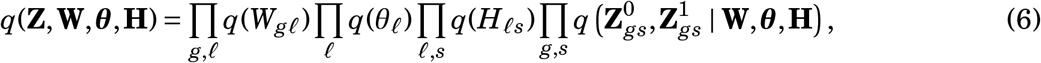

where {*q*(*W*_*gℓ*_), *q*(*θ*_*ℓ*_), *q*(*H*_*ℓs*_)} are required by SSVI to be in the same exponential family as the priors of {*W*_*gℓ*_, *θ*_*ℓ*_, *H*_*ℓs*_}while 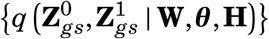 can have any distributional form. By restoring dependence between model parameters {**W, *θ*, H**} and latent variables **Z** through 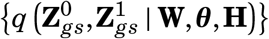 the variational distribution specified by Eq. (6) improves upon the standard mean-field variational distribution that (incorrectly) enforces independence between {**W, *θ*, H**} and **Z**. Consequently, SSVI often outperforms mean-field variational inference on a wide range of Bayesian hierarchical models ^29^.

Since our Bayesian model is defined by the Poisson likelihood (4) and Gamma priors (5), *q*(*W*_*gℓ*_), *q*(*θ*_*ℓ*_), *q*(*H*_*ℓs*_) in Eq. (6) are automatically Gamma distributions, which satisfy the distributional requirement in SSVI:

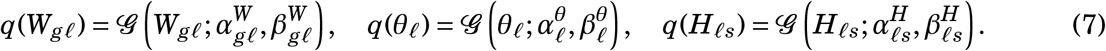

We specify 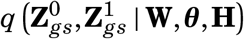 as the exact conditional posterior distributions of 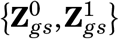 given {**W, *θ*, H**}:

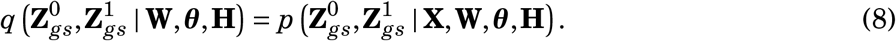

This specification is chosen for two reasons. First, Eq. (8) provides the best possible approximation by achieving zero KL divergence to the actual conditional posterior^29^. Second, because 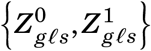 are independent Poisson random variables (3) that constitute the ST expression profile 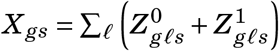, the right-hand side of Eq. (8) has a closed form of a multinomial distribution:

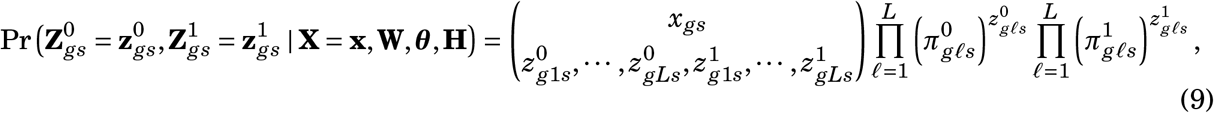

where 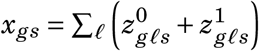 is the observed ST expression count for gene *g* at spot *s* and

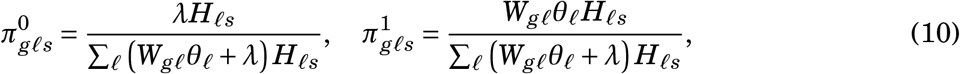

for each component *ℓ*. With the variational distribution defined by Eq.s (6)-(10), we optimize the corresponding variational parameters 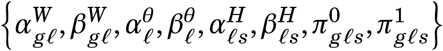 through an iterative and stochastic procedure ^29^ defined in Algorithm 1. The derivation of Algorithm 1 is provided in the Supplementary Note 1. For each synthetic or real ST dataset analyzed in this study, we ran Algorithm 1 with *I* = 4000 iterations.

In this study, we initialize 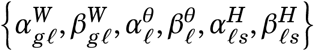 in Algorithm 1 as

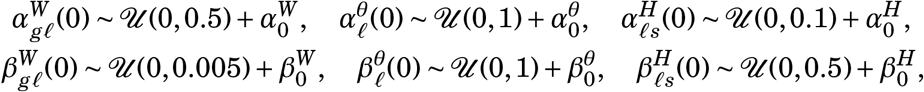

where 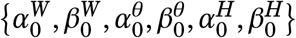 are hyper-parameters specified in the next section and 𝒰 (*a, b*) denotes a continuous uniform distribution on the interval [*a, b*]. In use cases where specific initialization schemes are available, they can be easily used in our R package to run Algorithm 1.

### Hyperparameter specification

When applying RETROFIT in this study, we fixed all the hyperparameters 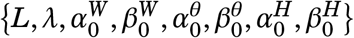 as known constants to further simplify large-scale computation. In addition, we systematically investigated the robustness of RETROFIT with respect to hyperparameter specification, as detailed below. In use cases where specific prior knowledge about 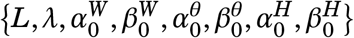 is available, such external information can be further utilized to guide the specification of hyperparameters and can be supplied to the RETROFIT software.

For each dataset analyzed here, we set *L* as twice the number of known cell types in the tissue sample (*K*) to ensure that all cell types present in the ST slide can potentially be captured by the *L* latent components. This choice of *L* follows the recommendation of a related method, GaP-NMF^44^, which demonstrates the effectiveness of using a relatively large *L* in practice. Additionally, we conducted an ablation study to investigate how the performance of RETROFIT is influenced by the choice of *L* using three synthetic ST datasets from Fig. 2. We found that RETROFIT consistently performed well across a broad range of *L* > *K* (Supplementary Fig. 15), including our choice of *L* = 2*K* .

On each synthetic or real ST dataset, we ran RETROFIT with multiple values of *λ* in the Poisson model (4). For the simulated data, we presented RETROFIT results using *λ* = 0.01 in Fig.2. Supplementary Fig. 16 contains additional results using other *λ* values (0, 0.5, 1), which showed comparable performance to *λ* = 0.01. For the analysis of the mouse cerebellum Slide-seq and human intestine Visium datasets, we expanded the range of *λ* to {0, 0.01, 0.05, 0.1, 0.5, 1} and selected the optimal value using a performance metric detailed in the Supplementary Note 3. This metric calculates correlations between estimated and reference cell-type-specific expression profiles for each matched cell type across different *λ* choices and then computes the mean correlation across all *λ* values to determine the gain in correlation for each cell type relative to the mean. (*δ*) The average gain in correlation across all cell types, denoted as *δ*, serves as the overall performance metric. A higher *δ* indicates a more suitable parameter choice. Based on this metric, we chose *λ* = 0.05 for the analysis of mouse cerebellum Slide-seq data and *λ* = 0.01 for the analysis of human intestine Visium data (Supplementary Table 28).

For all datasets, we set the hyper-parameters in Gamma priors (5) as 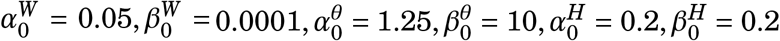, also informed by GaP-NMF^44^. In particular, the Gamma prior on the component contribution 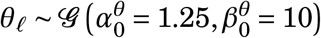 has a mean of 0.125 and variance of 0.0125, shrinking *θ*_*ℓ*_ values towards zero for most components and favoring a simpler model with fewer components in practice. To assess RETROFIT’s sensitivity to these Gamma hyper-parameters, we reran RETROFIT on each of the three synthetic ST datasets from Fig. 2 using four different sets of 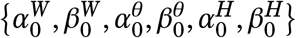 and obtained similar results (Supplementary Fig. 17).

#### Algorithm 1

SSVI for reference-free decomposition of ST data matrix

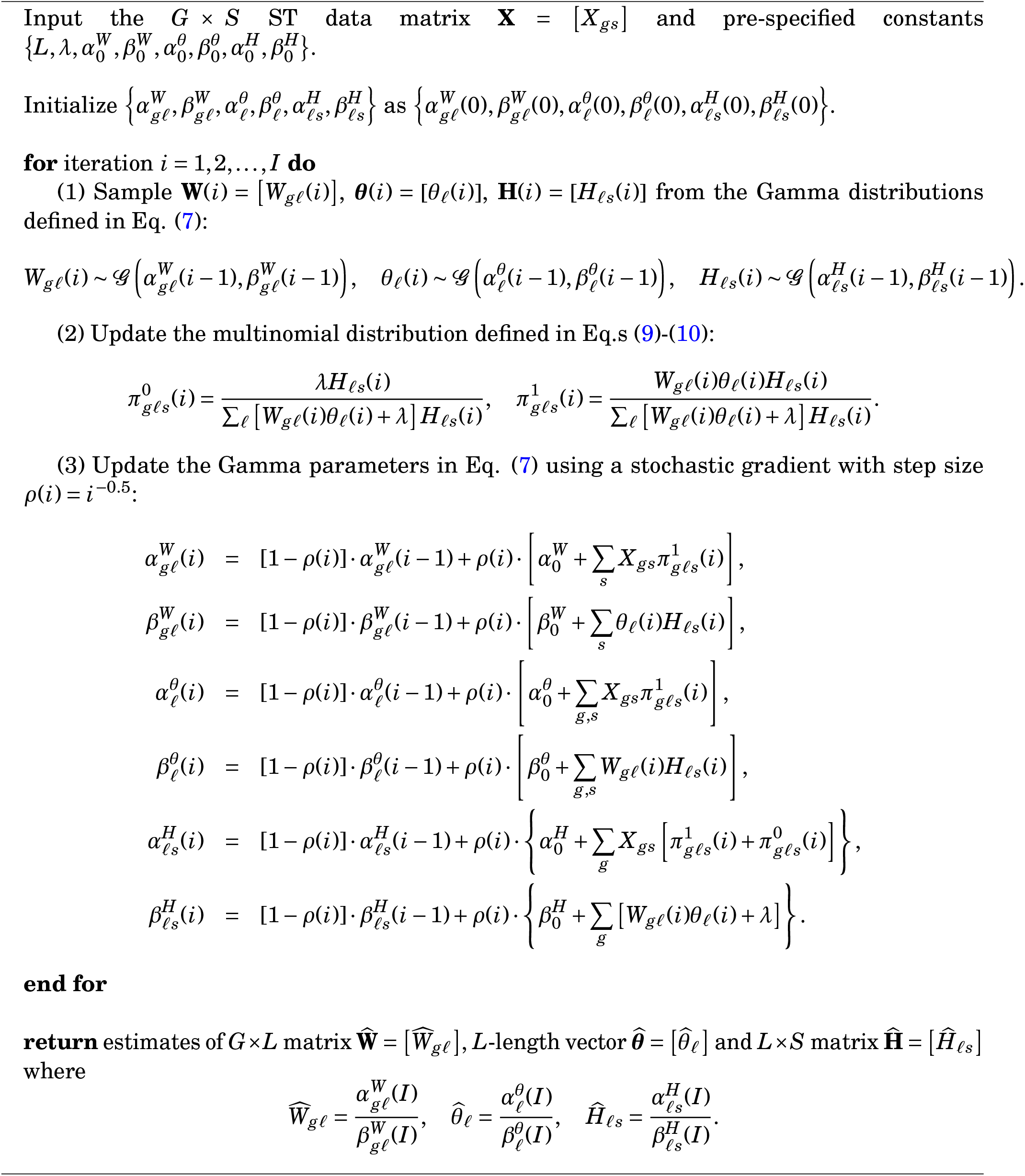

### Cell-type annotation strategies

After running Algorithm 1 on the ST data matrix, the expression profile of each gene at each spot is deconvolved into *L* latent components represented by columns of the *G* × *L* matrix **Ŵ**. To map the *L* latent components to *K* known cell types present in the ST data, we develop two simple strategies (Fig. 1). The first approach is suitable when a reference of cell-type-specific gene expression is available, such as cell-type-annotated scRNA-seq data from the same tissue type. It computes correlations between the deconvolved component-specific expression profiles **Ŵ** and the cell-type-specific expression profiles (**W**^0^), and then matches each component to the cell type with the highest correlation (Algorithm 2). The second approach is suitable when marker genes for relevant cell types in the ST sample are known. It calculates a marker expression score for each component in each cell type (**M**), defined as the sum of normalized component-specific expressions of known marker genes in this cell type, and then annotates each component with the cell type having the largest score (Algorithm 3).

When both cell-type-specific gene expression references and marker gene lists are available, either method can be used. Our simulations show that these approaches often produce consistent results when high-quality references and markers are employed (Supplementary Fig. 18). However, annotation performance is contingent on the quality of single-cell references or cell-type markers, which can vary across studies. To address this practical issue, we use the performance metric (*δ*) for selecting the optimal *λ* (Supplementary Note 3) to aid in choice of the appropriate annotation strategy for real ST data analysis. We compute the *δ* metric for each annotation strategy and select the one with the higher *δ* value (Supplementary Table 12). Using this criterion, Algorithm 2 was chosen for the primary analysis of mouse cerebellum Slide-seq data, while Algorithm 3 was selected for the primary analysis of human intestine Visium data.

After matching the latent components to known cell types, we obtain a cell-type-specific expression matrix for all genes 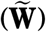 and a cell-type proportion matrix for all spots 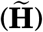 as follows. Let ℒ = {*ℓ*_1_, *ℓ*_2_, … , *ℓ*_*K*_} ⊆ {1, 2 … , *L*} denote the set of latent components matched to *K* cell types, where *ℓ*_*k*_ indicates that the *ℓ*_*k*_-th column of the *G* × *L* matrix **Ŵ** is matched to cell type *k*. We extract these columns from **Ŵ** to form a *G* × *K* matrix 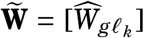, where *g* = 1, 2,…, *G* and *k* = 1, 2,…, *K* . This matrix 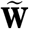 represents the cell-type-specific expression estimates of *G* genes in *K* cell types. Similarly, we extract the rows in **Ĥ** corresponding to the cell-type-matched columns of **Ŵ** and then normalize them to estimate the proportions of *K* cell types at *S* spots. We denote this *K* × *S* matrix 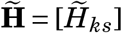, where 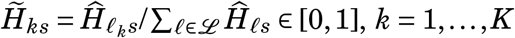, *k* = 1,…, *K* and *s* = 1, 2,…, *S*.

### Existing methods for comparison

In light of several recently published benchmark studies for cell-type deconvolution of ST data^18–21^, we selected five top-performing methods to compare performance with RETROFIT:

#### Algorithm 2

Cell-type mapping based on cell-type-specific gene expression

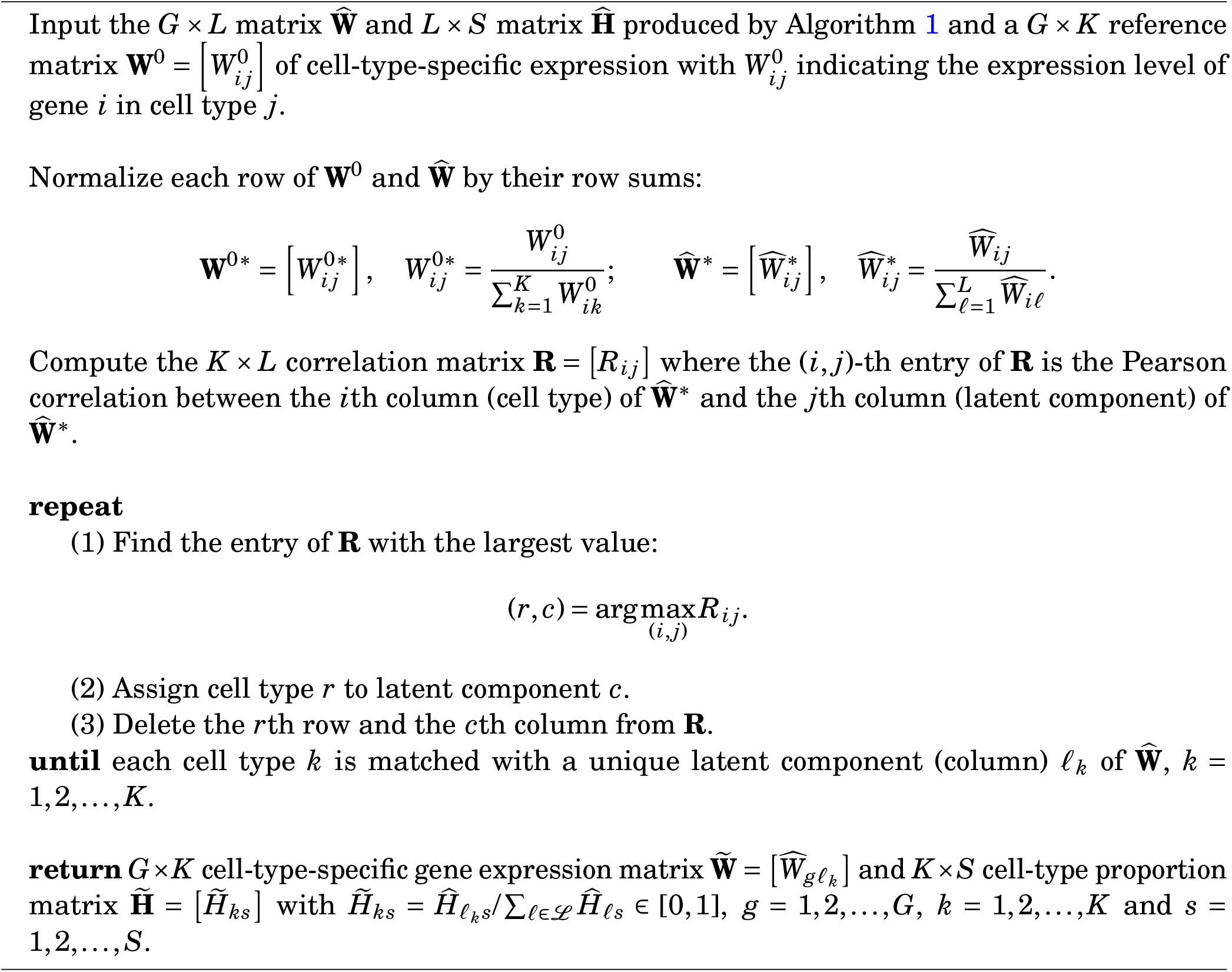

#### Algorithm 3

Cell-type mapping based on cell-type-specific marker gene list

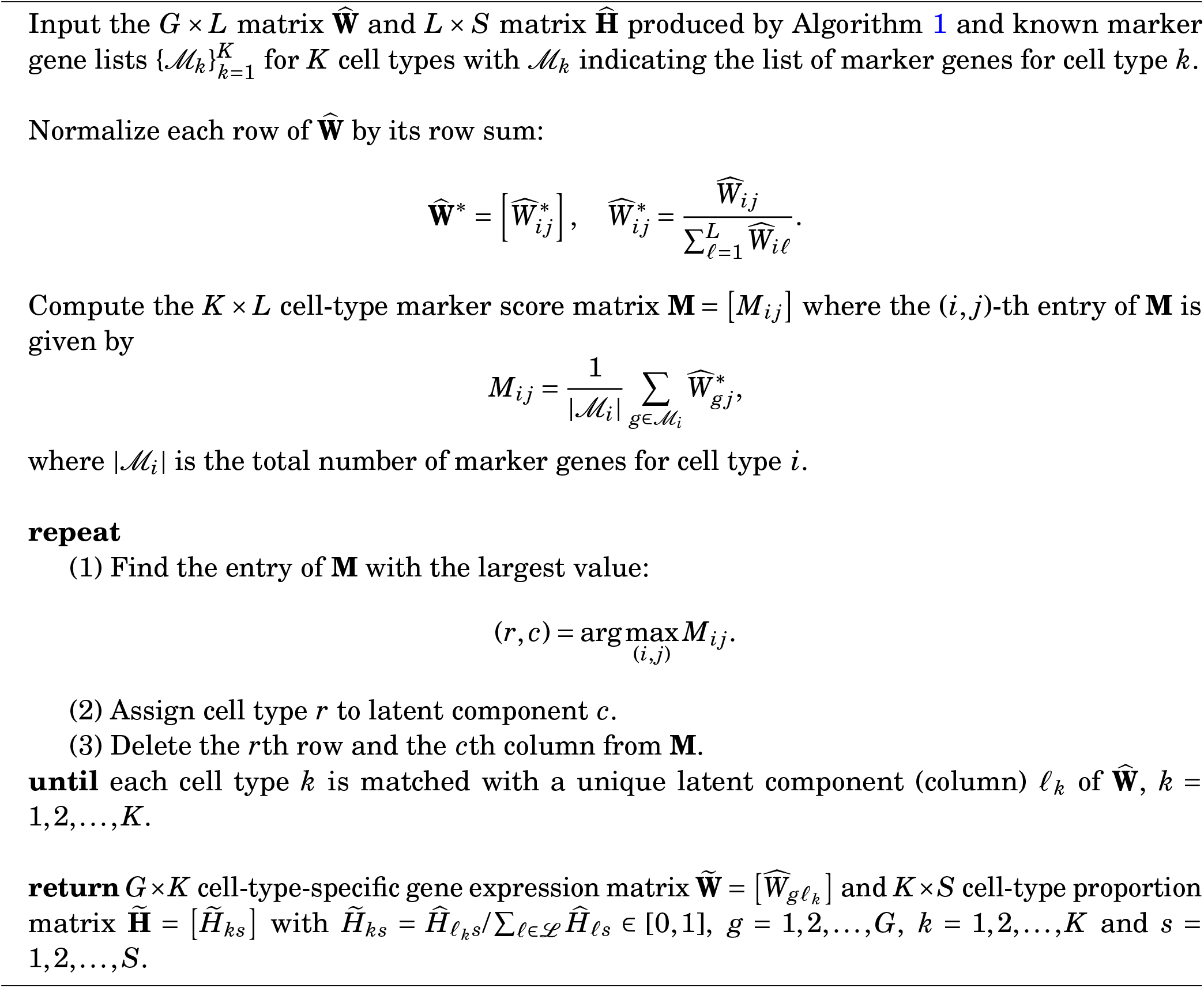

CARDfree ^17^ (https://yma-lab.github.io/CARD, version 1.1), Cell2location ^16^ (https://github.com/BayraktarLab/cell2location, version 0.1.3), STdeconvolve^28^ (https://bioconductor.org/packages/STdeconvolve/, version 1.4.0), RCTD ^15^ (https://github.com/dmcable/spacexr, version 2.2.1), and Stereoscope^13^ (https://github.com/almaan/stereoscope, version 0.3.1). We used the latest versions of all the software packages available at the time of this study. Unless otherwise stated, we used the default settings of each software package, including parameter specifications, quality control metrics of the ST data, and procedures for selecting genes used in the ST deconvolution.

Among the 5 existing methods, STdeconvolve and CARDfree are reference-free, while the other 3 methods require a single-cell gene expression reference for ST deconvolution. For each ST dataset analyzed in this study, we ran the 3 reference-based methods using the same single-cell expression reference, as described in the following sections. For STdeconvolve, we determined *L* in the same manner as RETROFIT, setting *L* = 2*K* . For CARDfree, *L* was determined based on the input marker gene list, as required by the algorithm.

Since STdeconvolve and CARDfree did not ensure a one-to-one mapping between deconvolved components and cell types post-annotation, we used Algorithms 2-3 in RETROFIT to annotate the deconvolution results of STdeconvolve and CARDfree, which ensured consistency in output format with RETROFIT and the reference-based methods. For each ST dataset, each method outputs an estimated proportion for each cell type at each spot, which can be compared with the cell-type proportion estimates 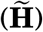 produced by RETROFIT.

STdeconvolve determined the optimal number of latent components as *L* = 7 in the mouse cerebellum Slide-seq study (*K* = 10). STdeconvolve determined the optimal number of latent components as *L* = 6 in the human intestine Visium study (*K* = 8).

### Simulation studies

Multiple factors in ST data may affect the performance of cell-type deconvolution. First, spot size differs across ST technologies and affects the complexity of the cell-type mixture at each spot. Second, cell-type heterogeneity in a ST slide also varies. A ST slide from a highly heterogeneous tissue (e.g., mammalian brains and intestines) tends to produce spots with multiple cell types. Methods that limit the number of cell types at a spot^15^ are likely inadequate for deconvolving ST data with high cell-type heterogeneity. Third, sequencing depths on the same slide may vary across spots, requiring methods to be adaptive and robust. Lastly, while RETROFIT does not require a single-cell gene expression reference for deconvolution, many existing methods do and thus their performance relies on the reference quality. We conducted simulations to investigate the impact of these factors on the performance of RETROFIT and several existing ST deconvolution methods.

To imitate ST experiments from various platforms and tissue samples, we simulate ST data with different spot sizes and cell-type heterogeneity levels. Specifically, we characterize spot size by the number of cells per spot (*N*) and cell-type heterogeneity by the maximum number of cell types per spot (*M*). For each combination of *N* and *M*, we simulate the ST data matrix of *G* genes and *S* spots as follows. For each spot *s*, we randomly select an integer *K*_*s*_ between 1 and *M*, and then randomly select *K*_*s*_ cell types from the *K* cell types present in the ST sample, denoted as 𝒦_*s*_. We simulate the proportions of the *K*_*s*_ selected cell types using a flat Dirichlet distribution, 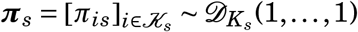, and obtain the cell counts for the *K*_*s*_ selected cell types at spot *s* as 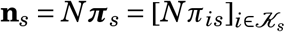, rounding to the nearest integer. We randomly select **n**_*s*_ unique cells from a single-cell gene expression reference of the *K*_*s*_ selected cell types and aggregate their single-cell expression profiles of *G* genes to produce the expression profile for spot *s*. For example, if a spot contains *N* = 10 cells from cell types a, b and c with proportions 0.1, 0.7 and 0.2, respectively, we randomly select 1, 7 and 2 unique cells from the corresponding single-cell expression reference of cell types a, b and c, and then add their gene expression profiles up as the aggregated expression profile for this spot. To incorporate sequencing depth variation across spots, we simulate a spot-specific effect *ϵ*_*s*_ for each spot *s* from a Gamma distribution, *ϵ*_*s*_ ∼ 𝒢 (3, 1), and multiply the aggregated expression level for each gene at spot *s* by *ϵ*_*s*_ to obtain the final ST expression level. The step-by-step protocol to generate synthetic ST data is given by Algorithm 4.

#### Algorithm 4

Synthetic ST data generation

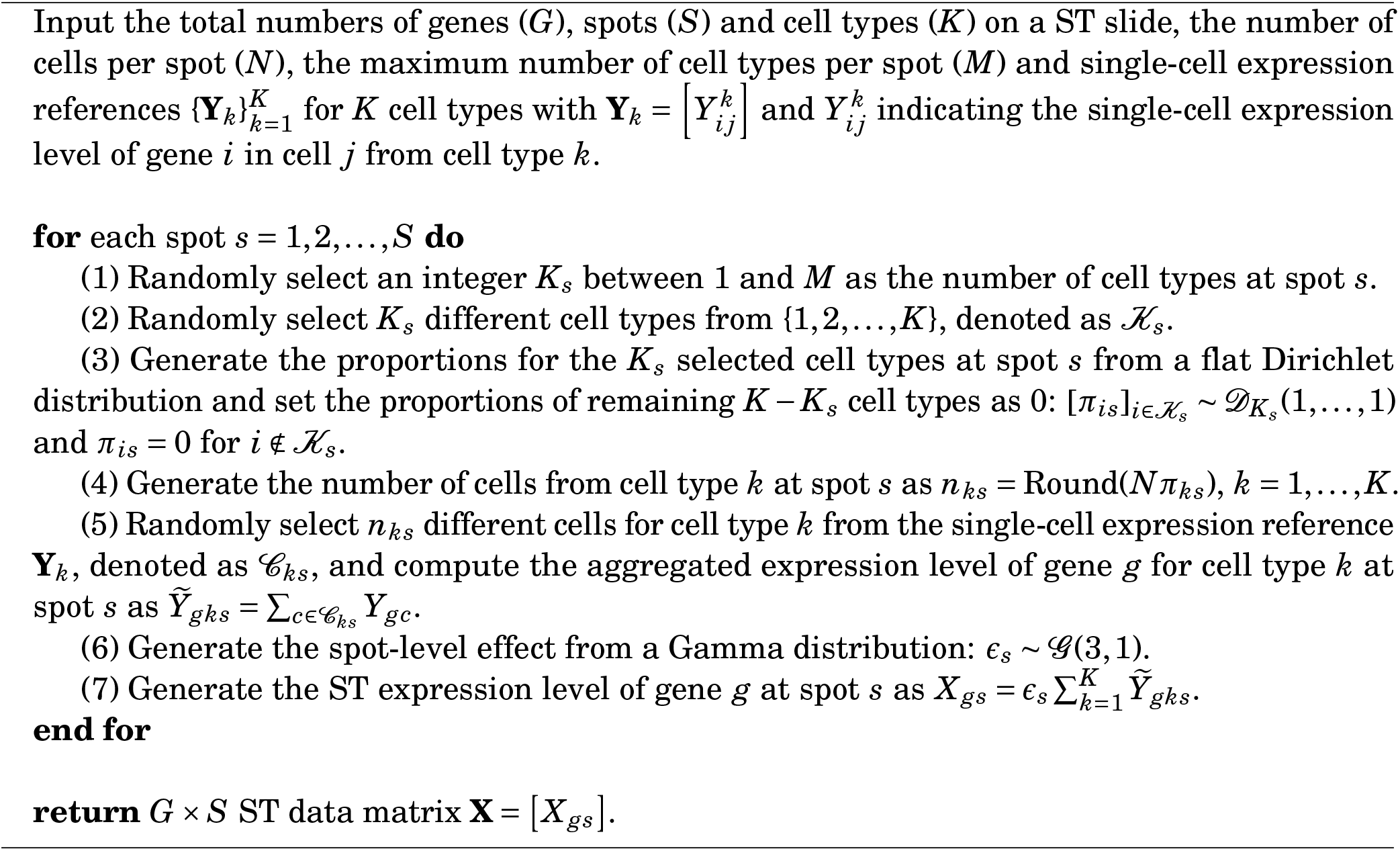

To simulate ST data for this study, we applied Algorithm 4 to a mouse cerebellum scRNA-seq dataset ^11^ of 2505 genes and 26139 cells for 10 annotated cell types; see the next section for more details on this dataset. We selected 30 cells from each of the 10 cell types in this scRNA-seq dataset, based on the highest sum of single-cell expression levels across the 2505 genes. Next, we identified the 500 genes with the most variability across the 300 selected cells and used them to simulate three ST datasets with *G* = 500 genes and *S* = 1000 spots: (1) *N* = 10 cells from up to *M* = 3 of the *K* = 10 cell types per spot (column 1 of Fig. 2); (2) *N* = 20 cells from up to *M* = 5 of the *K* = 10 cell types per spot (column 2 of Fig. 2); (3) *N* = 10 cells from up to *M* = 3 of the *K* = 5 ground-truth cell types per spot (columns 3-4 of Fig. 2) with the 5 ground-truth cell types being Bergmann glia, choroid plexus, endothelial, oligodendrocyte and Purkinje.

We used RETROFIT, STdeconvolve and CARDfree for reference-free deconvolution of each ST dataset with their default settings. When running RETROFIT and STdeconvolve, we set *L* = 20 for the first two ST datasets (*K* = 10) and *L* = 10 for the third ST dataset (*K* = 5). When running CARDfree, we used the *L* determined by the algorithm itself, as it did not allow users to specify *L*. To map deconvolved components to ground-truth cell types, we created a cell-type-specific transcriptomic reference 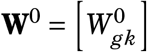, for the *G* genes and *K* ground-truth cell types in each ST dataset, using scRNA-seq data from the same sample^11^ hat produced the ST data. Specifically, we set 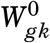 as the average scRNA-seq expression level of gene *g* across the 30 cells from cell type *k* that were used to simulate the ST data. We applied Algorithm 2 to this reference to annotate the results of RETROFIT, STdeconvolve and CARDfree.

To perform reference-based deconvolution in simulations, we applied Cell2location, RCTD and Stereoscope to each of the three simulated ST datasets using their default settings. For the first two ST datasets, we used the exact scRNA-seq data ^11^ of the 10 ground-truth cell types that were used to simulate the ST data as the single-cell gene expression reference (columns 1-2 of Fig. 2). For the third ST dataset, we created two ‘imperfect’ references for reference-based methods based on the same scRNA-seq dataset. Specifically, one reference contained 5 ground-truth and 5 irrelevant cell types (column 3 of Fig. 2), while the other reference contained only 3 of the 5 ground-truth cell types (absent: choroid plexus and oligodendrocyte) and 5 irrelevant cell types (column 4 of Fig. 2). When analyzing the third ST dataset with two ‘imperfect’ references, we used the non-default flag `-keep_noise` in Stereoscope to account for additional or missing cell types in the reference. When a ground-truth cell type was absent from the single-cell gene expression reference, all reference-based methods were unable to estimate its proportion at each spot, and thus we set the estimate as zero.

We evaluated the performance of RETROFIT and 5 existing methods on the synthetic ST data as follows. Given a ST dataset, each method produced a proportion estimate of each cell type *k* for each spot *s*: 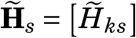. These estimates were used to reconstruct the ST expression profile for spot *s* as 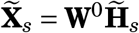 with **W** being the cell-type-specific expression reference of *G* genes for *K* cell types as described above. We compared 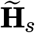 with the true cell-type proportions at the same spot, **H**_*s*_ = [*H*_*ks*_], by computing (1) their RMSE (Fig. 2a), defined as 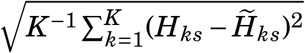, and(2) their Pearson correlation (Fig. 2b). Similarly, we compared 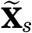 with the true ST expression profile at the same spot, **X**_*s*_ = [*X*_*gs*_], by computing (1) their normalized RMSE (Fig. 2c), defined as 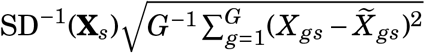 with SD(**X**_*s*_) being the standard deviation of ST expression levels across *G* genes at spot *s*, and (2) their correlation (Fig. 2d). For both estimated cell-type proportions 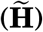 and reconstructed gene expression levels 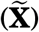, lower RMSEs and higher correlations indicate better cell-type deconvolution results that are closer to and more concordant with the ground truth, respectively. To evaluate the cell-type specificity of RETROFIT-extracted components, we computed the correlation between the estimated 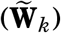 and observed 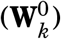 cell-type-specific expression levels across *G* genes for each cell type *k* (Fig. 2e), with a higher value indicating better performance.

### Mouse cerebellum data analysis

The mouse cerebellum study ^11^ provided Slide-seq data of 17919 genes at 27261 spots (https://singlecell.broadinstitute.org/single_cell/study/SCP354/slide-seq-study). This study also provided scRNA-seq data of 2505 genes from 26139 cells that were annotated as 10 cell types in the mouse cerebellum (astrocyte, Bergmann glia, choroid plexus, endothelial, granule, microglia, mural, oligodendrocyte, Purkinje and interneuron).

We used RETROFIT to analyze the mouse cerebellum Slide-seq data as follows. To ensure deconvolution accuracy and computation efficiency, we combined three complementary strategies to down-select genes before running RETROFIT. First, we selected 61 overdispersed genes with significantly higher-than-expected ST expression variances across spots ^28^ using the default setup of the STdeconvolve package. Second, we identified 54 cell-type-specific genes by computing entropy and Gini index on the companion scRNA-seq data (Supplementary Note 2). Third, we obtained 61 marker genes ^50^ curated for 3 mouse brain cell types (granule: 15; oligodendrocyte: 4; Purkinje: 42) from NeuroExpresso (www.neuroexpresso.org; Supplementary Table 2). We took the union of these 3 gene lists and used the resulting 153 unique genes to construct the input ST data matrix (**X**) for RETROFIT. We then ran RETROFIT on the 153 × 27261 ST data matrix **X** with *L* = 20 latent components. To map the RETROFIT-extracted components to the *K* = 10 mouse brain cell types, we applied Algorithm 2 to the cell-type-specific gene expression reference (**W**^0^) from the companion scRNA-seq data as described in the previous section.

We ran Cell2location, RCTD and Stereoscope on the Slide-seq and scRNA-seq data provided in this study. We ran STdeconvolve and CARDfree on the Slide-seq data only and used Algorithm 2 for annotation as in RETROFIT.

We evaluated the performance of 6 deconvolution methods on 3 mouse brain cell types (granule, oligodendrocyte, Purkinje) using curated marker genes available in NeuroExpresso^50^. Given a cell type *k*, we define the cell-type marker ST expression score at spot *s* in a slide as

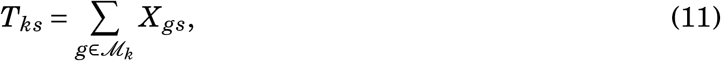

where ℳ_*k*_ denotes the list of marker genes for cell type *k* and *X*_*gs*_ denotes the ST expression level of gene *g* at spot *s*. For each combination of the 6 methods and the 3 cell types, we computed the Pearson correlation between the observed cell-type marker ST expression scores (**T**) and the estimated cell-type proportions 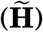 across all spots. A higher correlation indicates a better performance.

### Analysis of Visium HD colorectal cancer data

We analyzed the Visium HD dataset from patient 2 (P2) reported in ^33^, obtained from the Open Storage Framework (OSF) repository^51^, which provides both the spatial transcriptomics (ST) matrix (18,085 genes × 8,731,400 bins) and matched Chromium scRNA-seq data (18,082 genes × 67,568 cells) annotated into ten cell types.

Following 10x Genomics recommendations and the workflow of^51^, raw 2 *µ*m × 2 *µ*m bins were aggregated into 8 *µ*m × 8 *µ*m bins by merging each 2 × 2 block, yielding 545,913 spots. For computational tractability, we analyzed a contiguous 1/64 subregion (as in ^51^), resulting in 9,443 spots. After filtering spots with extremely low library size (<100 UMIs) or poor QC metrics (low log-library size, low feature count, or high mitochondrial fraction), 9,384 spots remained. We selected the top 3,000 highly variable genes and restricted analyses to genes and cell types shared across the ST and scRNA-seq assays, producing a final dataset of 3,000 genes × 8,880 spots. For RCTD, the scRNA-seq reference was downsampled to at most 4,000 cells per cell type.

RETROFIT was run with *L* = 10 components (seed = 929). STdeconvolve was run using its standard pipeline: overdispersed genes were selected (99 retained), spots with zero total counts were removed (8,747 remaining), and LDA models with 5–15 topics were fit. The best 10-topic model was selected. Proportions were mapped back to all 8,880 spots by assigning zeros to filtered-out entries. RCTD (spacexr v2.2.1) was run in *doublet* mode using the downsampled scRNA-seq reference. Cell2location (v0.1.5) was run using the matched scRNA-seq reference, with hyperparameters `N_cells_per_location=1` and `detection_alpha=20`, values commonly used for Visium-resolution data. Component annotations for RETROFIT and STdeconvolve were assigned via Pearson correlation with scRNA-seq cell-type profiles (Algorithm 2). Correlation-based evaluations were conducted on the 6,559 spots shared across all methods.

CARDfree was not included in this analysis because it requires at least five marker genes per cell type, whereas the Visium HD dataset provides only one or two validated markers per cell type and therefore does not meet its minimum input requirements.

### Human intestine data analysis

The human intestine study^10^ made available Visium ST data from 3 tissue slides, including a 12 PCW slide with 1080 spots, a 19 PCW slide with 1242 spots and an adult slide with 2649 spots, providing expression measurements for 33538 genes in each slide (https://www.ncbi.nlm.nih.gov/geo/query/acc.cgi?acc=GSE158328). The study provided H&E images of the ST slides (https://doi.org/10.17632/gncg57p5×9.2). This study also provided scRNA-seq data of 76592 cells from 77 intestinal samples spanning 8 to 22 PCW (https://www.ncbi.nlm.nih.gov/geo/query/acc.cgi?acc=GSE158702) that were grouped into 8 distinct cellular compartments (endothelial, epithelial, fibroblast, immune, muscle, MyoFB/MESO, neural, pericyte).

We used RETROFIT to analyze the human intestine Visium ST data as follows. Similar to our analysis of the mouse cerebellum Slide-seq data, we down-selected genes prior to running RETROFIT. Specifically, for each of the 3 ST slides, we only included (1) significantly overdispersed genes across spots^28^ and (2) 37 known marker genes of the 8 cellular compartments (Fig. 7a) in the input ST data matrix (**X**) for RETROFIT. This resulted in 722 genes for 12 PCW, 681 genes for 19 PCW, and 1051 genes for adult. RETROFIT was then run on each ST data matrix with *L* = 16 latent components. To match the RETROFIT-extracted components with the *K* = 8 intestinal compartments, we utilized Algorithm 3 alongside the 37 curated marker genes (Fig. 7a) from the human intestine study^10^. Additionally, for the 12 and 19 PCW slides, which have the corresponding scRNA-seq data, we also applied Algorithm 2 to annotate RETROFIT results, using the compartment-specific gene expression reference (**W**^0^) generated from the companion scRNA-seq data. Specifically, for each fetal stage, we selected 25 cells from each compartment that had the highest sum of single-cell expression levels across all genes. We then define 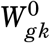 as the average scRNA-seq expression level of gene *g* across these selected 25 cells from compartment *k*. Unless specified otherwise, estimates of compartment proportion 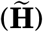 and compartment-specific expression 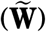 for all 3 ST slides were generated using Algorithm 3.

To benchmark RETROFIT’s performance, we also evaluated STdeconvolve, CARDfree and Cell2location using the same human intestine ST data. Given that Cell2location requires a single-cell gene expression reference, our comparison was limited to the ST data from 12 and 19 PCW tissue slides, which had companion scRNA-seq data. The execution details for these external methods mirror those outlined in previous sections.

To evaluate the accuracy of RETROFIT in estimating cellular compartment proportions 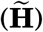, we computed the correlation between the ST expression scores of compartment-specific marker genes defined in Eq. (11) and the estimated compartment proportions across all spots for each of the 8 cellular compartments and 3 ST slides (Table **??**; Fig.s 5c-d; Supplementary Fig.s 7-12).

To evaluate the accuracy of RETROFIT in estimating compartment-specific expression levels 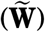, we compared the compartment-specific expression levels estimated from 12 and 19 PCW ST slides with the compartment-specific expression levels based on the companion scRNA-seq data from 12 and 19 PCW intestinal samples (**W**^0^). To account for different scales of ST and scRNA-seq data, we first normalized rows of the two expression matrices (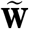 and **W**^0^) by their sums as we did in Algorithm 2:

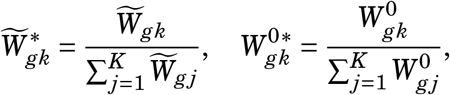

and then compared the normalized expression matrices 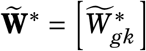 and 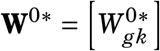 in each of the 8 cellular compartments and 2 fetal stages (Fig.s 7a and c).

Based on the normalized cell-type-specific expression levels 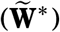 estimated by RETROFIT, we further developed a simple method to identify genes with high cell-type specificity. Given the normalized cell-type-specific expression estimates of gene *g* for *K* cell types 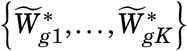, we calculated two dispersion measures:

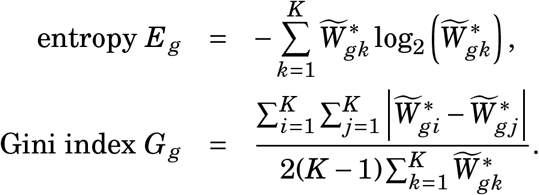

Lower entropy and higher Gini index indicate an excess of normalized expression for one cell type, thus suggesting the cell-type specificity. In the human intestine data analysis, we identified a cell-type-specific gene *g* from the ST data if this gene had (1) entropy *E*_*g*_ < 1.5, (2) Gini index *G*_*g*_ > 0.85, (3) maximum ST expression level max_*s*_ *X*_*gs*_ > 40 and (4) consistent cell-type specificity across all ST replicate samples (e.g., the same tissue type from the same developmental stage). We performed this analysis only on adult and 12 PCW stages (Fig. 7b), because they were the only stages with ST replicate samples available in the human intestine study ^10^ (Supplementary Tables 22-25), in addition to the ST samples used in our primary analysis (Fig. 5a).

To assess biological themes of cell-type-specific genes identified by RETROFIT (Fig. 7d; Supplementary Tables 26-27), we performed the gene set enrichment analysis using Metascape ^52^ (https://metascape.org, version 3.5). Metascape calculates the enrichment *P*-values based on the cumulative hypergeometric distribution and then adjusts the *P*-values for multiple testing based on the Benjamini-Hochberg procedure.

### Computational time and memory usage

The total computational time and memory usage of RETROFIT to deconvolve a ST slide is determined by the numbers of genes (*G*) and spots (*S*) analyzed, the number of latent components (*L*), and the number of SSVI iterations (*I*), all of which can vary considerably among studies. It is thus hard to make general statements about computation time and memory usage. However, to give a specific example, it took RETROFIT 39.92 minutes and 1.76 gigabytes to complete the analysis of mouse cerebellum Slide-seq dataset (the largest dataset analyzed in this study) on a single CPU from the Intel Xeon Gold 6354 Processor. For comparison, we recorded the computational time and memory usage associated with running existing methods on the same Slide-seq dataset in the same computing environment (Supplementary Table 29). The results show that RETROFIT is efficient in terms of time and memory compared to existing methods.

## Data availability

All the data used in this study are publicly available. Links and identifiers of all data are specified in Methods. Source data are provided with this paper. The simulation ST data generated in this study has been deposited in Zenodo and is accessible via DOI: 10.5281/zenodo.19241634^53^. The processed Mouse cerebellum Slide-seq data and processed Human intestine data have been deposited in Zenodo and is accessible via DOI: 10.5281/zenodo.19241634^53^. The decomposed results for these ST data generated in this study are provided in the Supplementary Information/Source Data file.

## Code availability

RETROFIT is available as an R package in Bioconductor (https://bioconductor.org/packages/retrofit/). The code used to develop the model, perform the analyses and generate results in this study is publicly available and has been deposited in GitHub at https://github.com/qunhualilab/retrofit-scripts, under MIT license. The specific version of the code associated with this publication is archived in Zenodo and is accessible via DOI: 10.5281/zenodo.19241634^53^.

## Acknowledgments

This work was supported by the National Institutes of Health (NIH) under grants R01GM109453, R21AI160138, and R03DE031361 (Q.L.), and R24DK106766 (R.C.H.). X.Z. and Q.L. were supported by seed grants from the Institute for Computational and Data Sciences, the Consortium on Substance Use and Addiction, the CTSI Cores Pilot Project, and the CTSI Bridges to Translation VIII program at The Pennsylvania State University. This study used computational resources provided by the Institute for Computational and Data Sciences at The Pennsylvania State University.

## Author contributions

X.Z. and Q.L. conceived and supervised the study. R.S., X.Z. and Q.L. developed the methods. R.S., X.H., X.W. A.K.P. and X.Z. performed data analysis and experiments. A.K.P. developed the R package with input from R.S., X.Z. and Q.L. R.C.H. provided biological consultation. R.S., X.H., X.W., X.Z. and Q.L. wrote the manuscript with input from all authors.

## Tables

**Table 1.** Comparison of RETROFIT and other deconvolution methods on human fetal intestine Visium data. Pearson correlations between ST expression scores of known marker genes and estimated proportions across spots in both fetal slides are reported for all methods and cellular compartments. For each combination of slide and compartment, the largest correlation across methods is highlighted in bold. The latent components extracted by the three reference-free methods (RETROFIT, STdeconvolve, CARDfree) were mapped to known cellular compartments using the same strategy based on a curated list of 37 intestinal compartment marker genes (Algorithm 3). Cell2location, the only reference-based method, was provided with the companion scRNA-seq data in fetal intestinal samples from the same study. Bold values indicate the highest correlation for each combination of slide and cellular compartment.

| Compartment | 12 PCW ST slide |  |  |  | 19 PCW ST slide |  |  |  |
| --- | --- | --- | --- | --- | --- | --- | --- | --- |
|  | RETROFIT | STdeconvolve | CARDfree | Cell2location | RETROFIT | STdeconvolve | CARDfree | Cell2location |
| Endothelial | **0.68** | 0.42 | 0.36 | 0.62 | **0.54** | 0.21 | 0.12 | 0.47 |
| Epithelial | 0.55 | 0.51 | 0.47 | **0.77** | **0.63** | 0.60 | 0.19 | 0.51 |
| Fibroblast | **0.68** | 0.59 | 0.63 | 0.50 | 0.52 | 0.35 | **0.79** | 0.61 |
| Immune | 0.10 | $-8.4 \times 10^{-2}$ | **0.28** | 0.20 | -0.80 | $-7.5 \times 10^{-2}$ | -0.14 | **$7.8 \times 10^{-2}$** |
| Muscle | **0.87** | -0.20 | 0.72 | 0.83 | 0.64 | 0.53 | 0.72 | **0.73** |
| MyoFB/MESO | **0.34** | $-1.2 \times 10^{-2}$ | 0.32 | $8.6 \times 10^{-2}$ | 0.48 | 0.48 | **0.70** | -0.33 |
| Neural | 0.69 | -0.12 | $-9.9 \times 10^{-2}$ | **0.75** | **0.60** | 0.62 | $9.3 \times 10^{-2}$ | 0.70 |
| Pericyte | 0.11 | $-3.5 \times 10^{-4}$ | $9.8 \times 10^{-2}$ | **0.13** | **0.54** | 0.24 | -0.13 | 0.13 |

