## Supplementary Information for "RETROFIT: Reference-free deconvolution of cell-type mixtures in spatial transcriptomics"

#### Supplementary Note 1

Here we show that Algorithm 1 in the main text is a direct application of structured stochastic variational inference[1] (SSVI) to the Bayesian hierarchical model underlying RETROFIT. We first review the SSVI algorithm and its assumptions, and then we establish the correspondence between SSVI and Algorithm 1 in the main text step by step. Most of the mathematical notations used here have been defined in the main text. Any new notation introduced here is defined at its first usage. "Equation" and "constant" are abbreviated to "Eq." and "Const." throughout this note, respectively.

##### Review of SSVI algorithm

SSVI makes two modeling assumptions. First, SSVI assumes a probability model of the following form:

$$p(y,u,\mu) = p(\mu)\prod_n p(y_n,u_n \mid \mu) \tag{1}$$

where $y = \{y_n\}$ denotes observed data, $u = \{u_n\}$ denotes local hidden variables and $\mu$ denotes global parameters. Eq. (1) indicates that global parameters $\mu$ are shared across all observations while local hidden variables $u$ are conditionally independent of one another given $\mu$. Second, SSVI further assumes that the prior $p(\mu)$ comes from an exponential family and $p(\mu)$ is conditionally conjugate for the complete-data likelihood $p(y_n,u_n \mid \mu)$. Specifically, the conjugate pair of $\{p(\mu), p(y_n,u_n \mid \mu)\}$ is given by:

$$\log p(\mu) = \eta \cdot T(\mu) + B(\mu) + \text{Const. of } \mu, \tag{2}$$

$$\log p(y_n,u_n \mid \mu) = T(\mu)\cdot \eta_n(y_n,u_n) + \text{Const. of } \mu, \tag{3}$$

where $\eta$ denotes natural parameters in the prior $p(\mu)$ and $\{T,B,\eta_n\}$ are known functions. To verify that $\{p(\mu), p(y_n,u_n \mid \mu)\}$ in Eq.s (2)-(3) form a conditionally conjugate model, we can show that the conditional posterior $p(\mu \mid y,u)$ is in the same exponential family as the prior $p(\mu)$, that is,

$$\begin{aligned}\log p(\mu \mid y,u) &= \log p(\mu) + \sum_n \log p(y_n,u_n \mid \mu) + \text{Const. of } \mu \\ &= \left[\eta + \sum_n \eta_n(y_n,u_n)\right]\cdot T(\mu) + B(\mu) + \text{Const. of } \mu.\end{aligned} \tag{4}$$

RETROFIT satisfies these modeling assumptions of SSVI, as shown in the following sections.

SSVI makes three approximating assumptions. First, SSVI assumes that the full posterior distribution $p(u,\mu \mid y)$ is approximated by a variational distribution of the following form:

$$q(u,\mu) = q(\mu)\prod_n q(u_n \mid \mu). \tag{5}$$

Second, SSVI assumes that $q(\mu)$ in Eq. (5) is in the same exponential family as the prior $p(\mu)$:

$$\log q(\mu) = \phi \cdot T(\mu) + B(\mu) + \text{Const. of } \mu, \tag{6}$$

where $\phi$ denotes natural parameters that control the variational distribution $q(\mu)$. Third, SSVI assumes that any dependence between $u_n$ and $\mu$ under the variational distribution $q$ is solely mediated by some function $\gamma_n(\mu)$ such that $q(u_n \mid \mu)$ in Eq. (5) can be rewritten as

$$q(u_n \mid \mu) = q\left[u_n \mid \gamma_n(\mu)\right]. \tag{7}$$

RETROFIT also satisfies these approximating assumptions, as explained in the following sections.

With all the assumptions in place, SSVI is implemented as the following algorithm.

- Initialize $\phi(0)$ randomly.
- For iteration $i = 1, 2, \ldots, I$,

Step 1 Sample global parameters $\mu(i)$ from $q(\mu)$ while setting $\phi = \phi(i-1)$ in Eq. (6);

Step 2 Compute the local variational parameters $\gamma_n\left[\mu(i)\right]$ in Eq. (7) such that the Kullback-Leibler (KL) divergence between $q\left[u_n \mid \mu(i)\right]$ and $p(u_n \mid y_n, \mu)$ is minimized;

Step 3 Update $\phi$ in Eq. (6) to $\phi(i)$ with the following stochastic natural gradient of step size $\rho(i)$:

$$\phi(i) = [1-\rho(i)]\cdot\phi(i-1) + \rho(i)\cdot\left[\eta + \sum_n \hat{\eta}_n(i)\right], \tag{8}$$

where $\eta$ denotes natural parameters in the prior (2), $\hat{\eta}_n(i)$ is an unbiased estimate of $\mathrm{E}_q\left[\eta_n(y_n, u_n) \mid \mu(i)\right]$ under the optimal local variational distribution $q\left[u_n \mid \mu(i)\right]$ obtained in Step 2, and $\eta_n(y_n, u_n)$ is defined in the complete-data likelihood (3).

Upon convergence, the SSVI algorithm approximates the intractable posterior distribution $p(u, \mu \mid y)$ with the variational distribution $q(u, \mu)$ of the form (5) whose parameters are estimated through the iterative and stochastic procedure above.

Algorithm 1 in the main text is a direct application of the SSVI algorithm to the Bayesian hierarchical model underlying RETROFIT, where $\mathbf{X}$ corresponds to observed data "$y$", $\mathbf{Z} = \left\{\mathbf{Z}^0_{gs}, \mathbf{Z}^1_{gs}\right\}$ corresponds to local hidden variables "$u$" and $\{\mathbf{W}, \boldsymbol{\theta}, \mathbf{H}\}$ correspond to global parameters "$\mu$". According to the assumption (7) of SSVI, the variational approximation for the full posterior distribution $p(\mathbf{Z}, \mathbf{W}, \boldsymbol{\theta}, \mathbf{H} \mid \mathbf{X})$ is given by

$$q(\mathbf{Z}, \mathbf{W}, \boldsymbol{\theta}, \mathbf{H}) = q(\mathbf{W}, \boldsymbol{\theta}, \mathbf{H})q(\mathbf{Z} \mid \mathbf{W}, \boldsymbol{\theta}, \mathbf{H}) = \prod_{g,\ell} q(W_{g\ell}) \prod_{\ell} q(\theta_\ell) \prod_{\ell,s} q(H_{\ell s}) \prod_{g,s} q\left(\mathbf{Z}^0_{gs}, \mathbf{Z}^1_{gs} \mid \mathbf{W}, \boldsymbol{\theta}, \mathbf{H}\right). \tag{9}$$

In the following sections, we describe how to implement Steps 1-3 of each iteration in SSVI to find $q(\mathbf{Z}, \mathbf{W}, \boldsymbol{\theta}, \mathbf{H})$.

#### Sampling of global parameters $\{\mathbf{W}, \boldsymbol{\theta}, \mathbf{H}\}$

In RETROFIT, we place the following independent Gamma priors on the global parameters $\{\mathbf{W}, \boldsymbol{\theta}, \mathbf{H}\}$:

$$W_{g\ell} \overset{\text{i.i.d.}}{\sim} \mathcal{G}\left(\alpha^W_0, \beta^W_0\right), \quad \theta_\ell \overset{\text{i.i.d.}}{\sim} \mathcal{G}\left(\alpha^\theta_0, \beta^\theta_0\right), \quad H_{\ell s} \overset{\text{i.i.d.}}{\sim} \mathcal{G}\left(\alpha^H_0, \beta^H_0\right), \tag{10}$$

where $\left\{\alpha^W_0, \beta^W_0, \alpha^\theta_0, \beta^\theta_0, \alpha^H_0, \beta^H_0\right\}$ are known hyper-parameters. Because Gamma distributions belong to the exponential family, the assumption (2) of SSVI is satisfied. According to the assumption (6) of SSVI, the variational distributions for $\{\mathbf{W}, \boldsymbol{\theta}, \mathbf{H}\}$ must be in the same exponential family as their prior distributions (10):

$$q(W_{g\ell}) = \mathcal{G}\left(W_{g\ell}; \alpha^W_{g\ell}, \beta^W_{g\ell}\right), \quad q(\theta_\ell) = \mathcal{G}\left(\theta_\ell; \alpha^\theta_\ell, \beta^\theta_\ell\right), \quad q(H_{\ell s}) = \mathcal{G}\left(H_{\ell s}; \alpha^H_{\ell s}, \beta^H_{\ell s}\right), \tag{11}$$

where the variational parameters $\left\{\alpha^W_{g\ell}, \beta^W_{g\ell}, \alpha^\theta_\ell, \beta^\theta_\ell, \alpha^H_{\ell s}, \beta^H_{\ell s}\right\}$ are estimated by SSVI.

To implement Step 1 of iteration $i$ in SSVI, we sample the global parameters $\{\mathbf{W}, \boldsymbol{\theta}, \mathbf{H}\}$ as

$$W_{g\ell}(i) \sim \mathcal{G}\left(\alpha^W_{g\ell}(i-1), \beta^W_{g\ell}(i-1)\right), \quad \theta_\ell(i) \sim \mathcal{G}\left(\alpha^\theta_\ell(i-1), \beta^\theta_\ell(i-1)\right), \quad H_{\ell s}(i) \sim \mathcal{G}\left(\alpha^H_{\ell s}(i-1), \beta^H_{\ell s}(i-1)\right), \tag{12}$$

where $\left\{\alpha^W_{g\ell}(i-1), \beta^W_{g\ell}(i-1), \alpha^\theta_\ell(i-1), \beta^\theta_\ell(i-1), \alpha^H_{\ell s}(i-1), \beta^H_{\ell s}(i-1)\right\}$ are the variational parameter values obtained from Step 3 of iteration $i-1$. The sampling scheme defined in (12) corresponds to the first step of each iteration in Algorithm 1 in the main text.

##### Optimization of local variational distribution $q(\mathbf{Z} \mid \mathbf{W}, \boldsymbol{\theta}, \mathbf{H})$

In RETROFIT, we specify the local variational distribution $q\left(\mathbf{Z}_{gs}^0, \mathbf{Z}_{gs}^1 \mid \mathbf{W}, \boldsymbol{\theta}, \mathbf{H}\right)$ as the exact conditional posterior distribution of $\left\{\mathbf{Z}_{gs}^0, \mathbf{Z}_{gs}^1\right\}$ given $\{\mathbf{W}, \boldsymbol{\theta}, \mathbf{H}\}$:

$$q\left(\mathbf{Z}_{gs}^0, \mathbf{Z}_{gs}^1 \mid \mathbf{W}, \boldsymbol{\theta}, \mathbf{H}\right) = p\left(\mathbf{Z}_{gs}^0, \mathbf{Z}_{gs}^1 \mid \mathbf{X}, \mathbf{W}, \boldsymbol{\theta}, \mathbf{H}\right), \tag{13}$$

which provides the best possible approximation by achieving zero KL divergence to the actual conditional posterior[1]. With this specification, SSVI integrates out the latent part $\mathbf{Z}$ and minimizes the KL divergence between the marginal variational distribution $q(\mathbf{W}, \boldsymbol{\theta}, \mathbf{H})$ and the marginal posterior distribution $p(\mathbf{W}, \boldsymbol{\theta}, \mathbf{H} \mid \mathbf{X})$.

Another reason for using Eq. (13) is that $p\left(\mathbf{Z}_{gs}^0, \mathbf{Z}_{gs}^1 \mid \mathbf{X}, \mathbf{W}, \boldsymbol{\theta}, \mathbf{H}\right)$ has a closed form of a multinomial distribution. To see this, we first review an important relationship between Poisson and multinomial distributions. If $P_1, \ldots, P_m$ are independent Poisson random variables with rate parameters $r_1, \ldots, r_m$, then $(P_1, \ldots, P_m)$ given $\sum_{i=1}^m P_i = M$ follows a multinomial distribution of size $M$ and probabilites $r_j / \sum_{i=1}^m r_i$, $j = 1, \ldots, m$. Next, we use this relationship to find the analytic form of $p\left(\mathbf{Z}_{gs}^0, \mathbf{Z}_{gs}^1 \mid \mathbf{X}, \mathbf{W}, \boldsymbol{\theta}, \mathbf{H}\right)$. In RETROFIT, $\left\{Z_{g\ell s}^0, Z_{g\ell s}^1\right\}$ are independent Poisson random variables constituting the ST expression profile $X_{gs}$:

$$Z_{g\ell s}^0 \sim \mathscr{P}(\lambda H_{\ell s}), \quad Z_{g\ell s}^1 \sim \mathscr{P}(W_{g\ell}\theta_\ell H_{\ell s}), \quad \sum_{\ell=1}^L \left(Z_{g\ell s}^0 + Z_{g\ell s}^1\right) = X_{gs}. \tag{14}$$

Consequently, the right-hand side of Eq. (13) has the following closed form of a multinomial distribution:

$$\Pr\left(\mathbf{Z}_{gs}^0 = \mathbf{z}_{gs}^0, \mathbf{Z}_{gs}^1 = \mathbf{z}_{gs}^1 \mid \mathbf{X} = \mathbf{x}, \mathbf{W}, \boldsymbol{\theta}, \mathbf{H}\right) = \binom{x_{gs}}{z_{g1s}^0, \cdots, z_{gLs}^0, z_{g1s}^1, \cdots, z_{gLs}^1} \prod_{\ell=1}^L \left(\pi_{g\ell s}^0\right)^{z_{g\ell s}^0} \prod_{\ell=1}^L \left(\pi_{g\ell s}^1\right)^{z_{g\ell s}^1}, \tag{15}$$

where the multinomial probabilities are given by

$$\pi_{g\ell s}^0 = \frac{\lambda H_{\ell s}}{\sum_\ell (W_{g\ell}\theta_\ell + \lambda) H_{\ell s}}, \quad \pi_{g\ell s}^1 = \frac{W_{g\ell}\theta_\ell H_{\ell s}}{\sum_\ell (W_{g\ell}\theta_\ell + \lambda) H_{\ell s}}, \quad \ell = 1, \ldots, L. \tag{16}$$

Note that the dependence under $q\left(\mathbf{Z}_{gs}^0, \mathbf{Z}_{gs}^1 \mid \mathbf{W}, \boldsymbol{\theta}, \mathbf{H}\right)$ between $\left\{\mathbf{Z}_{gs}^0, \mathbf{Z}_{gs}^1\right\}$ and $\{\mathbf{W}, \boldsymbol{\theta}, \mathbf{H}\}$ is solely mediated through $\left\{\pi_{g\ell s}^0(\mathbf{W}, \boldsymbol{\theta}, \mathbf{H}), \pi_{g\ell s}^1(\mathbf{W}, \boldsymbol{\theta}, \mathbf{H})\right\}$ defined in Eq. (16), which also satisfies the assumption (7) of SSVI.

To implement Step 2 of iteration $i$ in SSVI, we update the multinomial probabilities from Eq. (16) that optimize the local variational distribution $q\left[\mathbf{Z}_{gs}^0, \mathbf{Z}_{gs}^1 \mid \mathbf{W}(i), \boldsymbol{\theta}(i), \mathbf{H}(i)\right]$:

$$\pi_{g\ell s}^0(i) = \frac{\lambda H_{\ell s}(i)}{\sum_\ell [W_{g\ell}(i)\theta_\ell(i) + \lambda] H_{\ell s}(i)}, \quad \pi_{g\ell s}^1(i) = \frac{W_{g\ell}(i)\theta_\ell(i) H_{\ell s}(i)}{\sum_\ell [W_{g\ell}(i)\theta_\ell(i) + \lambda] H_{\ell s}(i)}, \tag{17}$$

where $\{W_{g\ell}(i), \theta_\ell(i), H_{\ell s}(i)\}$ are the global parameter values sampled from Step 1 of iteration $i$. Eq. (17) corresponds to the second step of each iteration in Algorithm 1 in the main text.

##### Stochastic gradient update of variational parameters for W

Before deriving the stochastic gradient update of variational parameters for $\mathbf{W}$ in Eq. (11), we need to verify the conditional conjugacy assumption (4) of SSVI. We write the log of complete-data likelihood for $\{\mathbf{X},\mathbf{Z}\}$ as

$$\begin{aligned}
\log p(\mathbf{X},\mathbf{Z} \mid \mathbf{W},\boldsymbol{\theta},\mathbf{H}) &= \sum_{g,s} \log p\left(X_{gs},\mathbf{Z}^0_{gs},\mathbf{Z}^1_{gs} \mid \mathbf{W},\boldsymbol{\theta},\mathbf{H}\right) \\
&= \sum_{g,s} \left[\log p\left(X_{gs} \mid \mathbf{Z}^0_{gs},\mathbf{Z}^1_{gs}\right) + \log p\left(\mathbf{Z}^0_{gs} \mid \mathbf{W},\boldsymbol{\theta},\mathbf{H}\right) + \log p\left(\mathbf{Z}^1_{gs} \mid \mathbf{W},\boldsymbol{\theta},\mathbf{H}\right)\right] \\
&= \sum_{g,\ell,s} \left[\log p\left(Z^0_{g\ell s} \mid \mathbf{W},\boldsymbol{\theta},\mathbf{H}\right) + \log p\left(Z^1_{g\ell s} \mid \mathbf{W},\boldsymbol{\theta},\mathbf{H}\right)\right] + \text{Const. of } \{\mathbf{W},\boldsymbol{\theta},\mathbf{H}\} \\
&= \sum_{g,\ell,s} \left[Z^1_{g\ell s}\cdot \log W_{g\ell} + Z^1_{g\ell s}\cdot\log\theta_\ell + \left(Z^0_{g\ell s}+Z^1_{g\ell s}\right)\cdot\log H_{\ell s} - (W_{g\ell}\theta_\ell+\lambda)H_{\ell s}\right] \qquad (18) \\
&\quad + \text{Const. of } \{\mathbf{W},\boldsymbol{\theta},\mathbf{H}\},
\end{aligned}$$

where the last equation holds true because $Z^0_{g\ell s} \sim \mathscr{P}(\lambda H_{\ell s})$ and $Z^1_{g\ell s} \sim \mathscr{P}(W_{g\ell}\theta_\ell H_{\ell s})$ independently. We write the log of Gamma prior density for $\mathbf{W}$ as

$$\log p(\mathbf{W}) = \sum_{g,\ell}\left[\left(\alpha^W_0 - 1\right)\cdot\log W_{g\ell} - \beta^W_0\cdot W_{g\ell}\right] + \text{Const. of } \mathbf{W}. \qquad (19)$$

We then combine Eq.s (18)-(19) to derive the log of conditional posterior distribution for $\mathbf{W}$ given $\{\boldsymbol{\theta},\mathbf{H}\}$ as

$$\begin{aligned}
\log p(\mathbf{W} \mid \mathbf{X},\mathbf{Z},\boldsymbol{\theta},\mathbf{H}) &= \log p(\mathbf{X},\mathbf{Z} \mid \mathbf{W},\boldsymbol{\theta},\mathbf{H}) + \log p(\mathbf{W} \mid \boldsymbol{\theta},\mathbf{H}) + \text{Const. of } \mathbf{W} \\
&= \log p(\mathbf{X},\mathbf{Z} \mid \mathbf{W},\boldsymbol{\theta},\mathbf{H}) + \log p(\mathbf{W}) + \text{Const. of } \mathbf{W} \\
&= \sum_{g,\ell}\left[\left(\alpha^W_0 + \sum_s Z^1_{g\ell s} - 1\right)\cdot\log W_{g\ell} - \left(\beta^W_0 + \sum_s \theta_\ell H_{\ell s}\right)\cdot W_{g\ell}\right] + \text{Const. of } \mathbf{W}, \qquad (20)
\end{aligned}$$

where the first equation is a direct application of Bayes' Theorem and the second equation holds true because $\{\mathbf{W},\boldsymbol{\theta},\mathbf{H}\}$ are mutually independent a priori. By comparing Eq.s (19)-(20) we can see that the conditional posterior $p(\mathbf{W} \mid \mathbf{X},\mathbf{Z},\boldsymbol{\theta},\mathbf{H})$ and the prior $p(\mathbf{W})$ come from the same Gamma family, and thus we verify the conditional conjugacy assumption (4) of SSVI for $\mathbf{W}$.

Eq. (20) further allows us to construct the SSVI stochastic gradient update (8) for variational parameters $\left\{\alpha^W_{g\ell},\beta^W_{g\ell}\right\}$ in Eq. (11). Specifically, we first replace the unobserved variable $Z^1_{g\ell s}$ in Eq. (20) with its unbiased estimate $\mathrm{E}_q\left(Z^1_{g\ell s} \mid \mathbf{W},\boldsymbol{\theta},\mathbf{H}\right) = X_{gs}\pi^1_{g\ell s}$, where $q$ denotes the optimal local variational distribution given by Eq. (15), and then we use $\alpha^W_0 + \sum_s X_{gs}\pi^1_{g\ell s}$ and $\beta^W_0 + \sum_s \theta_\ell H_{\ell s}$ as the "$\eta + \sum_n \hat{\eta}_n$" term in the stochastic gradient update (8) for $\alpha^W_{g\ell}$ and $\beta^W_{g\ell}$, respectively.

To implement Step 3 of iteration $i$ in SSVI for $W_{g\ell}$, we update its variational parameters $\left\{\alpha^W_{g\ell},\beta^W_{g\ell}\right\}$ as

$$\begin{aligned}
\alpha^W_{g\ell}(i) &= [1-\rho(i)]\cdot\alpha^W_{g\ell}(i-1) + \rho(i)\cdot\left[\alpha^W_0 + \sum_s X_{gs}\pi^1_{g\ell s}(i)\right], \qquad (21) \\
\beta^W_{g\ell}(i) &= [1-\rho(i)]\cdot\beta^W_{g\ell}(i-1) + \rho(i)\cdot\left[\beta^W_0 + \sum_s \theta_\ell(i)H_{\ell s}(i)\right], \qquad (22)
\end{aligned}$$

where $\left\{\alpha^W_{g\ell}(i-1),\beta^W_{g\ell}(i-1)\right\}$ are the values of $\left\{\alpha^W_{g\ell},\beta^W_{g\ell}\right\}$ from Step 3 of iteration $i-1$ and $\pi^1_{g\ell s}(i)$ is the local variational parameter value from Step 2 of iteration $i$. Eq.s (21)-(22) correspond to the first two updates in the third step of each iteration in Algorithm 1 in the main text.

##### Stochastic gradient update of variational parameters for $\boldsymbol{\theta}$

We use the same argument as above to verify the conditional conjugacy assumption (4) of SSVI for $\boldsymbol{\theta}$:

$$\begin{aligned}\log p(\boldsymbol{\theta}) &= \sum_{\ell}\left[\left(\alpha_0^{\theta}-1\right)\cdot\log\theta_{\ell}-\beta_0^{\theta}\cdot\theta_{\ell}\right]+\text{Const. of }\boldsymbol{\theta}, &(23)\\ \log p(\boldsymbol{\theta}\,|\,\mathbf{X},\mathbf{Z},\mathbf{W},\mathbf{H}) &= \log p(\mathbf{X},\mathbf{Z}\,|\,\mathbf{W},\boldsymbol{\theta},\mathbf{H})+\log p(\boldsymbol{\theta})+\text{Const. of }\boldsymbol{\theta}\\ &= \sum_{\ell}\left[\left(\alpha_0^{\theta}+\sum_{g,s}Z_{g\ell s}^{1}-1\right)\cdot\log\theta_{\ell}-\left(\beta_0^{\theta}+\sum_{g,s}W_{g\ell}H_{\ell s}\right)\cdot\theta_{\ell}\right]+\text{Const. of }\boldsymbol{\theta}. &(24)\end{aligned}$$

We then replace the unobserved variable $Z_{g\ell s}^1$ in Eq. (24) with $\mathrm{E}_q\left(Z_{g\ell s}^1\,|\,\mathbf{W},\boldsymbol{\theta},\mathbf{H}\right)=X_{gs}\pi_{g\ell s}^1$ to construct the SSVI stochastic gradient update (8) for variational parameters $\{\alpha_\ell^\theta,\beta_\ell^\theta\}$ in Eq. (11).

To implement Step 3 of iteration $i$ in SSVI for $\theta_\ell$, we update its variational parameters $\{\alpha_\ell^\theta,\beta_\ell^\theta\}$ as

$$\begin{aligned}\alpha_\ell^\theta(i) &= [1-\rho(i)]\cdot\alpha_\ell^\theta(i-1)+\rho(i)\cdot\left[\alpha_0^\theta+\sum_{g,s}X_{gs}\pi_{g\ell s}^1(i)\right], &(25)\\ \beta_\ell^\theta(i) &= [1-\rho(i)]\cdot\beta_\ell^\theta(i-1)+\rho(i)\cdot\left[\beta_0^\theta+\sum_{g,s}W_{g\ell}(i)H_{\ell s}(i)\right], &(26)\end{aligned}$$

where $\{\alpha_\ell^\theta(i-1),\beta_\ell^\theta(i-1)\}$ are the values of $\{\alpha_\ell^\theta,\beta_\ell^\theta\}$ from Step 3 of iteration $i-1$. Eq.s (25)-(26) correspond to the third and fourth updates in the third step of each iteration in Algorithm 1 in the main text.

##### Stochastic gradient update of variational parameters for $\mathbf{H}$

We use the same argument as above to verify the conditional conjugacy assumption (4) of SSVI for $\mathbf{H}$:

$$\begin{aligned}\log p(\mathbf{H}) &= \sum_{\ell,s}\left[\left(\alpha_0^H-1\right)\cdot\log H_{\ell s}-\beta_0^H\cdot H_{\ell s}\right]+\text{Const. of }\mathbf{H}, &(27)\\ \log p(\mathbf{H}\,|\,\mathbf{X},\mathbf{Z},\mathbf{W},\boldsymbol{\theta}) &= \log p(\mathbf{X},\mathbf{Z}\,|\,\mathbf{W},\boldsymbol{\theta},\mathbf{H})+\log p(\mathbf{H})+\text{Const. of }\mathbf{H}\\ &= \sum_{\ell,s}\left\{\left[\alpha_0^H+\sum_g\left(Z_{g\ell s}^0+Z_{g\ell s}^1\right)-1\right]\cdot\log H_{\ell s}-\left[\beta_0^H+\sum_g\left(W_{g\ell}\theta_\ell+\lambda\right)\right]\cdot H_{\ell s}\right\}+\begin{matrix}\text{Const.}\\ \text{of }\mathbf{H}.\end{matrix} &(28)\end{aligned}$$

In Eq. (28), we further replace the unobserved variable $Z_{g\ell s}^0$ with $\mathrm{E}_q\left(Z_{g\ell s}^0\,|\,\mathbf{W},\boldsymbol{\theta},\mathbf{H}\right)=X_{gs}\pi_{g\ell s}^0$ and replace $Z_{g\ell s}^1$ with $\mathrm{E}_q\left(Z_{g\ell s}^1\,|\,\mathbf{W},\boldsymbol{\theta},\mathbf{H}\right)=X_{gs}\pi_{g\ell s}^1$ to construct the SSVI stochastic gradient update (8) for variational parameters $\{\alpha_{\ell s}^H,\beta_{\ell s}^H\}$ in Eq. (11).

To implement Step 3 of iteration $i$ in SSVI for $H_{\ell s}$, we update its variational parameters $\{\alpha_{\ell s}^H,\beta_{\ell s}^H\}$ as

$$\begin{aligned}\alpha_{\ell s}^H(i) &= [1-\rho(i)]\cdot\alpha_{\ell s}^H(i-1)+\rho(i)\cdot\left\{\alpha_0^H+\sum_g X_{gs}\left[\pi_{g\ell s}^1(i)+\pi_{g\ell s}^0(i)\right]\right\}, &(29)\\ \beta_{\ell s}^H(i) &= [1-\rho(i)]\cdot\beta_{\ell s}^H(i-1)+\rho(i)\cdot\left\{\beta_0^H+\sum_g\left[W_{g\ell}(i)\theta_\ell(i)+\lambda\right]\right\} &(30)\end{aligned}$$

where $\{\alpha_{\ell s}^H(i-1),\beta_{\ell s}^H(i-1)\}$ are the values of $\{\alpha_{\ell s}^H,\beta_{\ell s}^H\}$ from Step 3 of iteration $i-1$. Eq.s (29)-(30) correspond to the last two updates in the third step of each iteration in Algorithm 1 in the main text.

##### Posterior estimation of global parameters $\{\mathbf{W},\boldsymbol{\theta},\mathbf{H}\}$

We use the following variational distributions obtained from the last iteration ($I$) of SSVI to approximate the posterior distributions of global parameters $\{\mathbf{W},\boldsymbol{\theta},\mathbf{H}\}$:

$$q(W_{g\ell})\approx\mathcal{G}\left(W_{g\ell};\alpha_{g\ell}^W(I),\beta_{g\ell}^W(I)\right),\quad q(\theta_\ell)\approx\mathcal{G}\left(\theta_\ell;\alpha_\ell^\theta(I),\beta_\ell^\theta(I)\right),\quad q(H_{\ell s})\approx\mathcal{G}\left(H_{\ell s};\alpha_{\ell s}^H(I),\beta_{\ell s}^H(I)\right).\qquad(31)$$

We use the means of these variational distributions (31) to estimate the global parameters $\{\mathbf{W}, \boldsymbol{\theta}, \mathbf{H}\}$:

$$\widehat{W}_{g\ell} = \frac{\alpha^{W}_{g\ell}(I)}{\beta^{W}_{g\ell}(I)}, \quad \widehat{\theta}_{\ell} = \frac{\alpha^{\theta}_{\ell}(I)}{\beta^{\theta}_{\ell}(I)}, \quad \widehat{H}_{\ell s} = \frac{\alpha^{H}_{\ell s}(I)}{\beta^{H}_{\ell s}(I)}. \tag{32}$$

Eq. (32) is the output of Algorithm 1 in the main text.

### Supplementary Note 2

We identified the 54 cell-type-specific genes for 10 mouse brain cell types from the companion scRNA-seq data of Slide-seq study[2] as follows. First, we normalized the expression level of gene $g$ in cell type $k$ as

$$W^{0*}_{gk} = \left(\sum_{j=1}^{K} W^{0}_{jk}\right)^{-1} W^{0}_{gk}, \tag{33}$$

where $W^{0}_{gk}$ was the average single-cell expression level of gene $g$ in cell type $k$. On the normalized expression levels of gene $g$ for $K$ cell types $\left\{W^{0*}_{gk}\right\}_{k=1}^{K}$, we further calculated two dispersion measures: entropy

$$E_g = -\sum_{k=1}^{K} W^{0*}_{gk} \log_2\left(W^{0*}_{gk}\right), \tag{34}$$

and bias-corrected Gini coefficient

$$G_g = \left[2(K-1)\sum_{k=1}^{K} W^{0*}_{gk}\right]^{-1} \left(\sum_{i=1}^{K}\sum_{j=1}^{K}\left|W^{0*}_{gi} - W^{0*}_{gj}\right|\right). \tag{35}$$

Next, we selected a set of cell-type-specific genes as

$$\mathscr{P} = \left\{g : \ G_g > 0.8, \ E_g < 1.5, \ \max_k W^{0*}_{gk} > 0.8 \text{ and } \max_k W^{0}_{gk} > 50\right\}. \tag{36}$$

We assigned each gene $g$ in this set to a cell type $k$ such that its normalized expression level was the highest among all cell types, $\arg\max_j W^{0*}_{gj} = k$. We assigned a maximum of 10 genes with the highest normalized expression levels to each cell type. We could not identify any gene $g$ with $\max_k W^{0*}_{gk} > 0.8$ for choroid plexus, granule, astrocyte, and interneuron. Hence, we relaxed the normalized expression level threshold for these cell types as 0.7, 0.6, 0.6, and 0.5 respectively.

### Supplementary Note 3

We devise a metric to evaluate the quality of RETROFIT results obtained from various parameter settings in scenarios where the true composition (**H**) is unknown, e.g., in real data analysis. In these instances, while **H** is unknown, the cell-type-specific gene expression profile, **W**, is often available. Our metric focuses on assessing the correlation between the estimated gene expression profile (**W**) and the corresponding cell-type specific marker genes across all cell types. Specifically, our metric undertakes a two-step approach. It first computes the correlation between the estimated and reference expression profiles for each matched cell type at different parameter choices (e.g., $\lambda$ or mapping approaches). Subsequently, it calculates the mean correlation across various parameter choices and determines the gain in correlation over this mean value for each cell type. The average gain in correlation across all cell types is then utilized as the statistic for assessment. Algorithm 1 below provides detailed steps. The parameter choice that provides the highest average gain in correlation ($\delta$) will be the best choice.

---

**Algorithm 1** Metric for choosing mapping algorithm or $\lambda$

---

Input: $(W^0, W^1, \ldots, W^D)$

$W^0$ is the gene expression profile of cell-type specific marker genes, $W^j$ is the RETROFIT estimate at $\lambda_j$ (or mapping algorithm $j$), $j = 1, \ldots, D$;

**procedure** COMPAREESTIMATE$(W^1, \ldots, W^D, W^0)$
- **for** $i = 1, \ldots, K$ **do** ▷ $K$: number of cell types
  - **for** $j = 1, \ldots, D$ **do**
    - $Corr_{ij} = Correlation(W_i^j, W_i^0)$ ▷ $W_i^j$ and $W_i^0$ are the $i^{th}$ column vector
  - **end for**
  - $\mu_i = \frac{1}{D} \sum_j^D Corr_{ij}$
  - $\delta_{ij} = Corr_{ij} - \mu_i$
- **end for**
- Compute $\delta_j = \frac{\sum_i^K \delta_{ij}}{K}$ for $j = 1, \ldots, D$
- Compute $Corr_j = \frac{\sum_i^K Corr_{ij}}{K}$ for $j = 1, \ldots, D$
- **return** $\delta = (\delta_1, \ldots, \delta_D)$, $Corr = (Corr_1, \ldots, Corr_D)$

**end procedure**

---

#### Supplementary Note 4

To evaluate RETROFIT's capacity to recover anatomical organization and putative cell states directly from spatial transcriptomics (ST) data, without the use of single-cell RNA-seq references or curated marker genes, we analyzed the mouse main olfactory bulb (MOB) dataset originally presented by Ståhl et al.[3] (data source: https://www.spatialresearch.org/resources-published-datasets/doi-10-1126science-aaf2403 ). This dataset was also examined in the STdeconvolve study[4], where it served as an example of fine-scale anatomical features that cannot be captured by clustering alone.

We applied RETROFIT using $L = 12$ latent components with default hyperparameters. One component (denoted X2) showed the highest abundances almost exclusively within the granule cell layer, precisely in the anatomical region corresponding to the rostral migratory stream (RMS), a narrow structure known to contain neuronal precursor cells (Supplementary Fig. 19a). This component also displayed elevated expression of three canonical RMS-associated genes (*Nrep*, *Sox11*, and *Dcx*), which are well-established markers of neuronal differentiation and neurogenesis (Supplementary Fig. 19b). These spatial and transcriptional patterns match the known biology of the RMS and are consistent with prior findings from STdeconvolve.

To independently validate the spatial localization of these RMS-associated genes, we examined publicly available in situ hybridization (ISH) images for Nrep and Sox11 from the Allen Brain Atlas. For both genes, the ISH staining exhibits a sharply localized band of expression traversing the granule cell layer in the region where the RMS is anatomically expected (Supplementary Fig. 19c). This high-resolution staining provides orthogonal support for the RETROFIT-identified component X2, confirming that its spatial footprint and gene-expression signature correspond to the neuronal precursor population of the RMS.

Together, these results demonstrate that RETROFIT can recover subtle, spatially organized biological structures and putative cell populations directly from ST data, without relying on external reference profiles or marker genes. This analysis illustrates RETROFIT's capability to extract fine-grained histological information from spatial transcriptomics measurements.

### Supplementary Note 5

**Highest achievable correlation for STdeconvolve in Visium HD dataset.**

Because the target cell types and their marker genes are known for this benchmark, we performed a diagnostic upper-bound evaluation for STdeconvolve to assess whether its low performance could be attributed to component annotation rather than model limitations. Supplementary Figure 20 shows, for each cell type, the STdeconvolve component whose estimated proportions achieved the highest Pearson correlation with the corresponding marker-gene expression.

This non-standard procedure provides an optimistic upper bound on STdeconvolve's performance rather than a valid annotation, as components may map to multiple cell types. Even under this favorable evaluation, STdeconvolve remained substantially worse than RETROFIT across all four cell types.

### Supplementary Note 6

**Sensitivity analysis using Starfysh on the human intestine dataset**

To examine the influence of auxiliary marker information on spatial deconvolution, we conducted a targeted sensitivity analysis using Starfysh[5] (v1.2.0), a recently proposed method that can optionally incorporate histological images and curated marker genes, on the human intestine 12 PCW dataset. Because RETROFIT and other reference-free methods evaluated in the main text do not use histological data, Starfysh was run without histological features to enable a comparison focused on transcriptomic information alone.

The human intestine 12 PCW dataset includes a curated list of marker genes and was analyzed in the main Results. When Starfysh was run without marker genes, the algorithm was stalled at the archtypical analysis (AA) stage. When run with marker genes (its recommended configuration), Starfysh completely successfully.

Using the same evaluation metric as in Table 1 in the section "RETROFIT extracts relevant cellular compartments from human intestine Visium data" (i.e. Pearson correlation between estimated cell-type proportions and spatial expression of corresponding marker genes), Starfysh achieved the highest correlation for 3 of 8 cell types, while RETROFIT achieved the highest correlation for another 3 (Supplementary Table 1, left panel).

Because Starfysh uses marker genes both as inputs and as evaluation targets, this comparison inherently favors methods that directly incorporate marker information. To assess robustness beyond the specific markers used for inference, we performed a held-out marker evaluation. For each cell type, marker genes were randomly split into two disjoint sets. Starfysh was run using only the first half, and performance was evaluated using the held-out half. RETROFIT, which does not use marker genes during deconvolution, was annotated and evaluated using the same held-out markers for a symmetric comparison.

Under this held-out evaluation (Supplementary Note 6 Table 1, right panel), RETROFIT achieved higher correlations than Starfysh for most cell types, with comparable performance for epithelial cells. We additionally observed that Starfysh's performance varied with the number and composition of marker genes provided (data not shown), whereas RETROFIT's results were stable across marker configurations due to its reference-free design.

| Compartment | 12 PCW (**full marker**) | | | | | 12 PCW (**held-out marker**) | |
|---|---|---|---|---|---|---|---|
| | RETROFIT | STdeconvolve | CARDfree | Cell2location | Starfysh | RETROFIT | Starfysh |
| Endothelial | **0.68** | 0.42 | 0.36 | 0.62 | 0.53 | **0.64** | 0.27 |
| Epithelial | 0.55 | 0.51 | 0.47 | **0.77** | 0.69 | 0.55 | **0.57** |
| Fibroblast | **0.68** | 0.59 | 0.63 | 0.50 | 0.51 | **0.68** | 0.26 |
| Immune | 0.10 | $-8.4 \times 10^{-2}$ | 0.28 | 0.20 | **0.83** | **0.20** | 0.12 |
| Muscle | **0.87** | -0.20 | 0.72 | 0.83 | 0.60 | **0.87** | 0.32 |
| MyoFB/MESO | 0.34 | $-1.2 \times 10^{-2}$ | 0.32 | $8.6 \times 10^{-2}$ | **0.75** | **0.26** | 0.12 |
| Neural | 0.69 | -0.12 | $-9.9 \times 10^{-2}$ | **0.75** | 0.65 | **0.65** | 0.30 |
| Pericyte | 0.11 | $-3.5 \times 10^{-4}$ | $9.8 \times 10^{-2}$ | 0.13 | **0.84** | **0.07** | 0.03 |

**Table 1:** Comparison of RETROFIT and other deconvolution methods on human intestine Visium data (12 PCW). Pearson correlations between ST expression scores of known marker genes and estimated proportions across spots. For each compartment, the highest correlation across methods is shown in bold. **Left panel**: Latent components from the three reference-free methods (RETROFIT, STdeconvolve, CARDfree) were mapped to intestinal cellular compartments using a curated list of 37 compartment marker genes (Algorithm 2). Cell2location was run using the matched scRNA-seq reference. Starfysh was run with the full list of 37 marker genes, and correlations were computed with the same list. **Right panel**: Starfysh was run using only half of the marker genes for each compartment, and correlation were computed using the held-out half. RETROFIT components were annotated using the held-out half and and evaluated using the same subset.

#### Supplementary Figure 1

Comparative analysis using synthetic ST data with different spot size, cell-type complexity and reference quality. The synthetic ST datasets for this study are the same datasets from **Figure 2** with $n = 1000$ simulated spots. Within the violin plots, the white box represents the interquartile range (IQR), the central horizontal line indicates the median, and the whiskers extend to 1.5x the IQR. **a**: small spots ($N = 10$ cells per spot) with low cell-type complexity (up to $M = 3$ cell types per spot from $K = 10$ cell types). **b**: large spots ($N = 20$) with high cell-type complexity ($M = 5$ and $K = 10$). **c, d**: $N = 10, M = 3$ and $K = 5$. Reference-based methods were provided with the following single-cell transcriptomic references. **a, b**: exact reference of all 10 ground-truth cell types. **c**: all 5 ground-truth plus 5 irrelevant cell types. **d**: only 3 out of 5 ground truth plus 5 irrelevant cell types. All figures show the distribution of NRMSE that is not clamped. The one-sided KS $P$-values are shown (black: $P < 0.05$; brown: $P > 0.05$). A small $P$-value indicates that RETROFIT estimates have stochastically lower RMSE compared to another method.

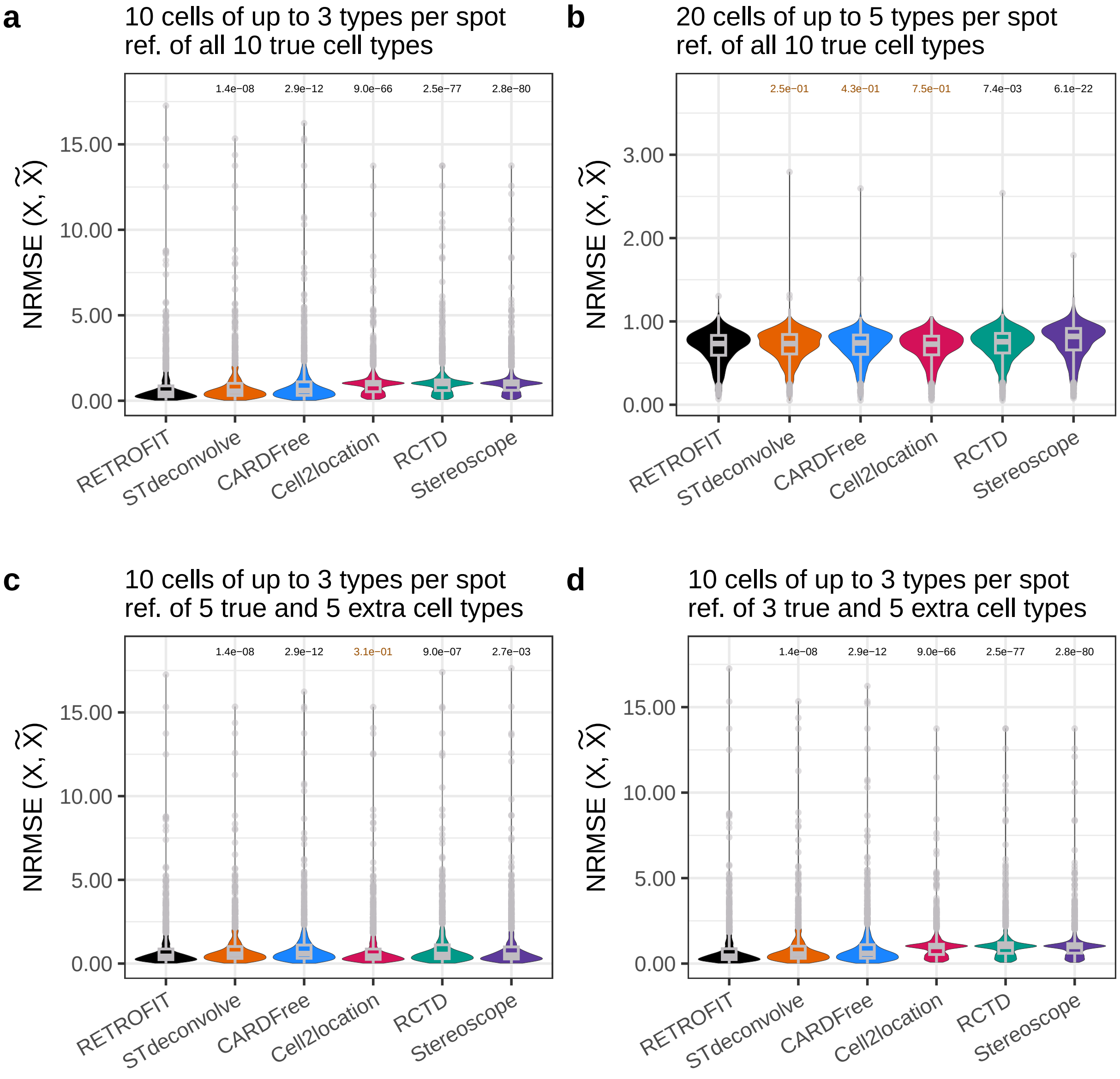
a
10 cells of up to 3 types per spot
ref. of all 10 true cell types
1.4e−08
2.9e−12
9.0e−66
2.5e−77
2.8e−80
NRMSE (X, X̃)
15.00
10.00
5.00
0.00
RETROFIT
STdeconvolve
CARDFree
Cell2location
RCTD
Stereoscope
b
20 cells of up to 5 types per spot
ref. of all 10 true cell types
2.5e−01
4.3e−01
7.5e−01
7.4e−03
6.1e−22
NRMSE (X, X̃)
3.00
2.00
1.00
0.00
RETROFIT
STdeconvolve
CARDFree
Cell2location
RCTD
Stereoscope
c
10 cells of up to 3 types per spot
ref. of 5 true and 5 extra cell types
1.4e−08
2.9e−12
3.1e−01
9.0e−07
2.7e−03
NRMSE (X, X̃)
15.00
10.00
5.00
0.00
RETROFIT
STdeconvolve
CARDFree
Cell2location
RCTD
Stereoscope
d
10 cells of up to 3 types per spot
ref. of 3 true and 5 extra cell types
1.4e−08
2.9e−12
9.0e−66
2.5e−77
2.8e−80
NRMSE (X, X̃)
15.00
10.00
5.00
0.00
RETROFIT
STdeconvolve
CARDFree
Cell2location
RCTD
Stereoscope

### Supplementary Figure 2

Comparative analysis using synthetic data with small spots ($N = 10$ cells per spot) with low cell-type complexity (up to $M = 3$ cell types per spot from $K = 5$ cell types). The synthetic ST data for this study is the same dataset as used for **Figure 2 column 4** with $n = 1000$ simulated spots. Reference-based methods are provided with single-cell transcriptomic reference consisting of only 3 out of 5 true cell types, plus 5 irrelevant cell types.

The left column shows the assessment based on the 5 true cell types used to generate the ST data, while the right column displays the assessment based only on the 3 true cell types present in the imperfect reference. **a** Distribution of RMSE and **b** ranked correlation between true ($\mathbf{H}$) and estimated cell-type proportions ($\widetilde{\mathbf{H}}$) across all cell types at each spot. Within the violin plots, the white box represents the interquartile range (IQR), the central horizontal line indicates the median, and the whiskers extend to 1.5x the IQR. The one-sided KS $P$-values are shown in **a** (black: $P < 0.05$; brown: $P > 0.05$). A small $P$-value indicates that RETROFIT estimates have stochastically lower RMSEs compared to another method. The Area Under the Curve (AUC) of ranked correlations is shown for each method with a matching color in **b**.

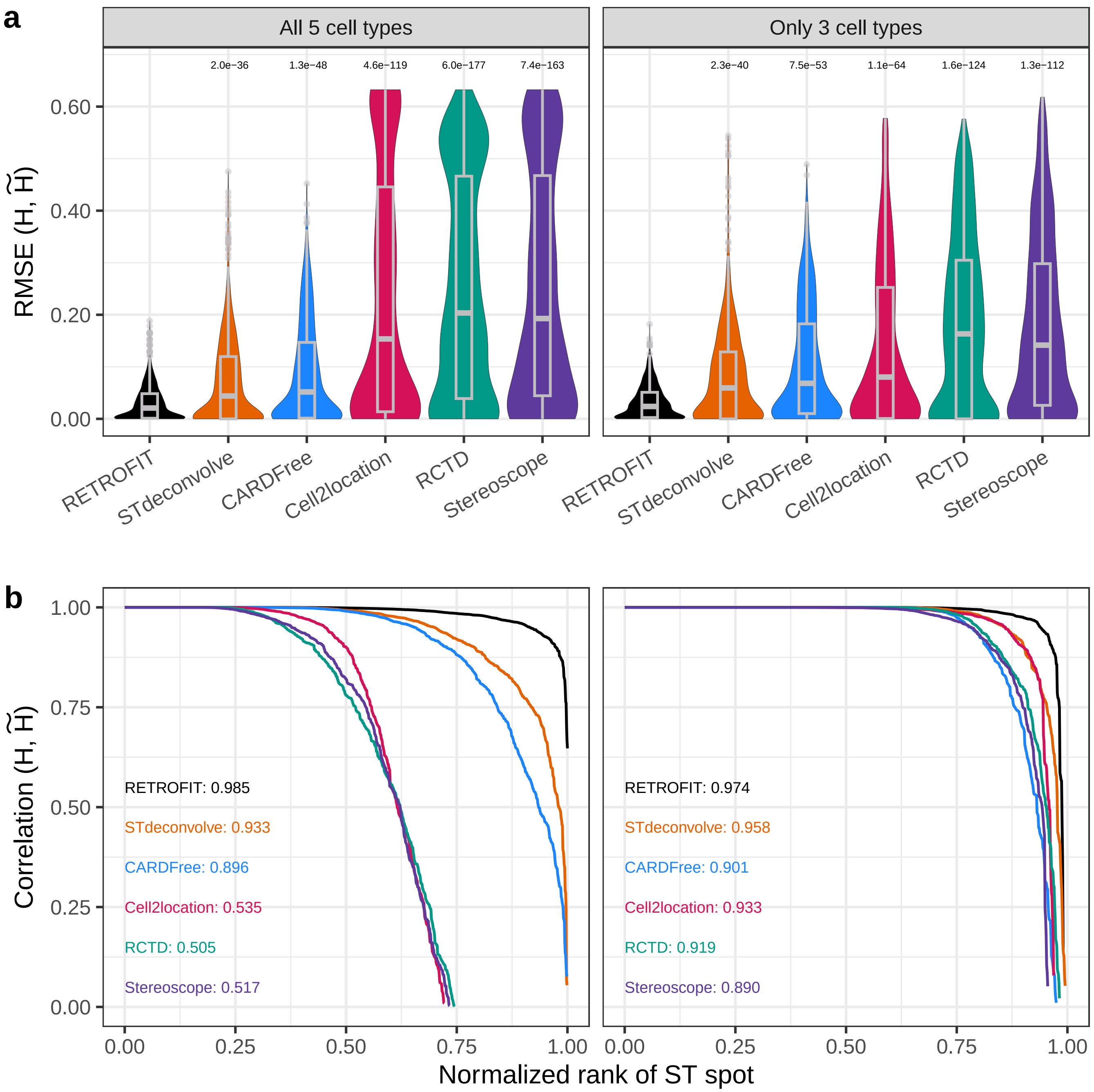
a
All 5 cell types
Only 3 cell types
RMSE (H, H̃)
2.0e−36
1.3e−48
4.6e−119
6.0e−177
7.4e−163
2.3e−40
7.5e−53
1.1e−64
1.6e−124
1.3e−112
RETROFIT
STdeconvolve
CARDFree
Cell2location
RCTD
Stereoscope
b
Correlation (H, H̃)
RETROFIT: 0.985
STdeconvolve: 0.933
CARDFree: 0.896
Cell2location: 0.535
RCTD: 0.505
Stereoscope: 0.517
RETROFIT: 0.974
STdeconvolve: 0.958
CARDFree: 0.901
Cell2location: 0.933
RCTD: 0.919
Stereoscope: 0.890
Normalized rank of ST spot

#### Supplementary Figure 3

Comparative analysis using synthetic ST data with high resolution. The synthetic ST data for this study is generated by applying Algorithm 4 to the mouse cerebellum scRNA-seq dataset[2] as described in methods section **Simulation Studies** with small spots ($N = 3$ cells per spot) with low cell-type complexity (up to $M = 2$ cell types per spot from $K = 10$ cell types). Reference-based methods were provided with the following single-cell transcriptomic references. Columns 1: exact reference of all 10 ground-truth cell types. Column 2: only 8 out of 10 ground truth cell types.

**a** Distribution of RMSE and **b** ranked correlation between true ($\mathbf{H}$) and estimated cell-type proportions ($\widetilde{\mathbf{H}}$) across all cell types at each spot. **c** Distribution of NRMSE and **d** ranked correlation between observed ($\mathbf{X}$) and reconstructed expression ($\widetilde{\mathbf{X}}$) across all genes at each spot. All panels represent $n = 1000$ total spots. In the violin plots, the center line represents the median, the box limits represent the interquartile range (IQR), and the whiskers extend to the maximum and minimum values within 1.5x the IQR. The one-sided KS $P$-values are shown in **a** and **c** (black: $P < 0.05$; brown: $P > 0.05$). A small $P$-value indicates that RETROFIT estimates have stochastically lower RMSEs compared to another method. The AUC of ranked correlations is shown for each method with a matching color in **b** and **d**.

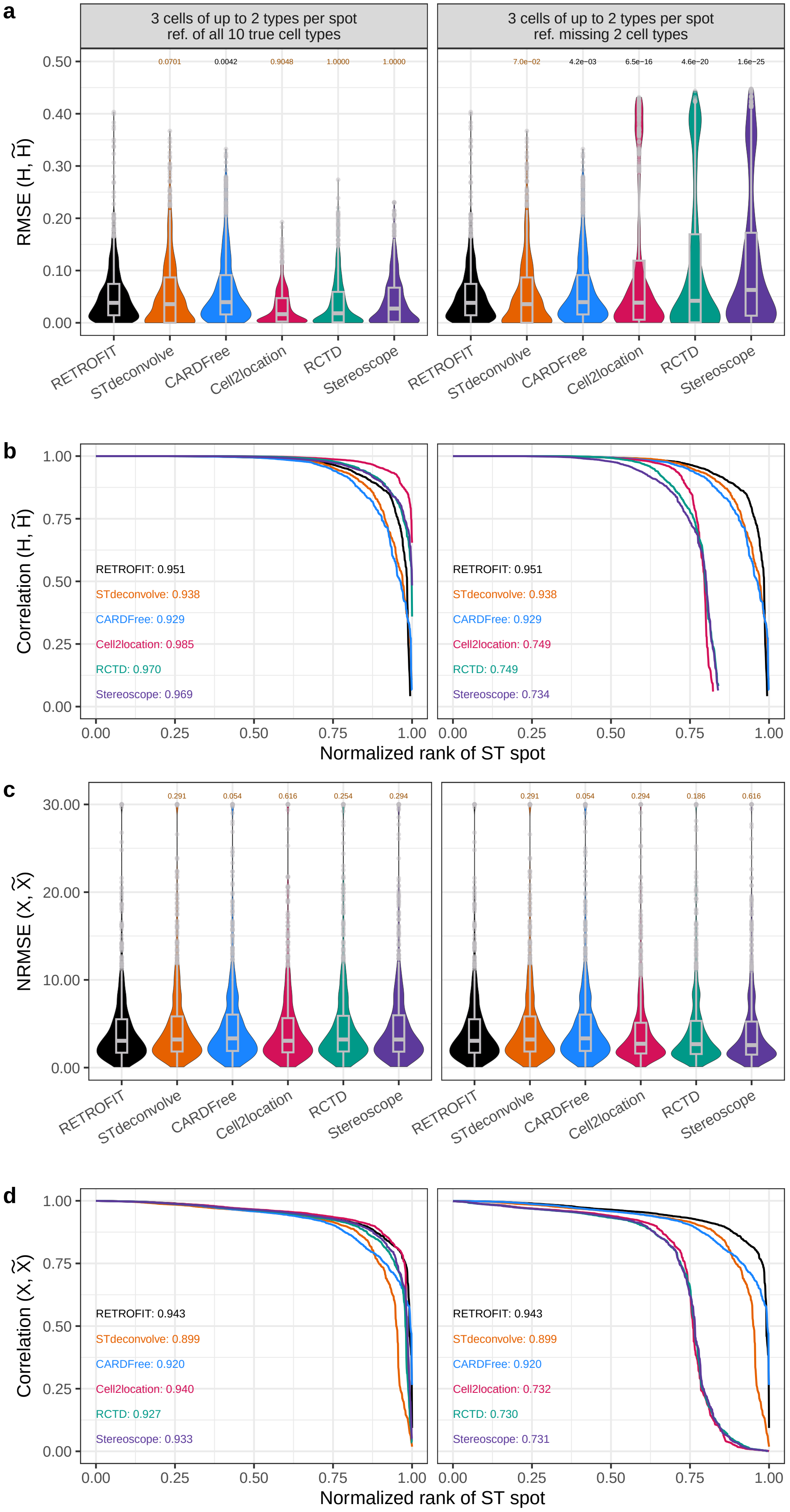
a
3 cells of up to 2 types per spot
ref. of all 10 true cell types
3 cells of up to 2 types per spot
ref. missing 2 cell types
RMSE (H, H̃)
0.50
0.40
0.30
0.20
0.10
0.00
0.0701
0.0042
0.9048
1.0000
1.0000
7.0e−02
4.2e−03
6.5e−16
4.6e−20
1.6e−25
RETROFIT
STdeconvolve
CARDFree
Cell2location
RCTD
Stereoscope
b
Correlation (H, H̃)
1.00
0.75
0.50
0.25
0.00
RETROFIT: 0.951
STdeconvolve: 0.938
CARDFree: 0.929
Cell2location: 0.985
RCTD: 0.970
Stereoscope: 0.969
RETROFIT: 0.951
STdeconvolve: 0.938
CARDFree: 0.929
Cell2location: 0.749
RCTD: 0.749
Stereoscope: 0.734
Normalized rank of ST spot
c
NRMSE (X, X̃)
30.00
20.00
10.00
0.00
0.291
0.054
0.616
0.254
0.294
0.291
0.054
0.294
0.186
0.616
d
Correlation (X, X̃)
RETROFIT: 0.943
STdeconvolve: 0.899
CARDFree: 0.920
Cell2location: 0.940
RCTD: 0.927
Stereoscope: 0.933
RETROFIT: 0.943
STdeconvolve: 0.899
CARDFree: 0.920
Cell2location: 0.732
RCTD: 0.730
Stereoscope: 0.731
Normalized rank of ST spot

#### Supplementary Figure 4

Comparative analysis using synthetic ST data when two major or two minor cell types are missing in the reference. The synthetic ST data for this study is generated by applying Algorithm 4 to the mouse cerebellum scRNA-seq dataset[2] as described in methods section **Simulation Studies** with $N = 10, M = 3, K = 5$. The dataset consists of 30% Purkinje, 30% endothelial, 20% Bergmann glia, 10% choroid plexus, and 10% oligodendrocyte. The reference-based methods were provided with following imperfect references. Left column: reference lacked the two major cell types, Purkinje and endothelial. Right column: reference lacked the minor cell types, choroid plexus and oligodendrocyte.

**a** Distribution of RMSE and **b** ranked correlation between true ($\mathbf{H}$) and estimated cell-type proportions ($\widetilde{\mathbf{H}}$) across all cell types at each spot. **c** Distribution of NRMSE and **d** ranked correlation between observed ($\mathbf{X}$) and reconstructed expression ($\widetilde{\mathbf{X}}$) across all genes at each spot. All panels represent $n = 1000$ total spots. In the violin plots, the center line represents the median, the box limits represent the interquartile range (IQR), and the whiskers extend to the maximum and minimum values within 1.5x the IQR. The one-sided KS $P$-values are shown in **a** and **c** (black: $P < 0.05$; brown: $P > 0.05$). A small $P$-value indicates that RETROFIT estimates have stochastically lower RMSEs compared to another method. The AUC of ranked correlations is shown for each method with a matching color in **b** and **d**.

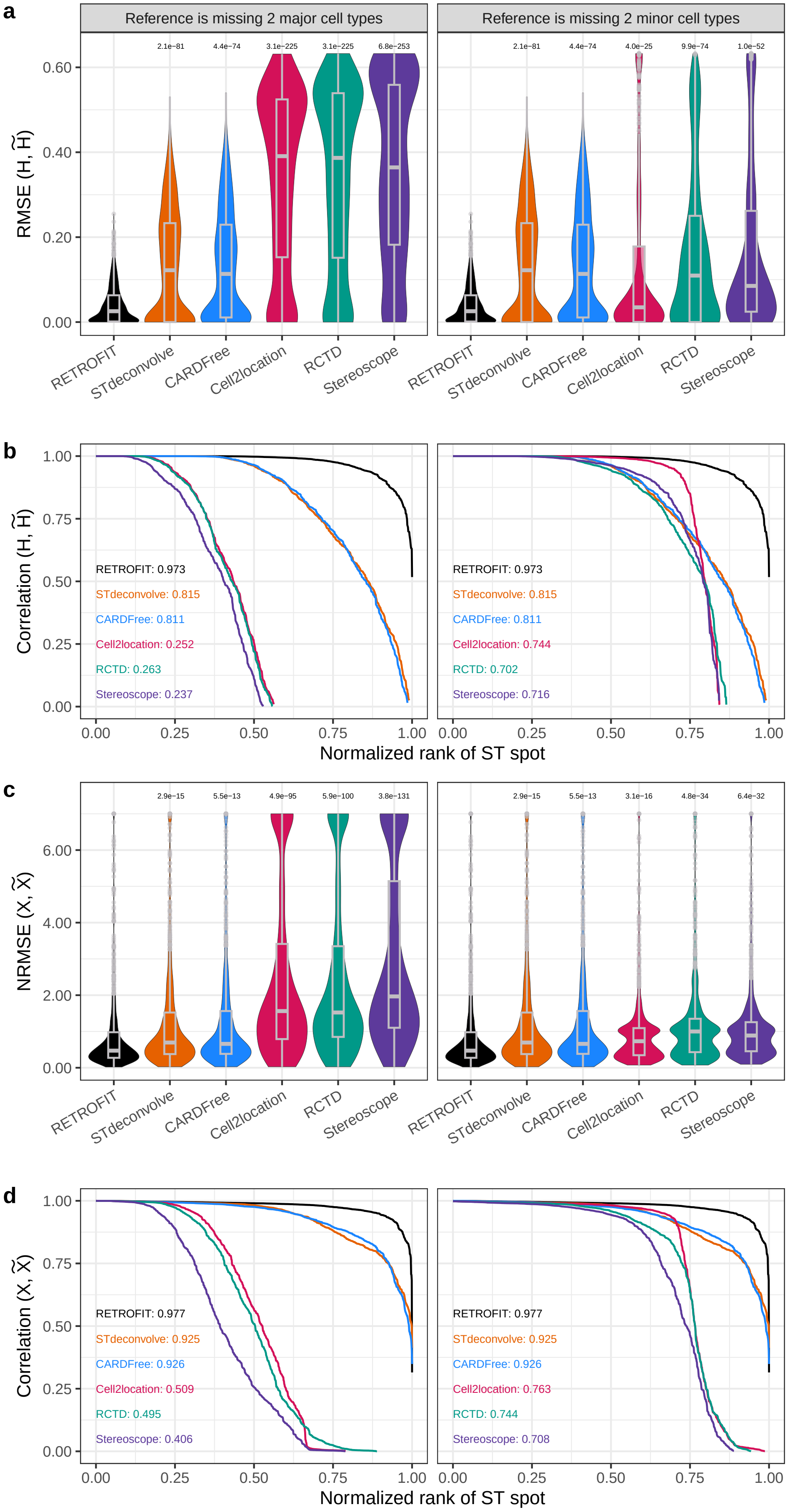
a
Reference is missing 2 major cell types
Reference is missing 2 minor cell types
RMSE (H, H̃)
2.1e−81
4.4e−74
3.1e−225
3.1e−225
6.8e−253
2.1e−81
4.4e−74
4.0e−25
9.9e−74
1.0e−52
RETROFIT
STdeconvolve
CARDFree
Cell2location
RCTD
Stereoscope
b
Correlation (H, H̃)
RETROFIT: 0.973
STdeconvolve: 0.815
CARDFree: 0.811
Cell2location: 0.252
RCTD: 0.263
Stereoscope: 0.237
RETROFIT: 0.973
STdeconvolve: 0.815
CARDFree: 0.811
Cell2location: 0.744
RCTD: 0.702
Stereoscope: 0.716
Normalized rank of ST spot
c
NRMSE (X, X̃)
2.9e−15
5.5e−13
4.9e−95
5.9e−100
3.8e−131
2.9e−15
5.5e−13
3.1e−16
4.8e−34
6.4e−32
d
Correlation (X, X̃)
RETROFIT: 0.977
STdeconvolve: 0.925
CARDFree: 0.926
Cell2location: 0.509
RCTD: 0.495
Stereoscope: 0.406
RETROFIT: 0.977
STdeconvolve: 0.925
CARDFree: 0.926
Cell2location: 0.763
RCTD: 0.744
Stereoscope: 0.708
Normalized rank of ST spot

#### Supplementary Figure 5

Comparative analysis using simulated data from STdeconvolve[4]. This dataset is created by overlaying an artificial grid onto MERFISH data, which offers single-cell resolution, and assigning cells to ”spots” based on their overlapping regions. This simulated data utilizes gene counts and metadata extracted from cells profiled via MERFISH[6]. It encompasses 12 bregma, 9 distinct cell types, 135 genes, and a total of 59,651 cells. The grid dimension of 100 $\mu m$ x 100 $\mu m$ was generated to mimic spots of varying sizes. For each grid size, the composition of cell types within each spot is determined by computing the proportion of cells for each type. The gene expression within a spot is obtained by aggregating the expression levels of individual genes across all cells within that spot. All panels represent $n = 3072$ total spots.

For reference-free methods (RETROFIT and STdeconvolve), the cell types are annotated using a single-cell reference obtained by computing the average expression levels of each cell type on the MERFISH data. This reference is also used as the single-cell reference for the reference-based method, Stereoscope. The single cell reference provided in the RCTD software[7], which analyzed the same MERFISH dataset, is used as the reference for the other two reference-based methods, RCTD and Cell2location. Lastly, CARDfree is inapplicable to this simulated dataset due to the insufficient number of marker genes for CARDfree to generate results.

The performance of methods is evaluated when the reference is imperfect. To create an imperfect reference, the two most prevalent cell types in this sample, “Inhibitory” and “Excitatory”, are removed from the single-cell reference for reference-based methods. The same parameter settings as in Main Text Methods Sections **Hyperparameter specification** and **Existing methods for comparison** are used to run each method and the same evaluation methods as in Main Text **Figure 2** are used to compare the performance. The one-sided KS $P$-values are shown in a and b.

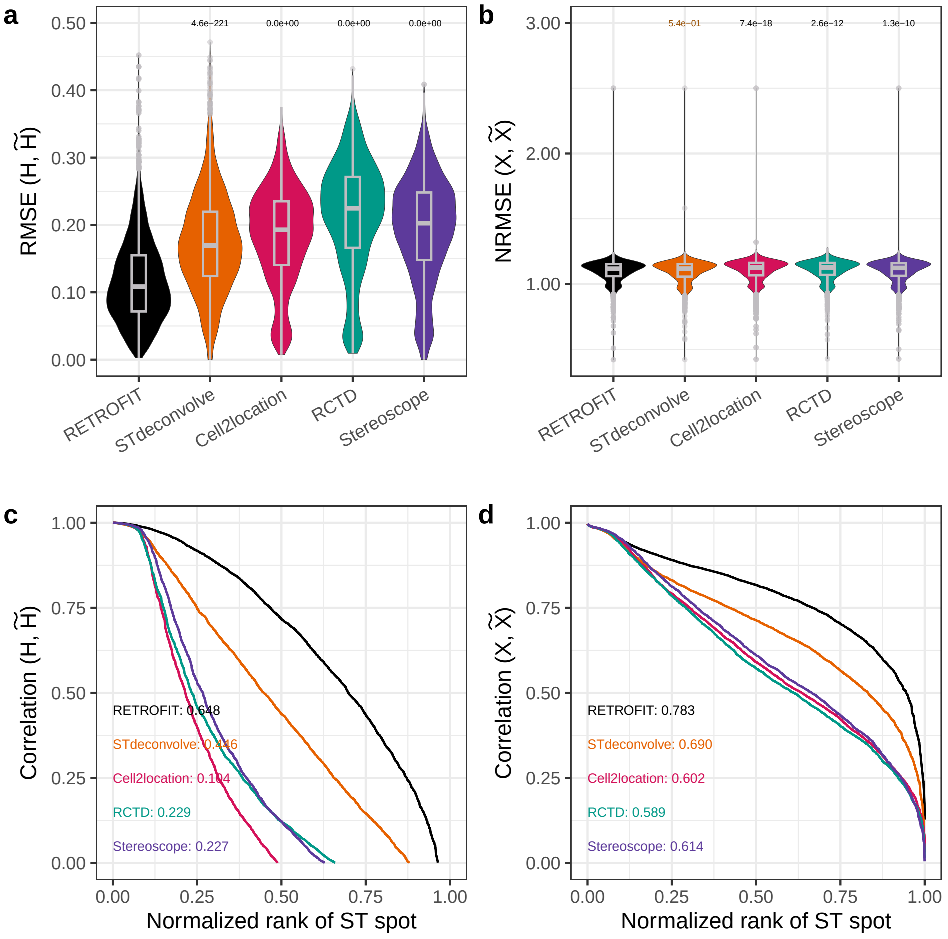

#### Supplementary Figure 6

Comparative analysis using simulated data with similar cell types. The dataset is generated from MERFISH data[6], which comprised 9 cell types totaling 59,651 cells. From the 37 subtypes of the "Inhibitory" cell type category, 5 subtypes are selected (I-4, I-7, I-8, I-13, and I-15), resulting in a total of 5711 cells. The data is processed as detailed in **Supplementary Figure 5**, resulting in $n = 1803$ spots with expression levels for 135 marker genes. Proportions for the selected subtypes are: I-4 (0.230), I-7 (0.258), I-8 (0.168), I-13 (0.154), and I-15 (0.191).

RETROFIT's performance is compared to STdeconvolve, Cell2location, RCTD, and Stereoscope, with I-4 and I-7 missing from the references. CARDfree could not be evaluated due to additional marker gene requirements. RETROFIT is configured with $L = 10$, $\lambda = 0.01$ , 4000 iterations, and uses Algorithm 2 for annotation. STdeconvolve uses $K = 10$, and other methods use their default parameters. The one-sided KS $P$-values are shown in a and b.

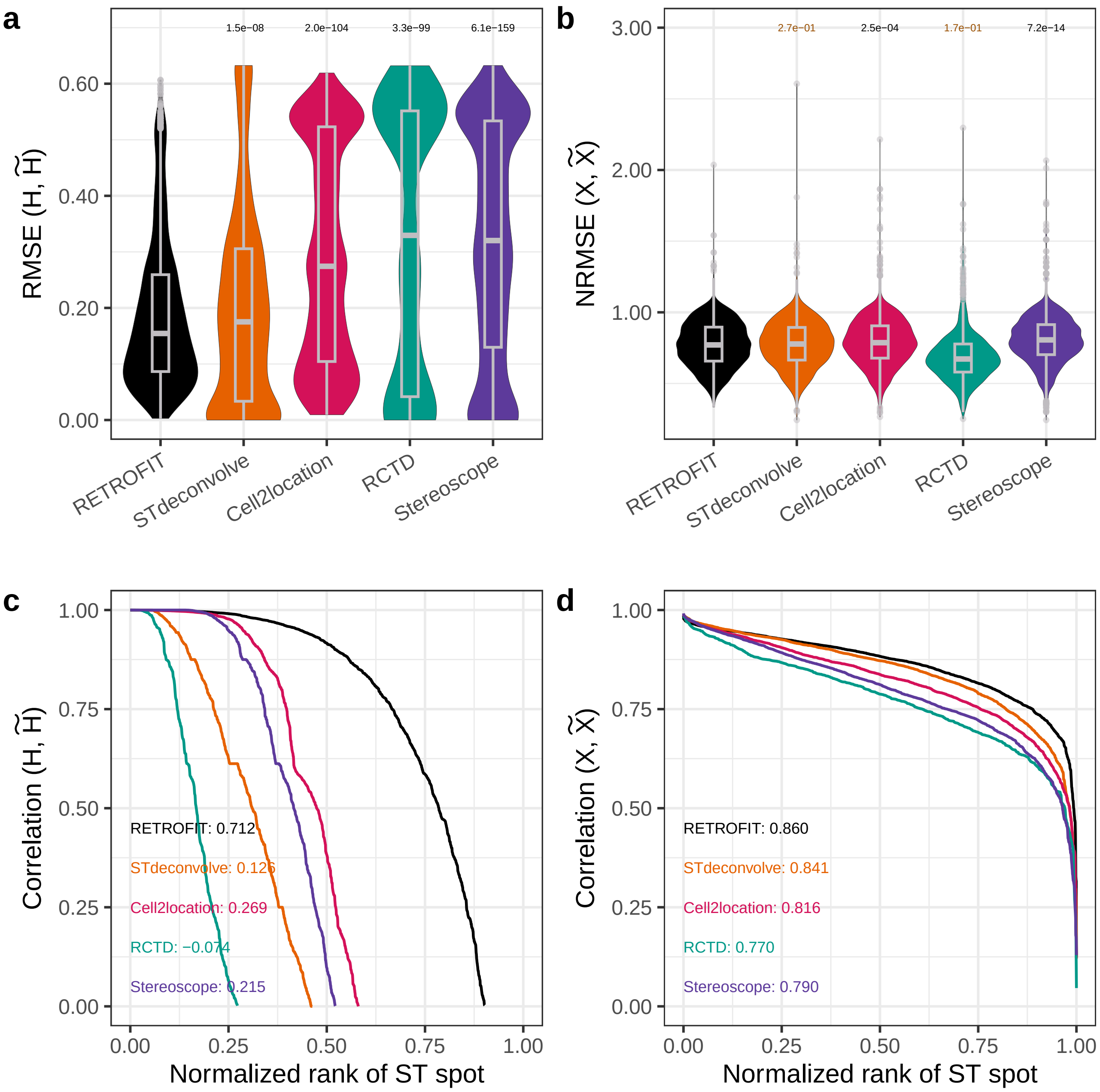

#### Supplementary Figure 7

Endothelial compartments identified by RETROFIT in human fetal and adult intestine Visium data (related to Fig. 5c-d). Shown are ST expression scores of compartment marker genes (row 1) and RETROFIT estimates of compartment proportions (row 2) across spots in the fetal (12 PCW, $n = 1080$ spots; 19 PCW, $n = 1242$ spots) and adult ($n = 2649$ spots) ST slides. Pearson correlation ($R$) between compartment marker ST expression scores and compartment proportion estimates across all spots is also given for each ST slide.

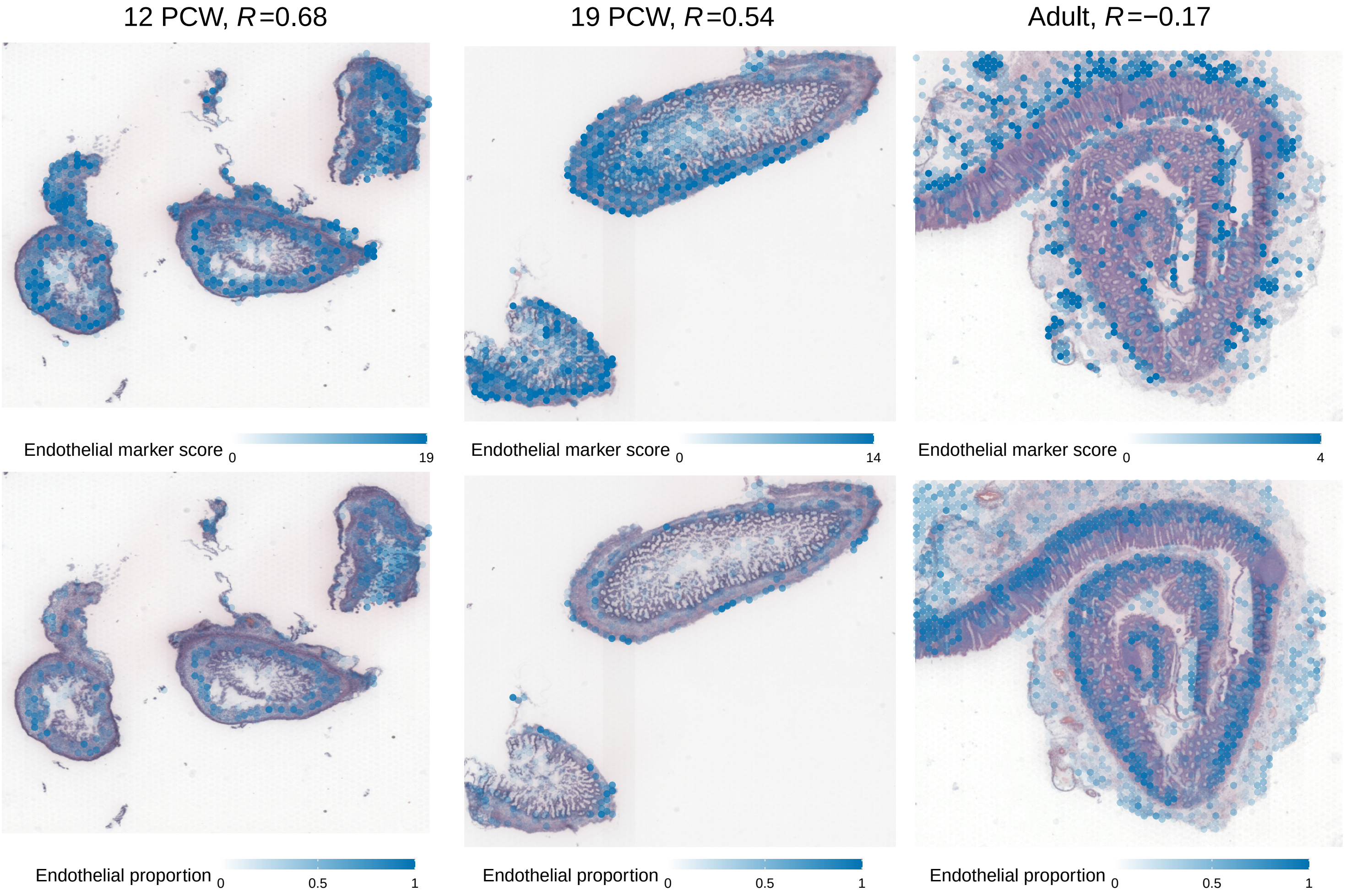

#### Supplementary Figure 8

Fibroblast compartments identified by RETROFIT in human fetal and adult intestine Visium data (related to Fig. 5c-d). Shown are ST expression scores of compartment marker genes (row 1) and RETROFIT estimates of compartment proportions (row 2) across spots in the fetal (12 PCW, $n$ = 1080 spots; 19 PCW, $n$ = 1242 spots) and adult ($n$ = 2649 spots) ST slides. Pearson correlation ($R$) between compartment marker ST expression scores and compartment proportion estimates across all spots is also given for each ST slide.

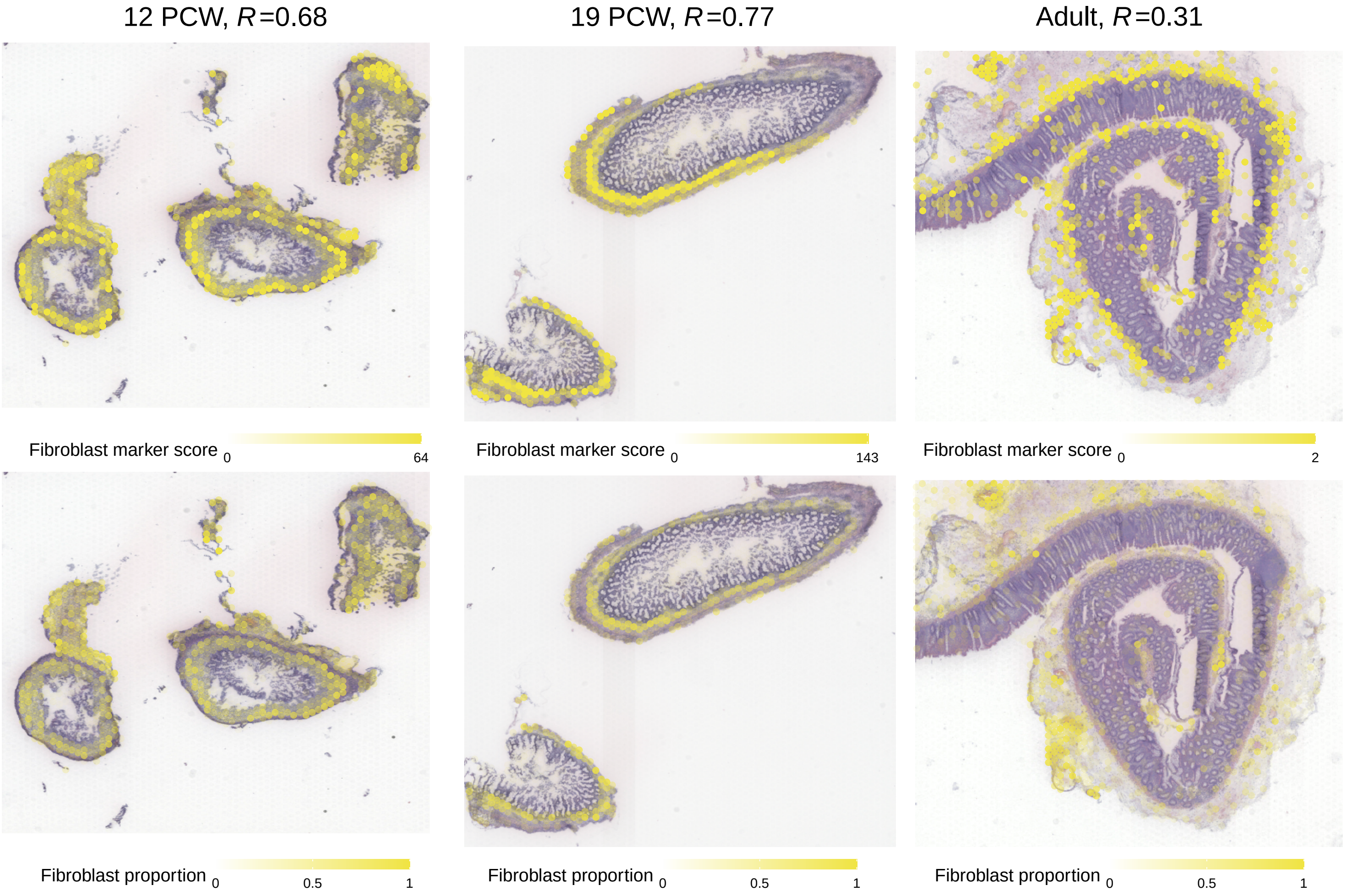

#### Supplementary Figure 9

Immune compartments identified by RETROFIT in human fetal and adult intestine Visium data (related to Fig. 5c-d). Shown are ST expression scores of compartment marker genes (row 1) and RETROFIT estimates of compartment proportions (row 2) across spots in the fetal (12 PCW, $n$ = 1080 spots; 19 PCW, $n$ = 1242 spots) and adult ($n$ = 2649 spots) ST slides. Pearson correlation ($R$) between compartment marker ST expression scores and compartment proportion estimates across all spots is also given for each ST slide.

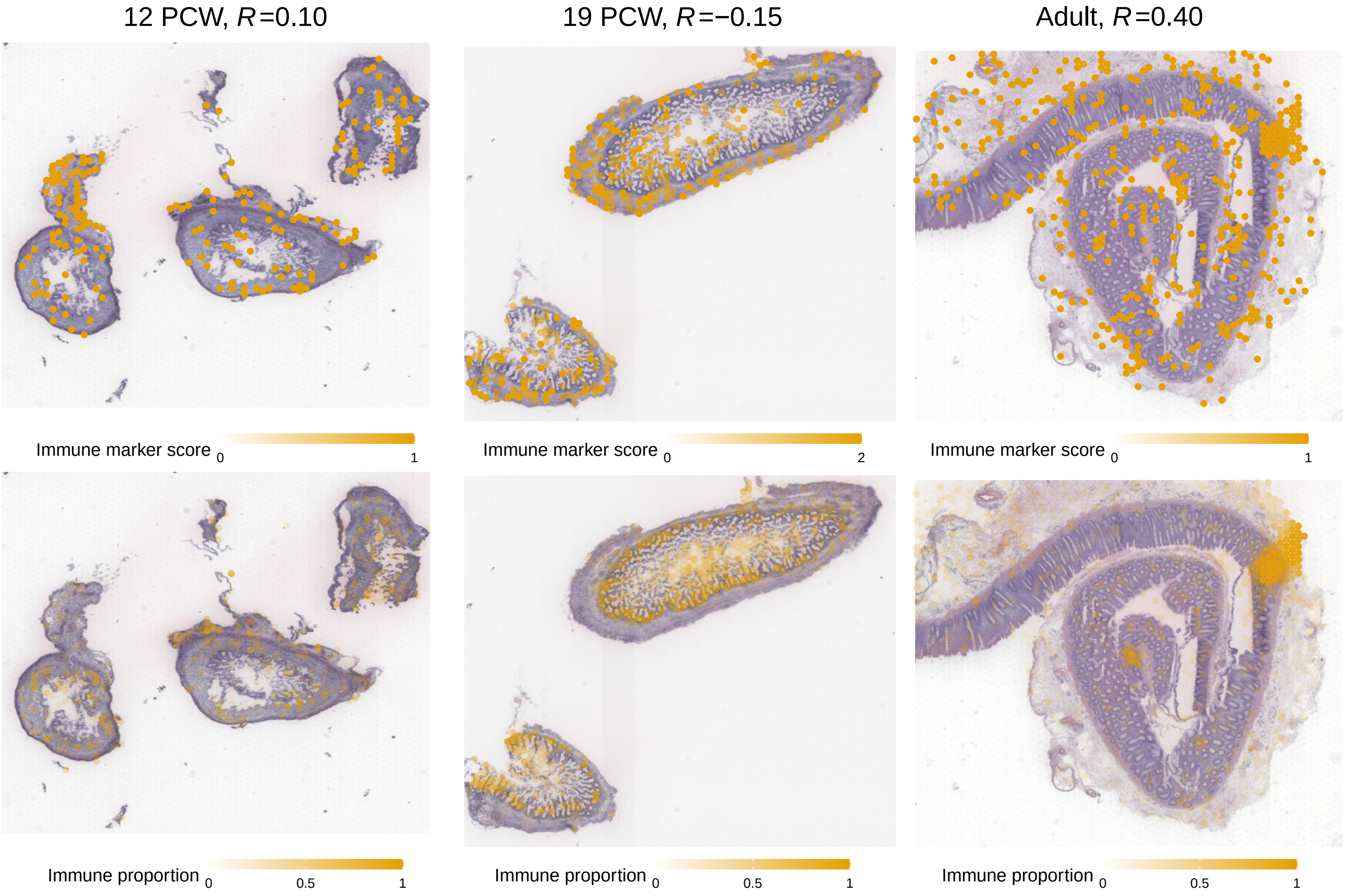

#### Supplementary Figure 10

MyoFB/MESO compartments identified by RETROFIT in human fetal and adult intestine Visium data (related to Fig. 5c-d). Shown are ST expression scores of compartment marker genes (row 1) and RETROFIT estimates of compartment proportions (row 2) across spots in the fetal (12 PCW, $n$ = 1080 spots; 19 PCW, $n$ = 1242 spots) and adult ($n$ = 2649 spots) ST slides. Pearson correlation ($R$) between compartment marker ST expression scores and compartment proportion estimates across all spots is also given for each ST slide.

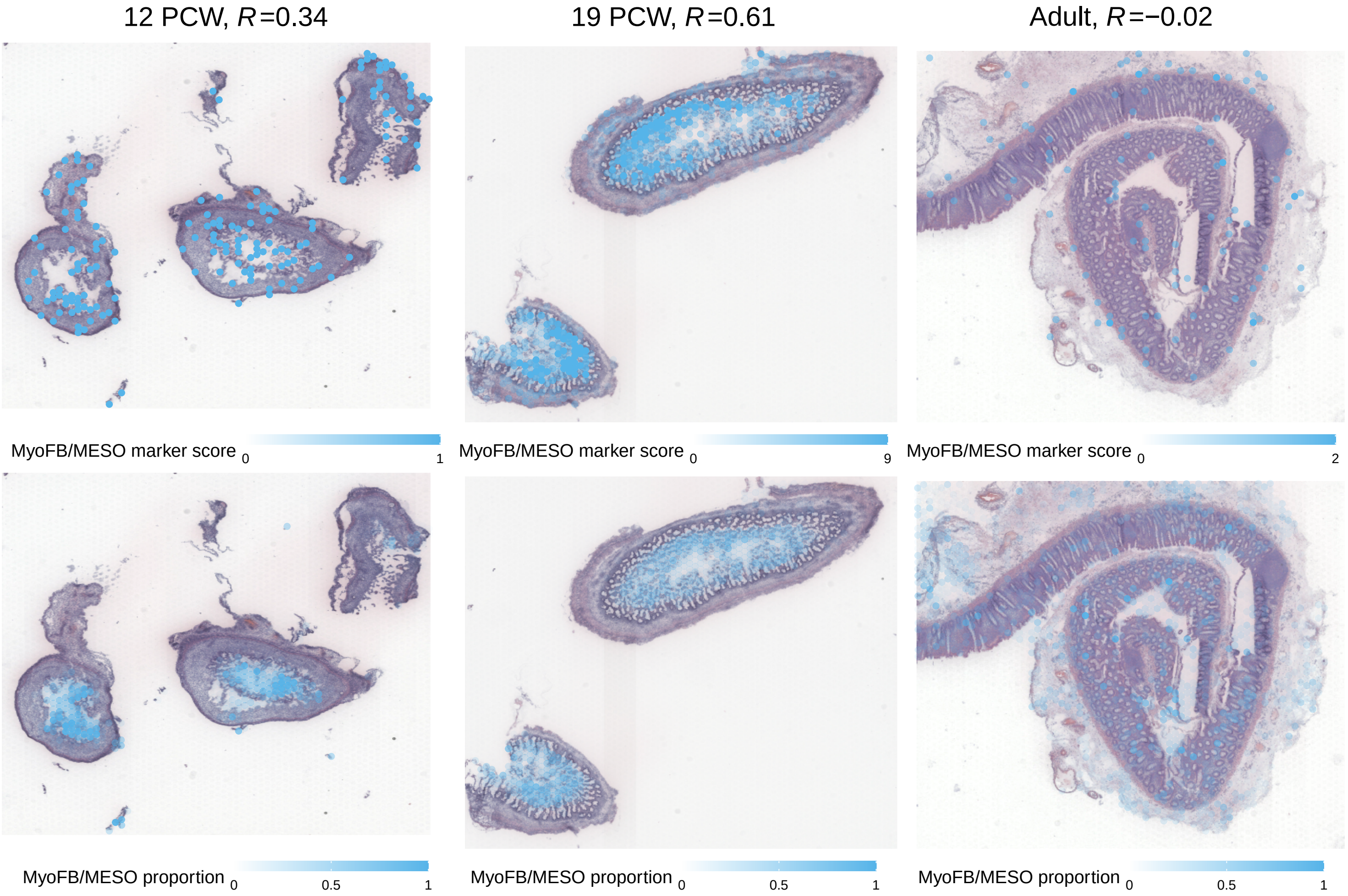

#### Supplementary Figure 11

Neural compartments identified by RETROFIT in human fetal and adult intestine Visium data (related to Fig. 5c-d). Shown are ST expression scores of compartment marker genes (row 1) and RETROFIT estimates of compartment proportions (row 2) across spots in the fetal (12 PCW, $n$ = 1080 spots; 19 PCW, $n$ = 1242 spots) and adult ($n$ = 2649 spots) ST slides. Pearson correlation ($R$) between compartment marker ST expression scores and compartment proportion estimates across all spots is also given for each ST slide.

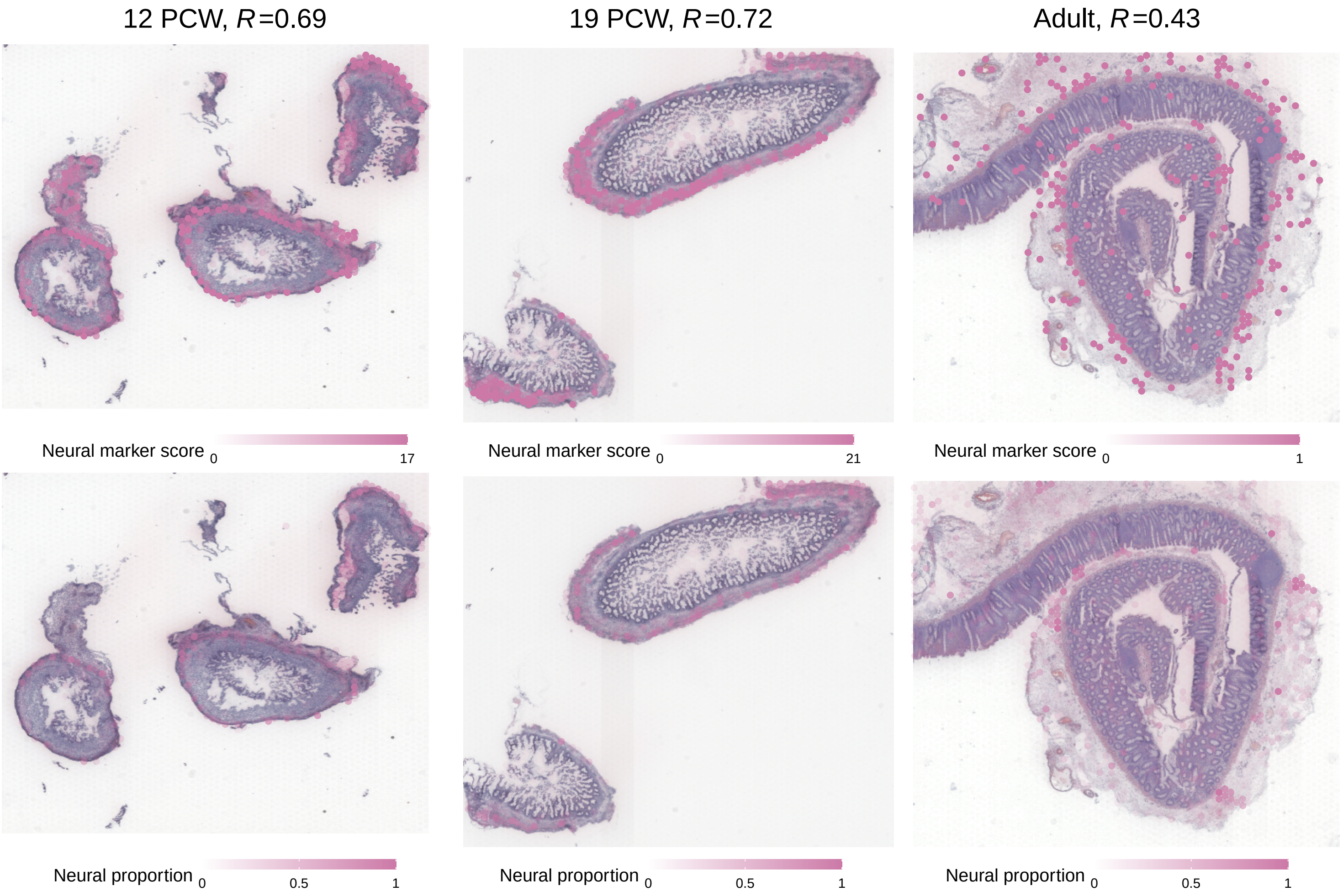

### Supplementary Figure 12

Pericyte compartments identified by RETROFIT in human fetal and adult intestine Visium data (related to Fig. 5c-d). Shown are ST expression scores of compartment marker genes (row 1) and RETROFIT estimates of compartment proportions (row 2) across spots in the fetal (12 PCW, $n = 1080$ spots; 19 PCW, $n = 1242$ spots) and adult ($n = 2649$ spots) ST slides. Pearson correlation ($R$) between compartment marker ST expression scores and compartment proportion estimates across all spots is also given for each ST slide.

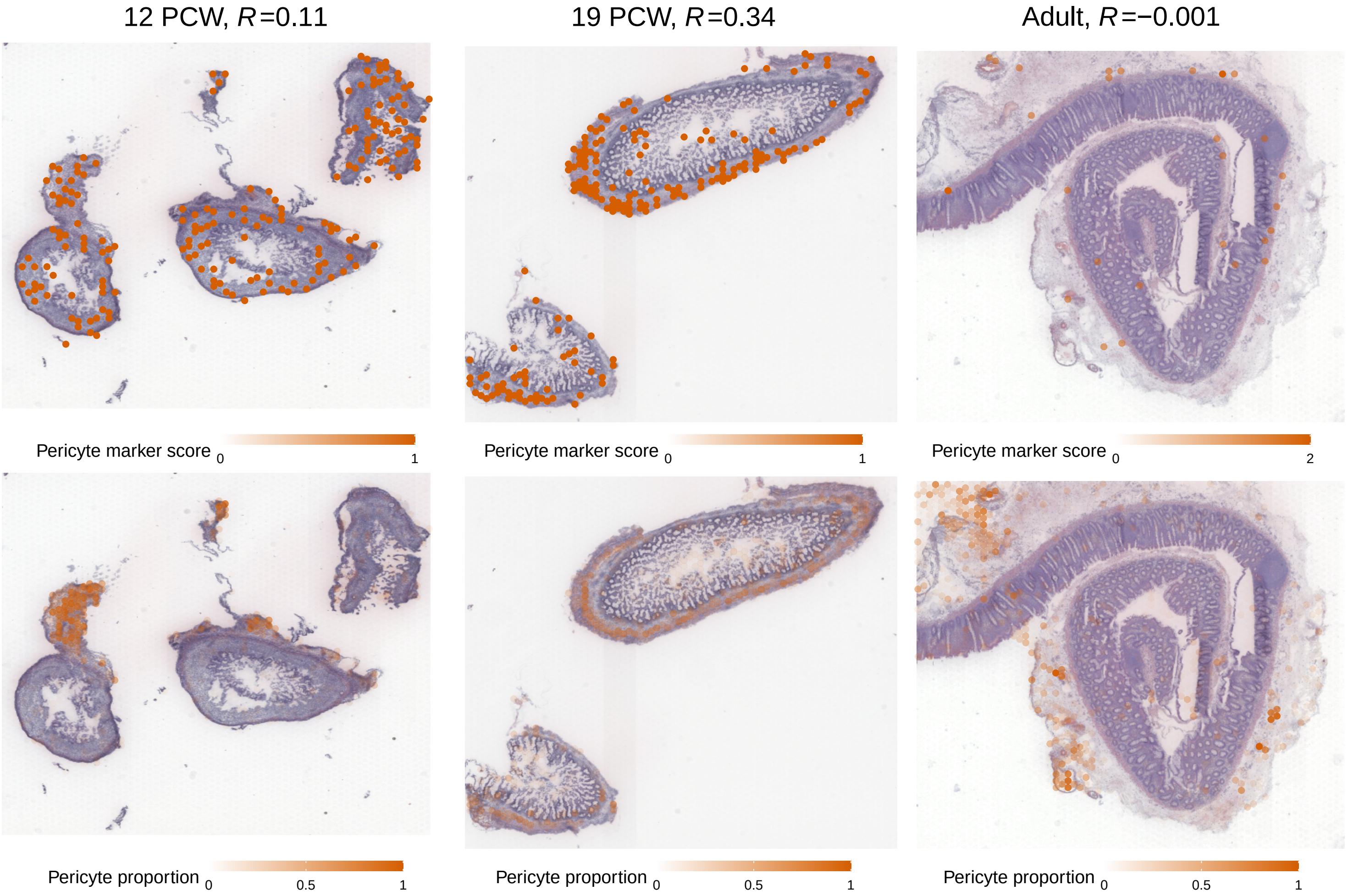

### Supplementary Figure 13

Additional results of compartment co-localization identified by RETROFIT in human intestinal development (related to Fig. 6f-h). Shown are spots with two moderately representative compartments (Group 2) in each of the 3 ST slides (12 PCW, $n$ = 1080 spots; 19 PCW, $n$ = 1242 spots; adult, $n$ = 2649 spots), with the anchor compartments as neural (left), immune (middle) or fibroblast (right) compartment. The color of each spot represents the other cellular compartment that co-localizes with the anchor compartment.

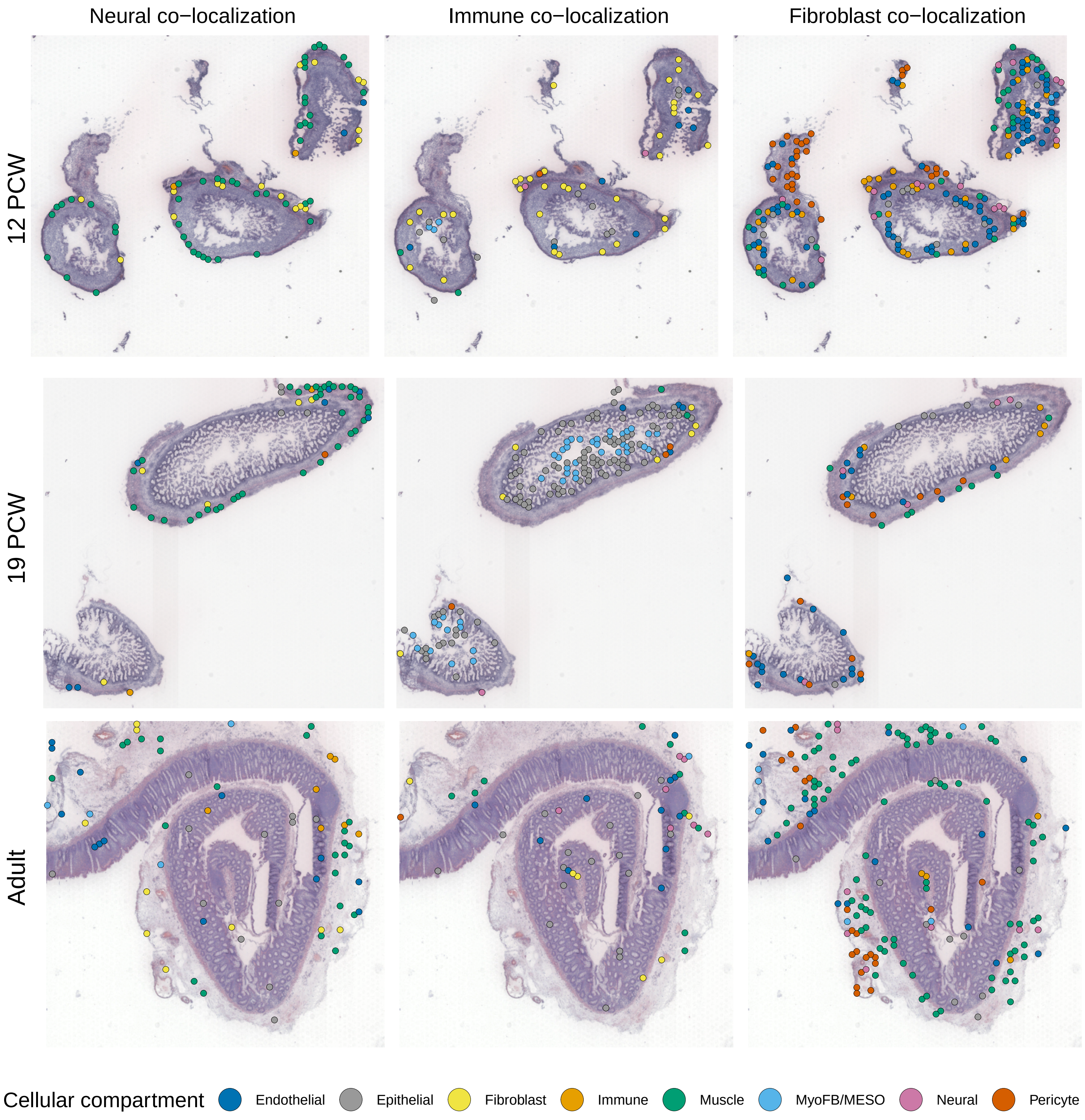

#### Supplementary Figure 14

Comparison between RETROFIT and nonnegative matrix factorization (NMF), a commonly used method for deconvolving matrices, for two simulated datasets used in **Figure 2 Columns 1 and 2**. Column 1 corresponds to small spots with low cell-type complexity (N = 10, M = 3, K = 10), and column 2 corresponds to large spots with higher complexity (N = 20, M = 5, K = 10). In this analysis, the method NMF is referred to as a pipeline where Step 1 in RETROFIT is replaced by NMF and Step 2 is performed as usual. We use the R package *NMF* with Euclidean (Frobenius norm) loss. Performance is evaluated using the area under the curve (AUC) derived from correlations between estimated and true cell-type proportions across spots, shown as a function of the number of components (L).

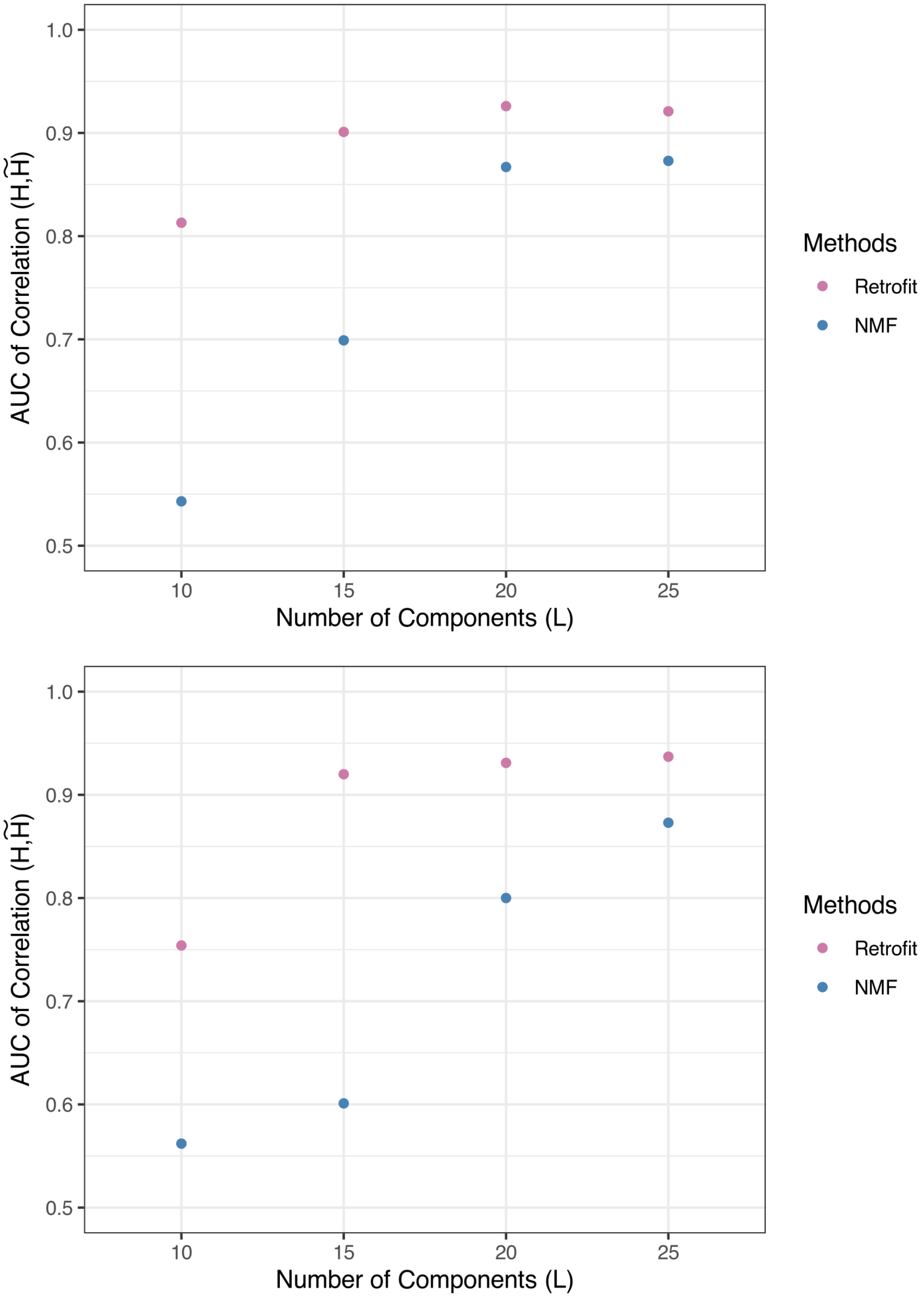

### Supplementary Figure 15

RETROFIT performance with different $L$ values on synthetic ST datasets with different spot size, cell-type complexity and reference quality. The datasets are the same simulated datasets as in Figure 2. Each dataset contains $n = 1000$ spots per simulation. Performance is evaluated using the area under the curve (AUC) derived from correlations between estimated and true cell-type proportions across spots. **a**: small spots ($N = 10$ cells per spot) with low cell-type complexity (up to $M = 3$ cell types per spot from $K = 10$ cell types). **b**: large spots ($N = 20$) with high cell-type complexity ($M = 5$ and $K = 10$). **c**: $N = 10, M = 3$ and $K = 5$.

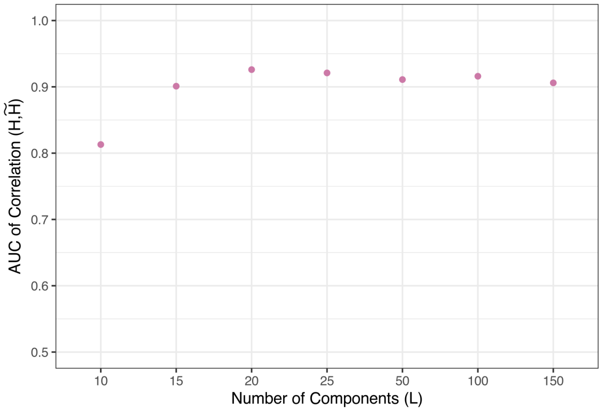


a. $N = 10, M = 3, K = 10$

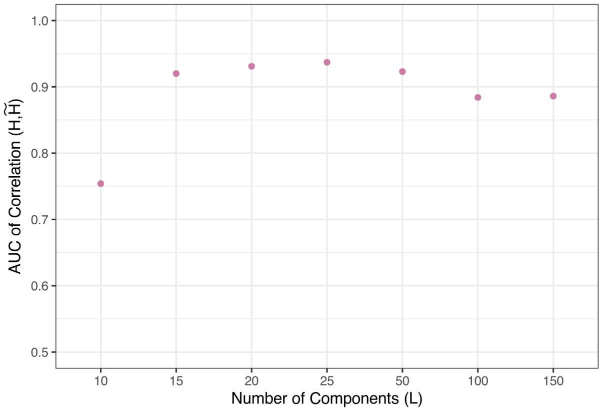


b. $N = 20, M = 5, K = 10$

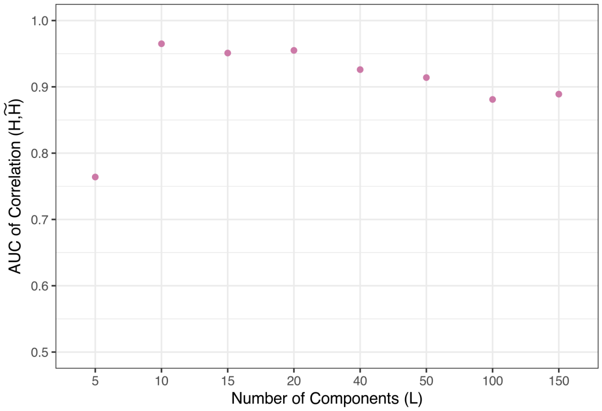


c. $N = 10, M = 3, K = 5$

### Supplementary Figure 16

RETROFIT performance with different $\lambda$ values on synthetic ST datasets with different spot size, cell-type complexity and reference quality. The datasets are the same simulated datasets as in Figure 2. Each dataset contains $n = 1000$ spots per simulation. Performance is evaluated using the area under the curve (AUC) derived from correlations between estimated and true cell-type proportions across spots.

**a**: small spots ($N = 10$ cells per spot) with low cell-type complexity (up to $M = 3$ cell types per spot from $K = 10$ cell types). **b**: large spots ($N = 20$) with high cell-type complexity ($M = 5$ and $K = 10$). **c**: $N = 10, M = 3$ and $K = 5$.

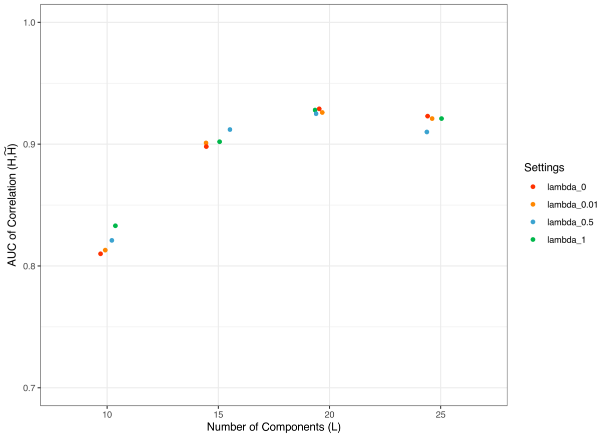


a. $N = 10, M = 3, K = 10$

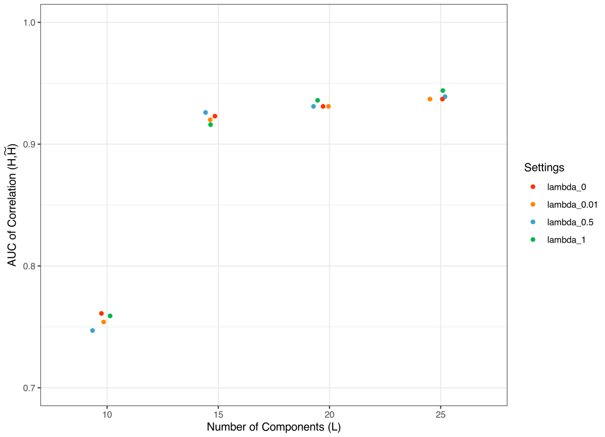


b. $N = 20, M = 5, K = 10$

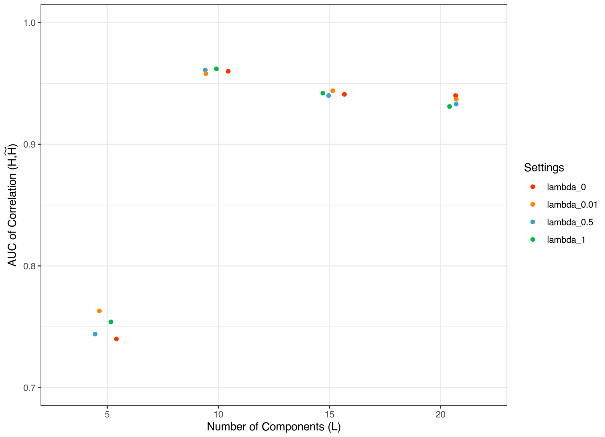


c. $N = 10, M = 3, K = 5$

#### Supplementary Figure 17

Evaluation of RETROFIT's robustness to prior hyperparameter initialization on synthetic ST datasets with different spot size, cell-type complexity and reference quality. The datasets used in this study the simulated datasets used in **Figure 2**, i.e., Column 1: small spots ($N = 10$ cells per spot) with low cell-type complexity (up to $M = 3$ cell types per spot from $K = 10$ cell types). Column 2: large spots ($N = 20$) with high cell-type complexity ($M = 5$ and $K = 10$). Column 3: $N = 10, M = 3$ and $K = 5$. For columns 1-2, exact reference of all 10 ground-truth cell types was provided while for column 3, the reference consisted of all 5 ground-truth plus 5 irrelevant cell types. Each dataset contains $n = 1000$ spots per simulation.

RETROFIT is initialized with the following five different specifications of $\{\alpha_0^W, \beta_0^W, \alpha_0^\theta, \beta_0^\theta, \alpha_0^H, \beta_0^H\}$ while keeping all other parameters the same (random seed=1; $\lambda = 0.01$; $I = 4000$ iterations):

| **Initialization** | **#1** | **#2** | **#3** | **#4** | **#5 (Default)** |
|---|---|---|---|---|---|
| $\alpha_0^W$ | 0.05 | 0.1 | 0.05 | 0.01 | 0.05 |
| $\beta_0^W$ | 0.0001 | 0.0005 | 0.0001 | 0.0005 | 0.0001 |
| $\alpha_0^\theta$ | 1 | 5 | 5 | 1 | 1.25 |
| $\beta_0^\theta$ | 10 | 20 | 20 | 10 | 10 |
| $\alpha_0^H$ | 1 | 0.5 | 1 | 1 | 0.2 |
| $\beta_0^H$ | 1 | 0.5 | 1 | 1 | 0.2 |

In the above table, Initialization #5 corresponds to the default setting of the RETROFIT software, which has been used throughout this study.

The evaluation is based on the following metrics: Row 1: Distribution of RMSE. Row 2: ranked correlation between true ($\mathbf{H}$) and estimated cell-type proportions ($\widetilde{\mathbf{H}}$) across all cell types at each spot. Row 3: Distribution of NRMSE, Row 4: ranked correlation between observed ($\mathbf{X}$) and reconstructed expression ($\widetilde{\mathbf{X}}$) across all genes at each spot. The one-sided KS $P$-values are shown in rows 1 and 3 (black: $P < 0.05$; brown: $P > 0.05$). A small P-value indicates a statistically significant difference in RMSE distributions compared to another parameter setting. The area under the curve (AUC) of ranked correlation curves across spots is shown for each method with a matching color in rows 2 and 4.

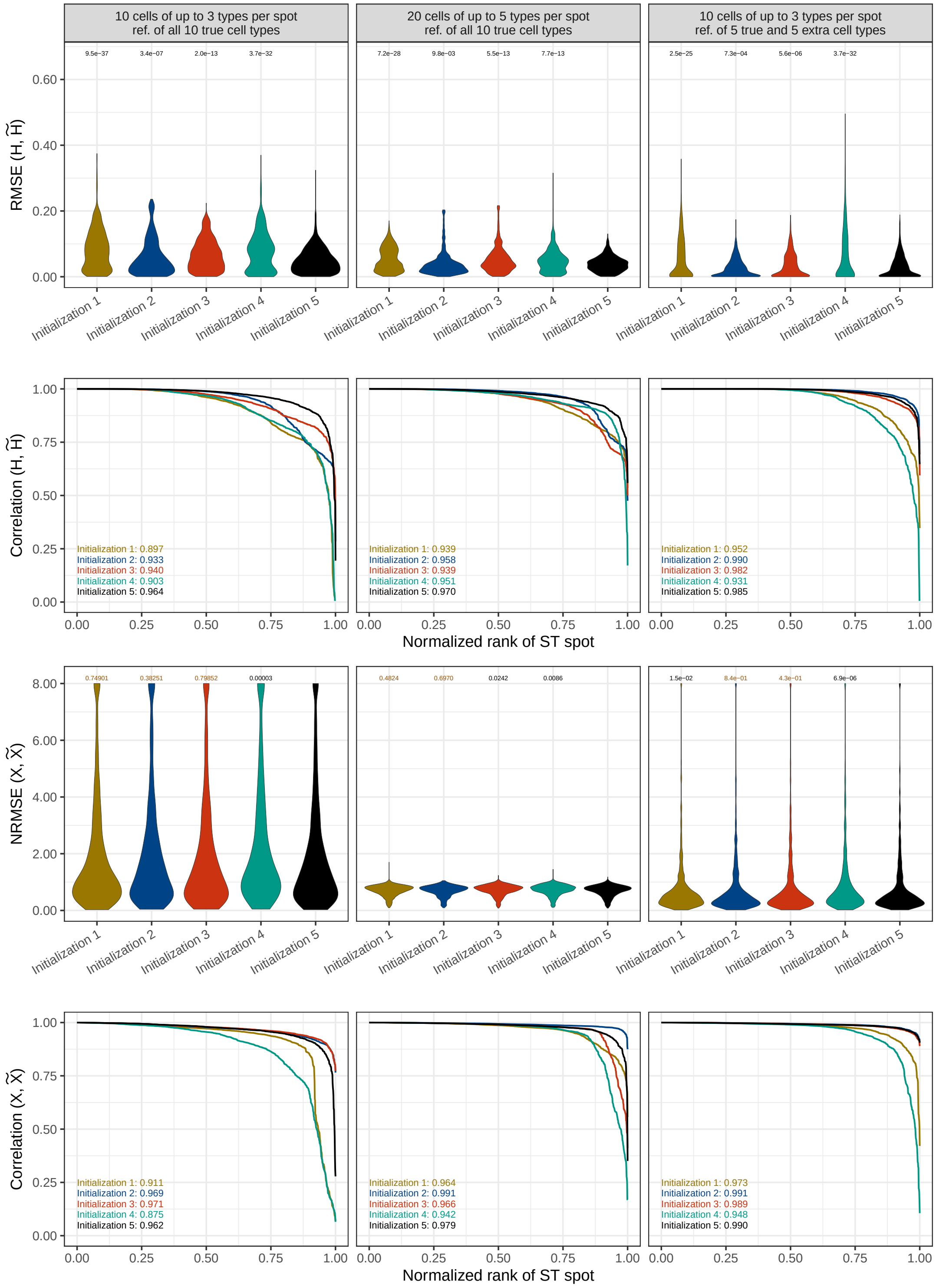
10 cells of up to 3 types per spot
ref. of all 10 true cell types
20 cells of up to 5 types per spot
ref. of all 10 true cell types
10 cells of up to 3 types per spot
ref. of 5 true and 5 extra cell types
RMSE (H, H̃)
9.5e−37
3.4e−07
2.0e−13
3.7e−32
7.2e−28
9.8e−03
5.5e−13
7.7e−13
2.5e−25
7.3e−04
5.6e−06
3.7e−32
Initialization 1
Initialization 2
Initialization 3
Initialization 4
Initialization 5
Correlation (H, H̃)
Initialization 1: 0.897
Initialization 2: 0.933
Initialization 3: 0.940
Initialization 4: 0.903
Initialization 5: 0.964
Initialization 1: 0.939
Initialization 2: 0.958
Initialization 3: 0.939
Initialization 4: 0.951
Initialization 5: 0.970
Initialization 1: 0.952
Initialization 2: 0.990
Initialization 3: 0.982
Initialization 4: 0.931
Initialization 5: 0.985
Normalized rank of ST spot
NRMSE (X, X̃)
0.74901
0.38251
0.79852
0.00003
0.4824
0.6970
0.0242
0.0086
1.5e−02
8.4e−01
4.3e−01
6.9e−06
Correlation (X, X̃)
Initialization 1: 0.911
Initialization 2: 0.969
Initialization 3: 0.971
Initialization 4: 0.875
Initialization 5: 0.962
Initialization 1: 0.964
Initialization 2: 0.991
Initialization 3: 0.966
Initialization 4: 0.942
Initialization 5: 0.979
Initialization 1: 0.973
Initialization 2: 0.991
Initialization 3: 0.989
Initialization 4: 0.948
Initialization 5: 0.990
Normalized rank of ST spot

#### Supplementary Figure 18

Comparison of annotation algorithms based on three synthetic ST datasets with different spot size, cell-type complexity and reference quality. The datasets used in this study the simulated datasets used in **Figure 2**, i.e., Column 1: small spots ($N = 10$ cells per spot) with low cell-type complexity (up to $M = 3$ cell types per spot from $K = 10$ cell types). Column 2: large spots ($N = 20$) with high cell-type complexity ($M = 5$ and $K = 10$). Columns 3-4: $N = 10, M = 3$ and $K = 5$. Columns 1-2 received exact reference of all 10 ground-truth cell types while Column 3 received all 5 ground-truth plus 5 irrelevant cell types. Each dataset contains $n = 1000$ spots per simulation.

Top: Distribution of RMSE with one-sided KS $P$-values (black: $P < 0.05$; brown: $P > 0.05$). A small P-value indicates a statistically significant difference in RMSE distributions compared to another annotation algorithm.

Bottom: Ranked correlation between true ($\mathbf{H}$) and estimated cell-type proportions ($\widetilde{\mathbf{H}}$) across all cell types at each spot. The area under the curve (AUC) of ranked correlation curves across spots is shown for each annotation algorithm with a matching color.

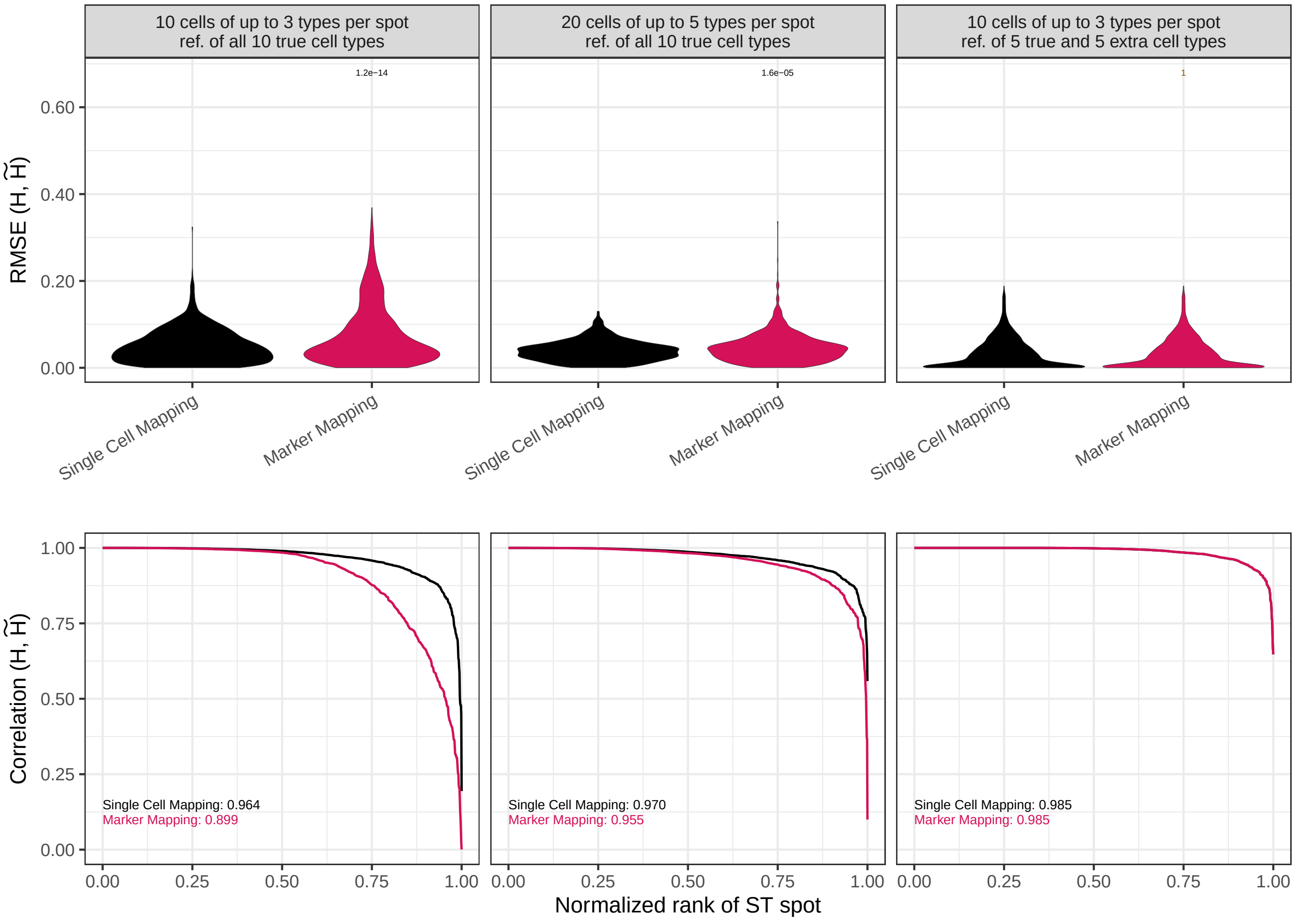

#### Supplemental Figure 19

RETROFIT analysis of the mouse main olfactory bulb (MOB) dataset from the STdeconvolve study. a. Spatial distribution of RETROFIT component X2 (black shading) overlaid on the cell clusters defined by STdeconvolve. b. Spatial expression patterns of the three RMS-associated genes (Nrep, Sox11, and Dcx), known markers of neuronal precursor cells in the rostral migratory stream (RMS). c. Corresponding in situ hybridization (ISH) images for Nrep and Sox11 from the Allen Brain Atlas[8].

a.

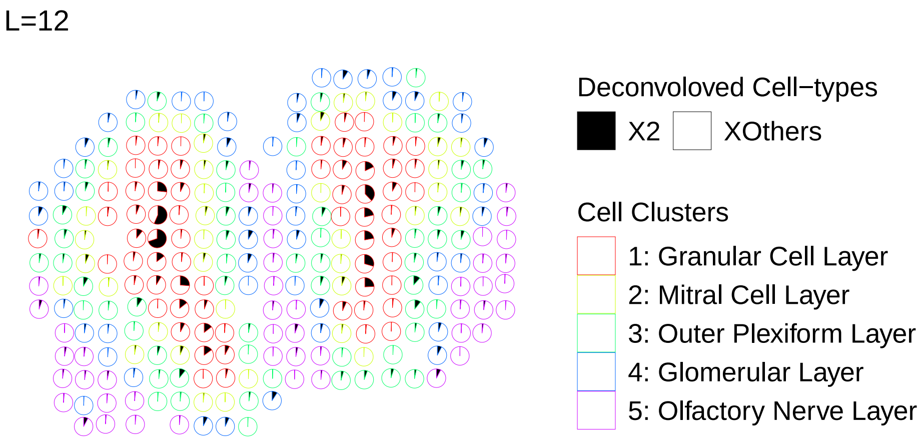


b.

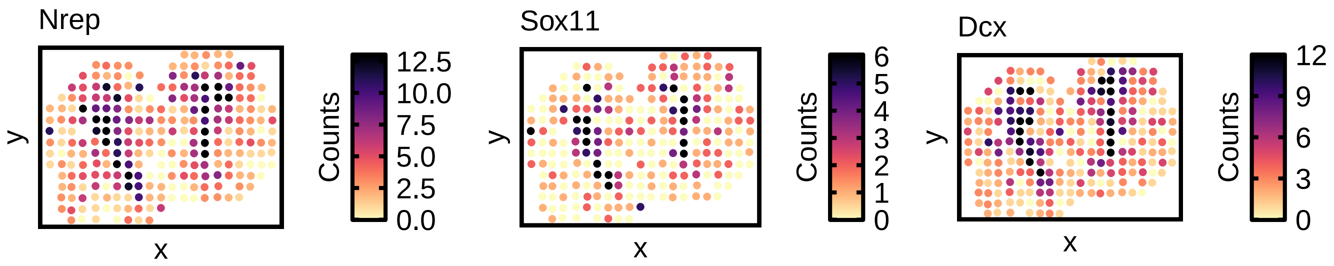


c.

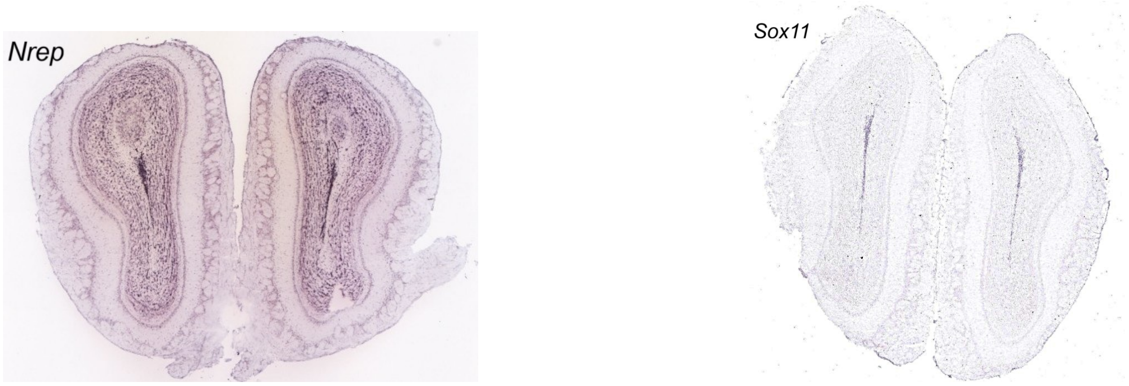

#### Supplemental Figure 20

Highest achievable correlation of STdeconvolve on Visium HD data. For each cell type, the STdeconvolve component with the highest Pearson correlation between spatial marker gene expression and estimated cell-type proportions is shown. Pearson correlation values (R) are reported for each matched component. This analysis represents the best case performance of STdeconvolve when component selection is guided by marker gene information itself.

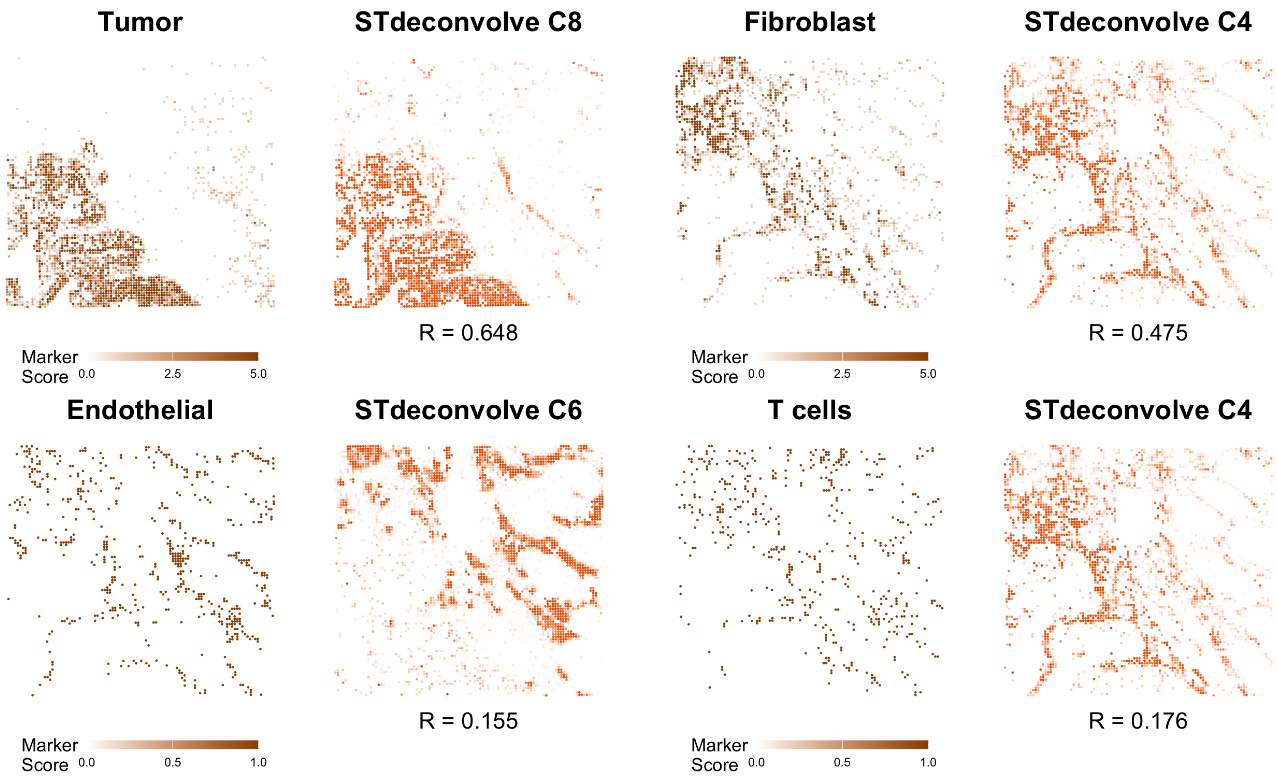

#### Supplementary Table Legends

All the supplementary tables listed below are included in a stand-alone Excel Workbook (xlsx) file.

**Table S1** Computational complexity of RETROFIT on 7 ST datasets. The size of each ST dataset is determined by the numbers of genes and spots. The computational complexity of RETROFIT on each ST dataset is captured by the average computational time per iteration and the total memory usage. For each of the 7 ST datasets, the computational time and memory usage were measured on a Mac mini 2018 model with R version 4.3.0.

**Table S2** Cell-type marker genes curated for the mouse brain. This table consists of 15 granule, 4 oligodendrocyte and 42 Purkinje marker genes. These genes were retrieved from www.neuroexpresso.org.

**Table S3** ST expression scores and proportion estimates of granule cells across all spots in the mouse cerebellum Slide-seq dataset. Each row indicates a spot in the ST slide. This table corresponds to the first row of Fig. 3.

**Table S4** ST expression scores and proportion estimates of oligodendrocytes across all spots in the mouse cerebellum Slide-seq dataset. Each row indicates a spot in the ST slide. This table corresponds to the second row of Fig. 3.

**Table S5** ST expression scores and proportion estimates of Purkinje cells across all spots in the mouse cerebellum Slide-seq dataset. Each row indicates a spot in the ST slide. This table corresponds to the third row of Fig. 3.

**Table S6** Proportions of 8 cellular compartments estimated by RETROFIT ($\widetilde{\mathbf{H}}$) across all spots in the human 12 PCW intestine Visium dataset. RETROFIT extracted $L = 16$ latent components from this ST slide (Algorithm 1) and mapped them to $K = 8$ cellular compartments on the basis of 37 compartment-specific marker genes (Algorithm 3). Each row indicates a spot in the ST slide. This table corresponds to the 12 PCW results shown in Fig.s 5-6 and Supplementary Fig.s 5-11.

**Table S7** Proportions of 8 cellular compartments estimated by RETROFIT ($\widetilde{\mathbf{H}}$) across all spots in the human 19 PCW intestine Visium dataset. RETROFIT extracted $L = 16$ latent components from this ST slide (Algorithm 1) and mapped them to $K = 8$ cellular compartments on the basis of 37 compartment-specific marker genes (Algorithm 3). Each row indicates a spot in the ST slide. This table corresponds to the 19 PCW results shown in Fig.s 5-6 and Supplementary Fig.s 5-11.

**Table S8** Proportions of 8 cellular compartments estimated by RETROFIT ($\widetilde{\mathbf{H}}$) across all spots in the human adult intestine Visium dataset. RETROFIT extracted $L = 16$ latent components from this ST slide (Algorithm 1) and mapped them to $K = 8$ cellular compartments on the basis of 37 compartment-specific marker genes (Algorithm 3). Each row indicates a spot in the ST slide. This table corresponds to the adult results shown in Fig.s 5-6 and Supplementary Fig.s 5-11.

**Table S9** Proportions of 8 cellular compartments estimated by RETROFIT ($\widetilde{\mathbf{H}}$) across all spots in the human 12 PCW intestine Visium dataset. RETROFIT extracted $L = 16$ latent components from this ST slide (Algorithm 1) and mapped them to $K = 8$ cellular compartments on the basis of a companion human 12 PCW intestine scRNA-seq dataset (Algorithm 2). Each row indicates a spot in the ST slide.

**Table S10** Proportions of 8 cellular compartments estimated by RETROFIT ($\widetilde{\mathbf{H}}$) across all spots in the human 19 PCW intestine Visium dataset. RETROFIT extracted $L = 16$ latent components from this ST slide (Algorithm 1) and mapped them to $K = 8$ cellular compartments on the basis of a companion human 19 PCW intestine scRNA-seq dataset (Algorithm 2). Each row indicates a spot in the ST slide.

**Table S11** Pearson correlations between RETROFIT estimated proportions (Tables S9-10) and ST expression scores of known marker genes across spots in fetal and adult slides for all cellular compartments when annotated by marker genes or scRNA-seq. This table corresponds to the adult results shown in Fig. 5.

**Table S12** Performance comparison of annotation strategies using the evaluation metric $\delta$ (Supplementary Note 3 Algorithm 1) in three ST datasets.

**Table S13** Pearson correlations between estimated proportions and ST expression scores of known marker genes across spots in both fetal slides for all cellular compartments when annotated by marker genes or scRNA-seq. Proportions are estimated by CARDfree (Tables S14-15), Cell2location (Tables S16-17), and STdeconvolve (Tables S18-19).

**Table S14** Proportions of 8 cellular compartments estimated by CARDfree across all spots in the human 12 PCW intestine Visium dataset. Each row indicates a spot in the ST slide.

**Table S15** Proportions of 8 cellular compartments estimated by CARDfree across all spots in the human 19 PCW intestine Visium dataset. Each row indicates a spot in the ST slide.

**Table S16** Proportions of 8 cellular compartments estimated by Cell2location across all spots in the human 12 PCW intestine Visium dataset. Each column indicates a spot in the ST slide.

**Table S17** Proportions of 8 cellular compartments estimated by Cell2location across all spots in the human 12 PCW intestine Visium dataset. Each column indicates a spot in the ST slide.

**Table S18** Proportions of 16 cellular compartments estimated by STDeconvolve across all spots in the human 12 PCW intestine Visium dataset. Each row indicates a spot in the ST slide.

**Table S19** Proportions of 16 cellular compartments estimated by STDeconvolve across all spots in the human 19 PCW intestine Visium dataset. Each row indicates a spot in the ST slide.

**Table S20** Compartment-specific expression levels estimated by RETROFIT ($\tilde{\mathbf{W}}$) across all genes in the human 12 PCW intestine Visium dataset. RETROFIT extracted $L = 16$ latent components from this ST slide (Algorithm 1) and mapped them to $K = 8$ cellular compartments on the basis of 37 compartment-specific marker genes (Algorithm 3). Each row indicates a gene whose ST expression levels were analyzed by RETROFIT. This table corresponds to the 12 PCW results shown in Fig. 7.

**Table S21** Compartment-specific expression levels estimated by RETROFIT ($\tilde{\mathbf{W}}$) across all genes in the human 19 PCW intestine Visium dataset. RETROFIT extracted $L = 16$ latent components from this ST slide (Algorithm 1) and mapped them to $K = 8$ cellular compartments on the basis of 37 compartment-specific marker genes (Algorithm 3). Each row indicates a gene whose ST expression levels were analyzed by RETROFIT. This table corresponds to the 19 PCW results shown in Fig. 7.

**Table S22** Compartment-specific expression levels estimated by RETROFIT ($\tilde{\mathbf{W}}$) across all genes in the human adult PCW intestine Visium dataset. RETROFIT extracted $L = 16$ latent components from this ST slide (Algorithm 1) and mapped them to $K = 8$ cellular compartments on the basis of 37 compartment-specific marker genes (Algorithm 3). Each row indicates a gene whose ST expression levels were analyzed by RETROFIT. This table was used to determine the compartment-specific genes shown in the adult panel of Fig. 7b.

**Table S23** Compartment-specific expression levels estimated by RETROFIT ($\tilde{\mathbf{W}}$) across all genes in the Visium dataset of a replicate human 12 PCW intestine sample (sample ID: D1). RETROFIT extracted $L = 16$ latent components from this ST slide (Algorithm 1) and mapped them to $K = 8$ cellular compartments on the basis of 37 compartment-specific marker genes (Algorithm 3). Each row indicates a gene whose ST expression levels were analyzed by RETROFIT. This table was used to determine the compartment-specific genes shown in the 12 PCW panel of Fig. 7b.

**Table S24** Compartment-specific expression levels estimated by RETROFIT ($\tilde{\mathbf{W}}$) across all genes in the Visium dataset of a replicate human 12 PCW intestine sample (sample ID: D2). RETROFIT extracted $L = 16$ latent components from this ST slide (Algorithm 1) and mapped them to $K = 8$ cellular compartments on the basis of 37 compartment-specific marker genes (Algorithm 3). Each row indicates a gene whose ST expression levels were analyzed by RETROFIT. This table was used to determine the compartment-specific genes shown in the 12 PCW panel of Fig. 7b.

**Table S25** Compartment-specific expression levels estimated by RETROFIT ($\widetilde{\mathbf{W}}$) across all genes in the Visium dataset of a replicate human adult intestine sample (sample ID: A2). RETROFIT extracted $L = 16$ latent components from this ST slide (Algorithm 1) and mapped them to $K = 8$ cellular compartments on the basis of 37 compartment-specific marker genes (Algorithm 3). Each row indicates a gene whose ST expression levels were analyzed by RETROFIT. This table was used to determine the compartment-specific genes shown in the adult panel of Fig. 7b.

**Table S26** Biological pathway enrichments of compartment-specific genes identified by RETROFIT in the human fetal (12 PCW) and adult intestine Visium datasets. These results were generated at http://metascape.org. Each row indicates a pathway with statistically significant enrichment after multiple-testing correction (FDR < 0.05). This table was used to determine the top-ranked biological pathways enriched in muscle-specific genes shown in Fig. 7b.

**Table S27** Cell-type signature gene set enrichments of compartment-specific genes identified by RETROFIT in the human fetal (12 PCW) and adult intestine Visium datasets. These results were generated at http://metascape.org. Each row indicates a gene set with statistically significant enrichment after multiple-testing correction (FDR < 0.05).

**Table S28** Values of the diagnostic statistic ($\delta$) across different $\lambda$ settings for the mouse cerebellum Slide-seq and human intestine Visium datasets. The highest $\delta$ for each dataset is shown in bold.

**Table S29** Computation time and memory consumption for each method applied to the mouse cerebellum Slide-seq dataset.

**Table S30** Estimated cell-type proportions of tumor, T cells, endothelial and fibroblast cells across all spots in the Visium HD colorectal cancer dataset. Each row indicates a spot in the ST slide. Different column correspond to different methods - RETROFIT, RCTD, STDeconvolve and Cell2location. This table corresponds to the Fig. 4.
